# Phase separating RNA binding proteins form heterogeneous distributions of clusters in subsaturated solutions

**DOI:** 10.1101/2022.02.03.478969

**Authors:** Mrityunjoy Kar, Furqan Dar, Timothy J. Welsh, Laura Vogel, Ralf Kühnemuth, Anupa Majumdar, Georg Krainer, Titus M. Franzmann, Simon Alberti, Claus A. M. Seidel, Tuomas P.J. Knowles, Anthony A. Hyman, Rohit V. Pappu

## Abstract

Macromolecular phase separation is thought to be one of the processes that drives the formation of membraneless biomolecular condensates in cells. The dynamics of phase separation, especially at low endogenous concentrations found in cells, are thought to follow the tenets of classical nucleation theory describing a sharp transition between a dense phase and a dilute phase characterized by dispersed monomers. Here, we used *in vitro* biophysical studies to study subsaturated solutions of phase separating RNA binding proteins with intrinsically disordered prion like domains (PLDs) and RNA binding domains (RBDs). Surprisingly, we find that subsaturated solutions are characterized by heterogeneous distributions of clusters comprising tens to hundreds of molecules. These clusters also include low abundance mesoscale species that are several hundreds of nanometers in diameter. Our results show that cluster formation in subsaturated solutions and phase separation in supersaturated solutions are strongly coupled via sequence-encoded interactions. Interestingly, however, cluster formation and phase separation can be decoupled from one another using solutes that impact the solubilities of phase separating proteins. They can also be decoupled by specific types of mutations. Overall, our findings implicate the presence of distinct, sequence-specific energy scales that contribute to the overall phase behaviors of RNA binding proteins. We discuss our findings in the context of theories of associative polymers.

**Significance Statement:** Membraneless biomolecular condensates are molecular communities with distinct compositional preferences and functions. Considerable attention has focused on phase separation as the process that gives rise to condensates. Here, we show that subsaturated solutions of RNA binding proteins form heterogeneous distributions of clusters in subsaturated solutions. The formation of clusters in subsaturated solutions and condensates in supersaturated solution are coupled through sequence-specific interactions. Given the low endogenous concentrations of phase separating proteins, our findings suggest that clusters in subsaturated conditions might be of functional relevance in cells.

## Introduction

Phase separation of RNA binding proteins with disordered prion-like domains (PLDs) and RNA binding domains (RBDs) is implicated in the formation and dissolution of membraneless biomolecular condensates such as RNA-protein (RNP) granules (1–9). Phase separation is a process whereby a macromolecule in a solvent separates into a dilute, macromolecule-deficient phase that coexists with a dense, macromolecule-rich phase (10, 11). In a binary mixture, the soluble phase, comprising dispersed macromolecules that are well-mixed with the solvent, becomes saturated at a concentration designated as *c*_sat_. Above *c*_sat_, for bulk concentrations *c*_tot_ that are between the binodal and spinodal, phase separation of full-length RNA binding proteins and PLDs is thought to follow classical nucleation theory (12–15).

In theories of classical nucleation, clusters representing incipient forms of the new dense phase form within dispersed phases of supersaturated solutions defined by *c*_tot_ > *c*_sat_ (16, 17). In the simplest formulation of classical nucleation theory (16–18), the free energy of forming a cluster of radius *a* is: 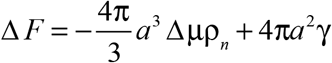. Here, μμ is the difference in the chemical potential between the dense and dilute phases, which is negative in supersaturated solutions and positive in subsaturated solutions; ρ_*n*_ is the number of molecules per unit volume and *σ* is the interfacial tension between dense and dilute phases. At temperature *T*, in a seed-free solution, the degree of supersaturation *s* is defined as: 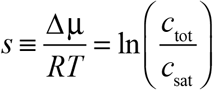 where *R* is the ideal gas constant. Here, *s* is positive for *c*_tot_ > *c*_sat_, and as *s* increases, cluster formation becomes more favorable. Above a critical radius *a*^*^, the free energy of cluster formation overcomes the interfacial penalty, and the new dense phase grows in a thermodynamically downhill fashion. Recently, these ideas from classical nucleation theory were applied to analyze and interpret the dynamics of phase separation in supersaturated solutions in cells (12, 13, 15).

Classical nucleation theories stand in contrast to two-step nucleation theories that predict the existence of pre-nucleation clusters in supersaturated solutions (19, 20). These newer theories and recent measurements (14) hint at the prospect of interesting features in subsaturated solutions. To condition our expectations, we return to classical nucleation theory and note that when applied to polymer solutions (10, 11), there is only one effective energy scale as captured by the Flory parameter χ (10). This determines solvent quality and is quantified by the effective strengths of interactions between pairs of polymer segments in solution and hence χ (21). Phase separation is thermodynamically favored if χ is positive, and this implies a negative Δμ (21). Accordingly, for a given polymer-solvent mixture, defined by a positive value of χ, the magnitude of χ determines both *c*_sat_ and g. However, *s* is negative in subsaturated solutions, and hence cluster formation should be a purely uphill process for *c*_tot_ < *c*_sat_. This is true even if we set g to be zero in subsaturated solutions. As a result, classical nucleation theory, applied without any modifications, predicts that the probability of forming clusters comprising more than 3-5 molecules will be essentially zero in subsaturated solutions (Fig. 1). This is true irrespective of the degree of subsaturation and is independent of details of the system of interest.

**Fig. 1:**
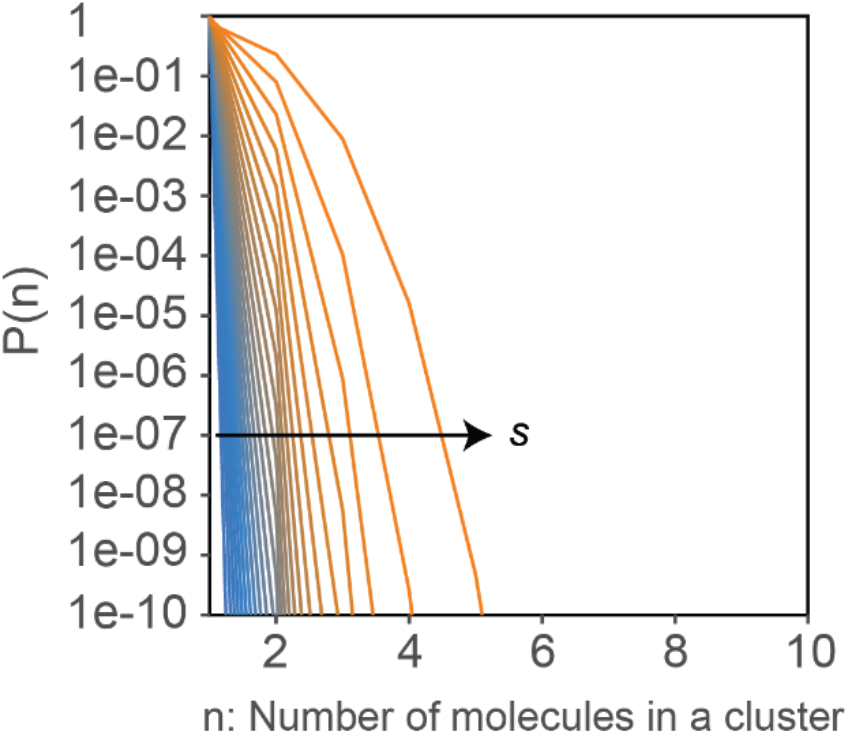
Predictions from classical nucleation theory show that clusters larger than 3-5 molecules do not form in subsaturated solutions. Here, we set g to be zero to illustrate the fact that cluster formation is thermodynamically suppressed in subsaturated solutions even without considerations of an interfacial penalty. Values of *s* were drawn from the interval −3.2 ≤ *s* ≤ −0.04. This range of *s* values matches the range probed in the current study.

To date, there have not been systematic analyses of the size distribution of protein clusters below the saturation concentration for phase separation. Here, we report results from measurements of cluster size distributions in subsaturated solutions of phase-separating RNA binding proteins from the FUS-EWSR1-TAF15 (FET) family. Surprisingly, we find that these systems form heterogeneous distributions of clusters in subsaturated solutions. While the abundant species are small clusters, the distributions of cluster sizes appear to be heavy tailed. Further, as bulk concentrations increase, the distributions of cluster sizes shift continuously toward larger values. We discuss these findings in the context of theories for associative polymers (9, 22–29).

## Results

### Macroscopic phase separation is not observed in subsaturated solutions

Following Wang et al., (9) we quantified *c*_sat_ using a spin-down based absorbance assay. At low concentrations, any prior aggregates that are present in solution are removed upon centrifugation at 20,000 relative centrifugal force (RCF) units. Results from this assay are summarized in Fig. 2A for FUS-SNAP. At low concentrations, the concentration in the supernatant increases monotonically with the bulk concentration of SNAP-tagged FUS. However, above a threshold concentration, which we designate as *c*_sat_, the concentration in the supernatant remains fixed at a plateau value. This is consistent with the establishment of phase equilibrium between dilute and dense phases (Fig. 2A).

**Fig. 2:**
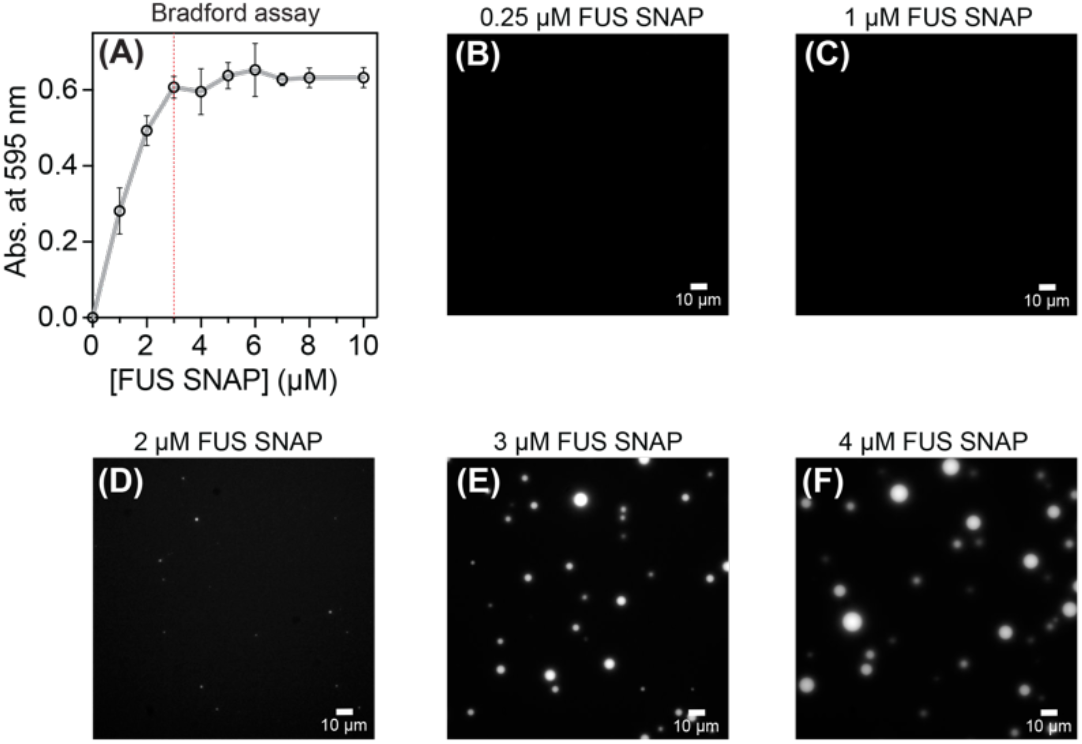
FUS-SNAP has a quantifiable *c*_sat_ whereby condensates form only in supersaturated solutions. (A) Sample data for absorbance-based spin-down assays. Data shown here are for FUSSNAP in 20 mM Tris, pH 7.4, and 100 mM KCl at ≈ 25°C. The red dashed line intersects the abscissa at 3 μM, which is the inferred *c*_sat_ for this construct. Panels (B)-(F) show microscopy images collected at the 18-hr time point for solutions containing different concentrations of FUSSNAP in 20 mM Tris, pH 7.4, and 100 mM KCl at ≈ 25°C. For imaging purposes, 5% of the total mixture in each sample is made up of FUS-eGFP. The scale bar in each panel corresponds to 10 μm. The data show the presence of condensates at or above *c*_sat_ of ≈ 3 μM. The FUS constructs used were expressed and purified using method A (see *SI Appendix*.)

To test the veracity of the estimated *c*_sat_ as a true saturation concentration, we performed microscopy-based measurements for solutions containing different amounts of FUS-SNAP and untagged FUS (*SI Appendix*, Fig. S1). The concentrations investigated range from 0.25 μM to 4 μM. Data were collected approximately 30 minutes after sample preparation and for each sample, a series of images were collected at different time points over an 18-hour period. Results at the 18-hour time point are shown in Figs. 2B-F for different concentrations of FUS-SNAP. Irrespective of the time point of interrogation, condensate formation of FUS-SNAP is only detectable at or above *c*_sat_ ≈ 3 μM. Similar data were obtained for untagged FUS (*SI Appendix*, Fig. S1). Based on these results, we conclude that macroscopic phase separation is not realized in subsaturated solutions and that the FUS molecules are defined by construct- and solution condition-specific saturation concentrations.

### Clusters spanning a range of sizes form in subsaturated solutions of FUS

We used a panel of spectroscopic methods to investigate subsaturated solutions of FUS and other RNA binding proteins. Unless otherwise specified, all measurements were performed in 20 mM Tris, pH 7.4, and 100 mM KCl at ≈ 25°C. First, we used dynamic light scattering (DLS) to characterize subsaturated solutions of untagged FUS (Fig. 3A, *SI Appendix*, Fig. S2) and FUS-SNAP (Figs. 3B, 3C, *SI Appendix*, Fig. S3). The DLS results we obtain are robust to the protocols used to express and purify FUS and FUS-SNAP molecules (*SI Appendix*, Fig. S4).

**Fig. 3:**
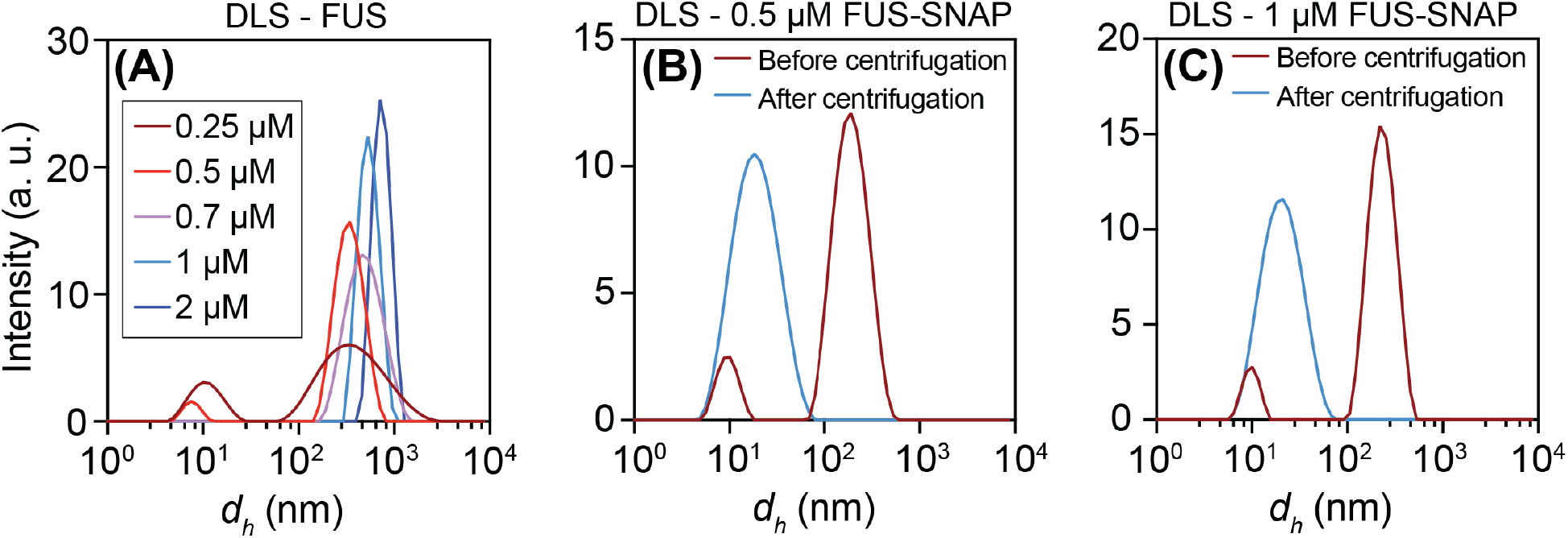
DLS data show that clusters spanning a range of sizes form in subsaturated solutions. **(A)** Measurements were performed at different bulk concentrations of untagged FUS (see legend) that represent different degrees of subsaturation −2 ≤ *s* ≤ 0. Scattering intensities, measured 8 minutes after sample preparation, shift toward larger values as the bulk concentration *c*_tot_ approaches the *c*_sat_ of ≈ 2 μM. Scattering intensity was measured at 0.5 μM (B) and 1 μM (C) of FUS-SNAP at 8-minutes after sample preparation and before centrifugation (brown curves). The solutions were centrifuged at 20,000 RCF for five minutes at room temperature. Then scattering intensities of the supernatant were measured at the 8-minute time point. These data are shown in blue curves in panels (B) and (C). They show the presence of species of sizes ranging between 7 nm and 100 nm. The FUS constructs were expressed and purified using method A (*SI Appendix*).

The DLS data are shown as scattering intensities plotted against the apparent hydrodynamic diameter (*d*_h_). For calibration, the *d*_h_ of untagged FUS monomers is ≈7 nm. The intensity profiles for untagged FUS collected at 0.25 μM show two peaks (Fig. 3A). The peak at ~8 nm corresponds to monomers and oligomers. The second peak at ~200 nm corresponds to mesoscale clusters. The observed bimodality could imply clusters of fixed size forming via either microphase separation (30) or micellization (31). To test for this possibility, we investigated how the DLS signals change with increasing protein concentrations (Fig. 3A). For microphase separation or micellization, we expect the locations of the peaks to stay in place while the intensities of the peaks change respect to one another. We do not observe this behavior. Instead, the location of the peak corresponding to larger *d*_h_ values shifts to the right. At higher concentrations, DLS becomes blind to the presence of smaller species. As a result, the peak at lower values of *d*_h_ vanishes above a concentration of 0.7 μM.

The apparent bimodality seen in intensity profiles can also arise from a heavy tailed distribution of cluster sizes, whereby the most abundant species are monomers and oligomers. In this scenario, bimodality at low concentrations below 0.7 μM can result from smaller species being the most abundant, and the largest, mesoscale species having the highest scattering cross-sections (32). This would mask the presence of species of intermediate sizes. To test for the presence of such species, we subjected subsaturated solutions of FUS-SNAP molecules to centrifugation at 20,000 RCF that removes species larger than 100 nm in diameter. Although, the bulk concentrations prior to centrifugation were 0.5 μM and 1 μM, respectively, the centrifugation step lowers the bulk concentration. The top of the centrifuged solution was then collected for DLS measurements. The DLS data, collected after centrifugation, are shown in Figs. 3B and 3C for FUS-SNAP. The scattering intensity profiles, measured after centrifugation, show the presence of species in the size range of 7 – 100 nm for the apparent *d*_h_. These species include monomers and an assortment of higher-order species. The upper limit on the number of FUS molecules per cluster (*n*_mol_) is estimated as the ratio of the hydrodynamic volume of a cluster of size *d*_h_ to that of the monomer. This yields upper bounds on estimates of *n*_mol_ of 10, 10^2^, and 10^3^ for *d*_h_ values of ~15 nm, ~30 nm, and ~70 nm, respectively.

Taken together with data obtained prior to centrifugation, we conclude that clusters spanning a range of sizes form in solution. The preferred size ranges are governed by the bulk concentration of FUS molecules. The parsimonious interpretation of the data is that the distributions of cluster sizes are likely to be exponentials with heavy tails. This would be concordant with reversible processes such as isodesmic associations that lack a threshold concentration with the average sizes evolve continuously with increasing concentration (33, 34).

### Morphologies of the largest clusters that form in subsaturated solutions

We used negative stain transmission electron microscopy (TEM) to characterize the morphologies of the largest clusters that form in subsaturated solutions. Given their sizes, we refer to these species as mesoscale clusters. These clusters formed by FUS-SNAP have roughly spherical morphologies (Fig. 4A and *SI Appendix*, Fig. S5). This was further confirmed using DLS data collected for untagged FUS at two different angles, Fig. 4B. Based on their quasi-spherical morphologies and plausible densities within clusters, we estimate that mesoscale clusters with diameters of ~150 nm encompass 10^2^-10^3^ molecules.

**Fig. 4:**
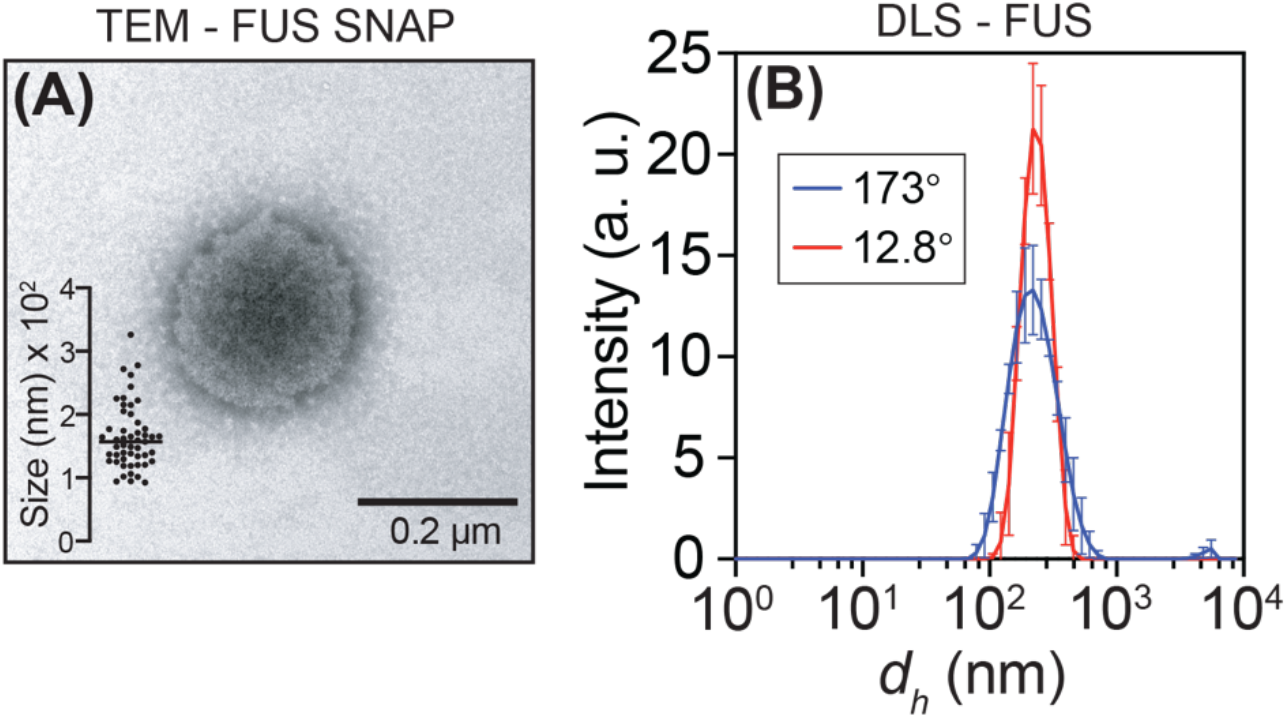
Mesoscale clusters that form in subsaturated solutions have spherical morphologies. Representative TEM image of 2 μM FUS-SNAP. The collections of TEM images were used to quantify the distribution of sizes (see inset and *SI Appendix*, Fig. S6). (B) DLS data collected at two different angles corroborate the spherical morphologies of clusters observed in sub-saturated solutions. Samples used here were prepared using method A (*SI Appendix*).

### Mesoscale clusters are of low overall abundance

The DLS data suggest that the larger, mesoscale clusters (*dh* >100 nm) might be of low abundance. This can be resolved using nanoparticle tracking analysis (NTA) (35). In these measurements, we track the Brownian motion of scatterers using dark field microscopy (Movies S1 and S2). We collected NTA data for untagged FUS (Fig. 5A) and FUS-SNAP (Fig. 5B). The distributions of mesoscale cluster sizes shift toward larger values with increasing protein concentrations. NTA also enables quantification of the percentage of molecules in solution that make up mesoscale clusters (*SI Appendix*, Fig. S6). In a 250 nM solution of FUS-SNAP the bulk volume fraction of proteins is φ ≈ 1.4’ 10^−5^, and the fraction of the solution volume taken up by mesoscale clusters is ϕ_meso_ ≈ 4’ 10^−8^. This translates to 0.15% of the protein molecules in solution being a part of mesoscale clusters (Fig. 5C). For a concentration just below *c*_sat_, the relative abundance of mesoscale clusters increases to ≈ 1%. Similar results were obtained for untagged FUS (*SI Appendix*, Fig. S6).

**Fig. 5:**
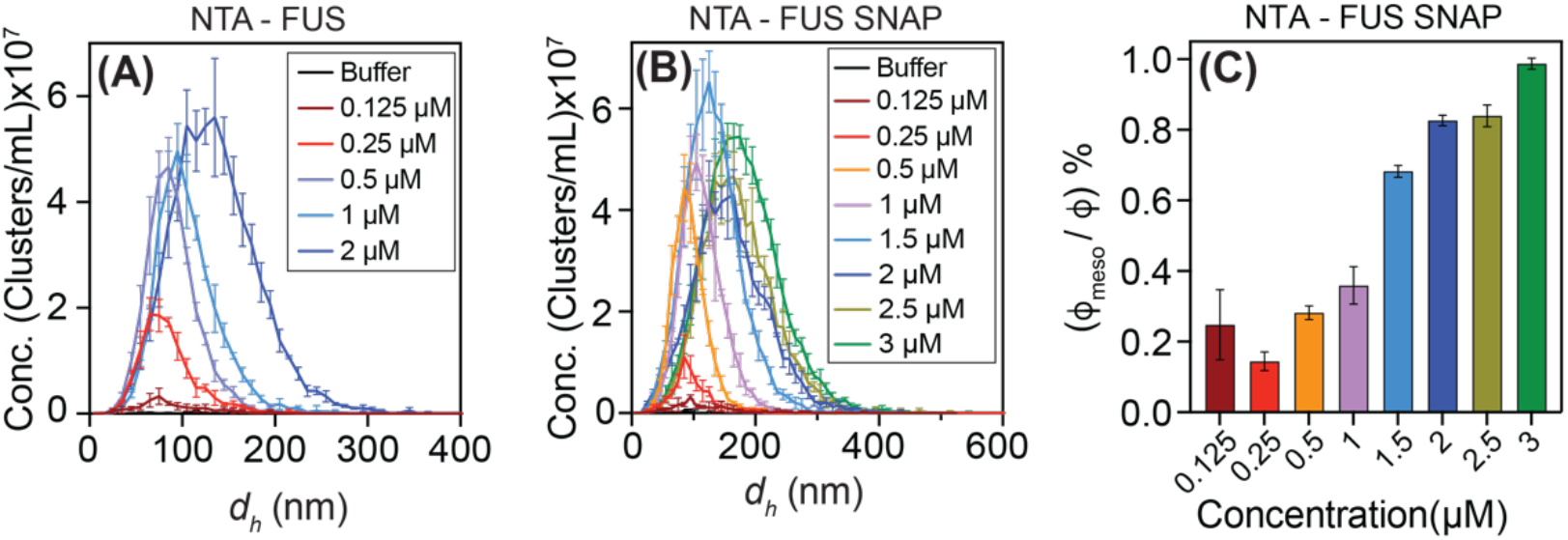
Mesoscale clusters have low overall abundance. (A) NTA data for untagged FUS (*c*_sat_ ≈ 2 μM) were collected at different concentrations that represent different degrees of subsaturation. (B) NTA data for FUS-SNAP (*c*_sat_ ≈ 3 μM) at different degrees of subsaturation. (C) Relative abundance of mesoscale clusters, quantified at different degrees of subsaturation for FUS-SNAP. The constructs used were expressed and purified using method A (*SI Appendix*)

### Clusters increase in size with increasing concentrations and the size distributions are heavy tailed

The data from DLS and NTA measurements show that clusters form in subsaturated solutions and that the average cluster sizes increase monotonically as protein concentrations increase. To probe the distributions across a broader range of sizes, we used orthogonal, fluorescence-based approaches to characterize subsaturated solutions, and access length scales that are distinct from those accessed using DLS and NTA measurements.

Fluorescence anisotropy can be applied for detecting the presence of clusters with different numbers of molecules. These experiments use a small fraction of fluorescently labeled molecules (tracers) and a majority fraction of molecules that lack a fluorescent label. We used confocal multiparameter detection (MFD) which measures the fluorescence anisotropies of diffusing particles (36). This allows us to monitor the size dependent rotational diffusion of molecules. We performed experiments with a fixed concentration (100 nM) of FUS-eGFP and added FUS-SNAP as a titrant with the concentration of FUS-SNAP ranging from 0 to 5 μM. We find that anisotropy increases monotonically at low concentrations of FUS-SNAP (Fig. 6A). This indicates the presence of smaller clusters such as dimers and oligomers with increased volumes that result in longer times for molecular rotational diffusion. The anisotropy of eGFP is especially sensitive to the presence of small clusters. However, this signal saturates for larger clusters. This is because the the apparent volume that is measured has a non-linear scaling due to the inverse relationship between anisotropy and volume. Further, the flexible FUS backbone and long eGFP linker may diminish the actual change in anisotropy. To test for the presence of larger clusters, we therefore used the environmentally sensitive dye Nile Red, which offers a higher dynamic range in anisotropy. With Nile Red, we see a strong continuous increase in anisotropy as the concentration of FUS-SNAP is increased (Fig. 6A). The signal does not saturate as the concentrations of FUSSNAP increase. That this lack of saturation emanates from large bright clusters is readily confirmed in the binned histogram of fluorescence intensity of the photon trace (Fig. 6B). These data show that the size distributions of clusters that form in subsaturated solutions are heavy tailed.

**Fig. 6:**
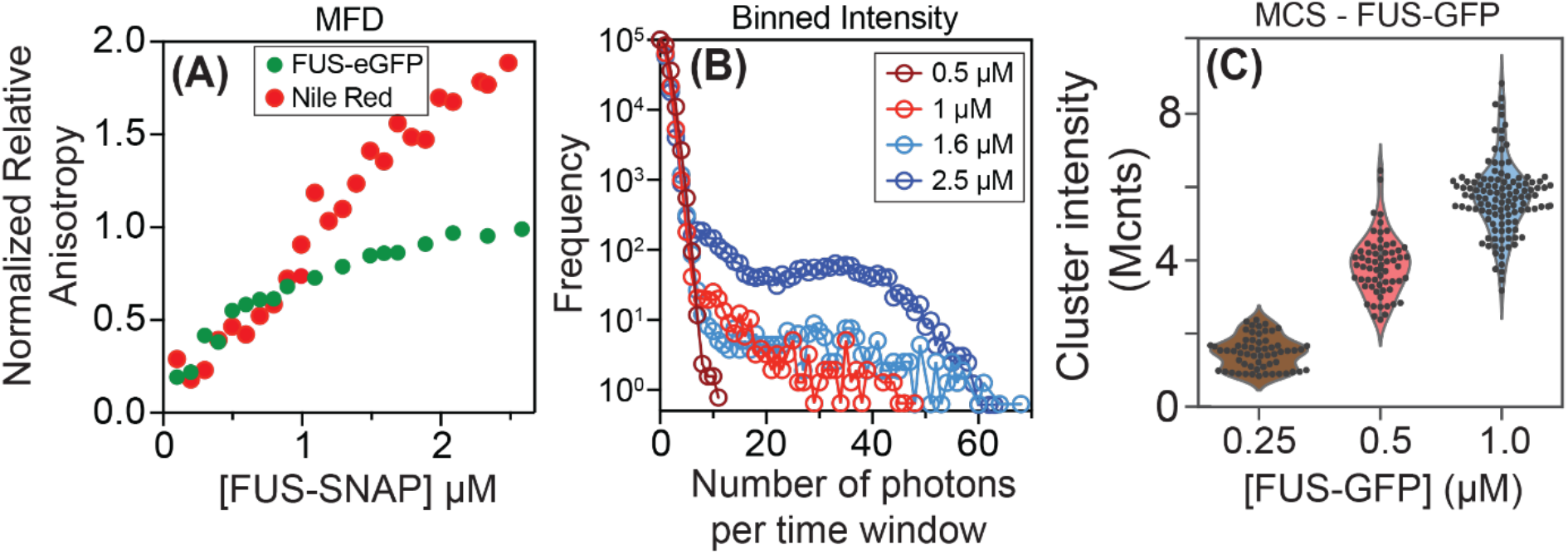
Fluorescence anisotropy and brightness-based measurements show the presence of clusters in subsaturated solutions. (A) Normalized fluorescence anisotropy from confocal multiparameter detection (MFD) plotted against the concentration FUS-SNAP, which is the titrant. Data were collected at 50 mM KCl. The sample was prepared using method B. We also titrated FUS-SNAP from 0 to 3 μM in presence of 20 nM Nile red. Unlike the anisotropy measured using eGFP fluorescence (green), the measurements based on Nile Red (red) do not show saturation behavior, implying a continuous growth of cluster sizes with the concentration of FUS-SNAP. (B) With increasing FUS-SNAP concentrations in the presence of 20 nM Nile red, we observe bright particles, with apparent heavy tailed distributions. (C) MCS data showing the formation of clusters in subsaturated solutions comprising FUS-GFP, prepared using method A. The apparent sizes of clusters, quantified in terms of intensity per molecule, increase with increasing protein concentration. The FUS-GFP constructs used in the MCS experiments were expressed and purified using method A (*SI Appendix*.)

The data in Fig. 6B suggest the presence of species that span a broad spectrum of sizes. As an independent probe for the presence of such species, we turned to confocal detection under flow by using Microfluidic confocal spectroscopy (MCS). This is a brightness-based method that combines microfluidic mixing and flow with confocal detection (37–39). MCS measurements are sensitive to the presence of species that span a broad size range from tens to hundreds of nanometers, and due to convective flow, is not reliant on diffusion alone for sampling species of different sizes in solution (*SI Appendix*, Fig. S7). We used FUS-eGFP (*c*_sat_ ≈ 4 μM, *SI Appendix*, Fig. S8) for the MCS measurements. Data were obtained at concentrations of 0.25 μM, 0.5 μM, and 1 μM, plotted as the brightness per event, clearly show the formation of clusters in subsaturated solutions (Fig. 6B). The average sizes of clusters in subsaturated solutions increase with increasing concentrations.

### Other FET family proteins also form clusters in subsaturated solutions

Next, we asked if the formation of clusters in subsaturated solutions is unique to FUS or if this is a feature that is shared by other members of the FET family of RNA binding proteins. We collected DLS data in subsaturated solutions for untagged hnRNP-A3 (*c*_sat_ ≈ 6 μM, *SI Appendix*, Fig. S9A) (Fig. 7A), EWSR1-SNAP (*c*_sat_ ≈ 2 μM, *SI Appendix*, Fig. S9B) (Fig. 7B), and TAF15-SNAP (*c*_sat_ ≈ 2 μM, *SI Appendix*, Fig. S9C) (Fig. 7C). As with FUS, these data show that each of the proteins forms clusters in subsaturated solutions. The average sizes of these clusters increase with protein concentration. Raw DLS data in the form of autocorrelation functions are shown in *SI Appendix*, Figs. S10-S12.

**Fig. 7:**
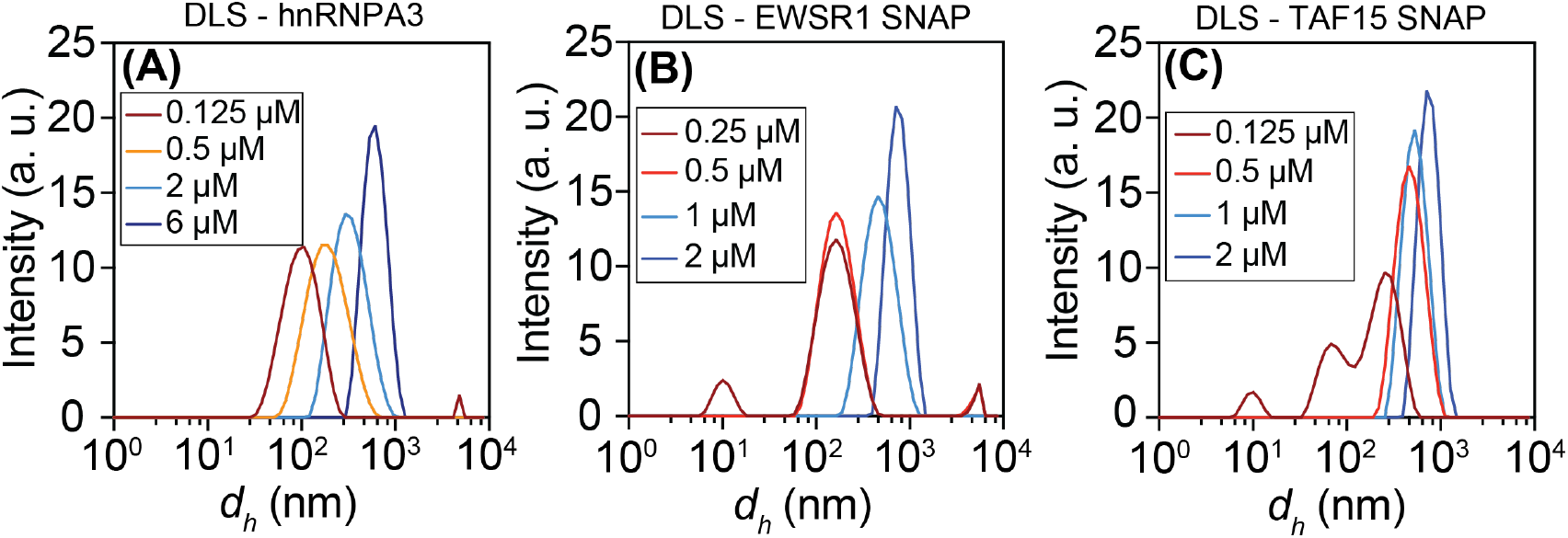
Clusters also form in subsaturated solutions of FET family proteins. DLS data show evidence for mesoscale clusters in subsaturated solutions for (A) hnRNP-A3 (*c*_sat_ ≈ 6 μM), (B) SNAP tagged TAF15 (*c*_sat_ ≈ 2 μM), and (C) SNAP tagged EWSR1 (*c*_sat_ ≈ 2 μM).

### Clusters are reversible and molecules exchange between clusters

Using DLS, we find that the sizes of clusters decrease upon dilution and increase with increased concentration in the subsaturated regime (Fig. 8A-C, *SI Appendix*, Fig. S13). This observation points to the reversibility of cluster formation in subsaturated solutions whereby they shrink upon dilution and grow when concentrations increase. For cluster formation to be reversible, the molecules must exchange between clusters or between clusters and the bulk solution. To test for this, we used Förster resonance energy transfer (FRET) experiments (setup shown in Fig. 8D). These experiments show that FUS molecules readily exchange across clusters (Fig. 8E, *SI Appendix*, Fig. S14).

**Fig. 8:**
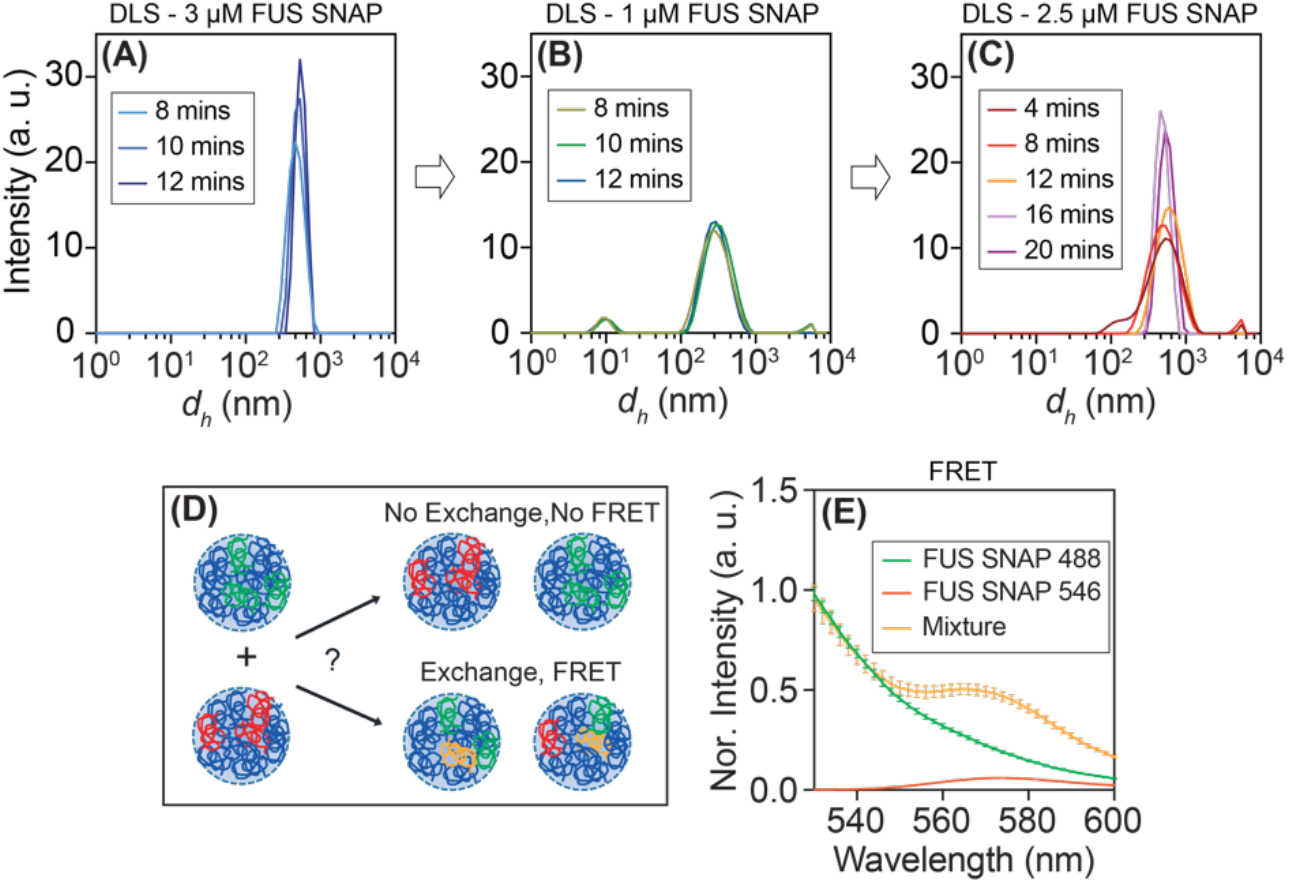
Clusters in subsaturated solution form via reversible associations and molecules readily exchange between clusters. (A) DLS data were collected at different time points for SNAP-tagged FUS at 3 μM. (B) Upon dilution to 1 μM, the DLS data show changes to the intensity profiles, with the appearance of smaller species. (C) Increasing the concentration from 1 to 2.5 μM leads to an increased preference for larger species. (D) Design of the bulk FRET assay. Here, two sets of clusters are formed, each using a total concentration of 1 μM of SNAP-tagged FUS molecules. In each set, 5% of the molecules carry a fluorescent label (Alexa Fluor 488 - green or Alexa Fluor 46 - red). The clusters are mixed to achieve a total concentration of 1 μM. The mixture is excited using a 488 nm laser and the emission spectrum is measured from 520 nm to 600 nm. If molecules exchange between the clusters, then we expect to see a peak at the excitation maximum of 573 nm. (E) Fluorescence emission spectra show the decay of fluorescence for molecules with the Alexa 488 label and for the mixture. The latter shows a maximum at 573 nm, which is indicative of the exchange of molecules between the clusters. The FUS constructs used in this experiment were expressed and purified using method A.

The data presented in Figs. 3–8 show the presence of heterogenous distributions of clusters in subsaturated solutions. Cluster sizes increase as concentrations increase and approach *c*_sat_. The clusters appear to be equilibrium species that form and dissolve via reversible associations. The low abundance of mesoscale clusters quantified using NTA, the presence of smaller species, readily detected using fluorescence anisotropy, and the presence of a broad spectrum of species indicated by MFD, fluorescence intensity analysis, and MCS suggest that the distributions of cluster sizes are concentration-dependent, heterogeneous, and heavy tailed.

### Growth of macroscopic phases above *c*_sat_

Guided by precedents in the literature (40, 41), we asked if there are discernible signatures of dynamical transitions from finite-sized clusters to condensates just above *c*_sat_. Specifically, we used DLS to probe the presence of slow modes (41) in the temporal evolution of autocorrelation functions for concentrations of untagged FUS that are below and above *c*_sat_. Below *c*_sat_, the autocorrelation functions reach a steady-state and do not change with time after a few minutes (Fig. 9A). The timescales interrogated here are roughly seven orders of magnitude longer than the time it takes for individual FUS molecules to diffuse across 10 nm. Just above *c*_sat_ (Fig. 9B), the autocorrelation functions show the presence of slow modes. Such modes have been observed for polymers in dense phases, and have been attributed to reptation (41). However, we interpret this to imply that clusters grow into micron-scale condensates above *c*_sat_. The relevant data are shown for hnRNPA3, EWSR1-SNAP, and TAF15-SNAP (Fig. 9C, *SI Appendix*, Fig S15). That the slow modes in autocorrelation functions point to the onset of condensation processes was independently verified using microscopy. These data show that condensation as a function of time leads to the formation of dense phases (Figs. 9D–9I). The observed temporal evolution is consistent with a coarsening process whereby larger condensates grow at the expense of smaller ones (42, 43). Importantly, our analysis shows that the lowest concentration at which one observes the onset of slow modes in autocorrelation functions measured using DLS can be used as an efficient, centrifugation-free protocol for estimating *c*_sat_.

**Fig. 9:**
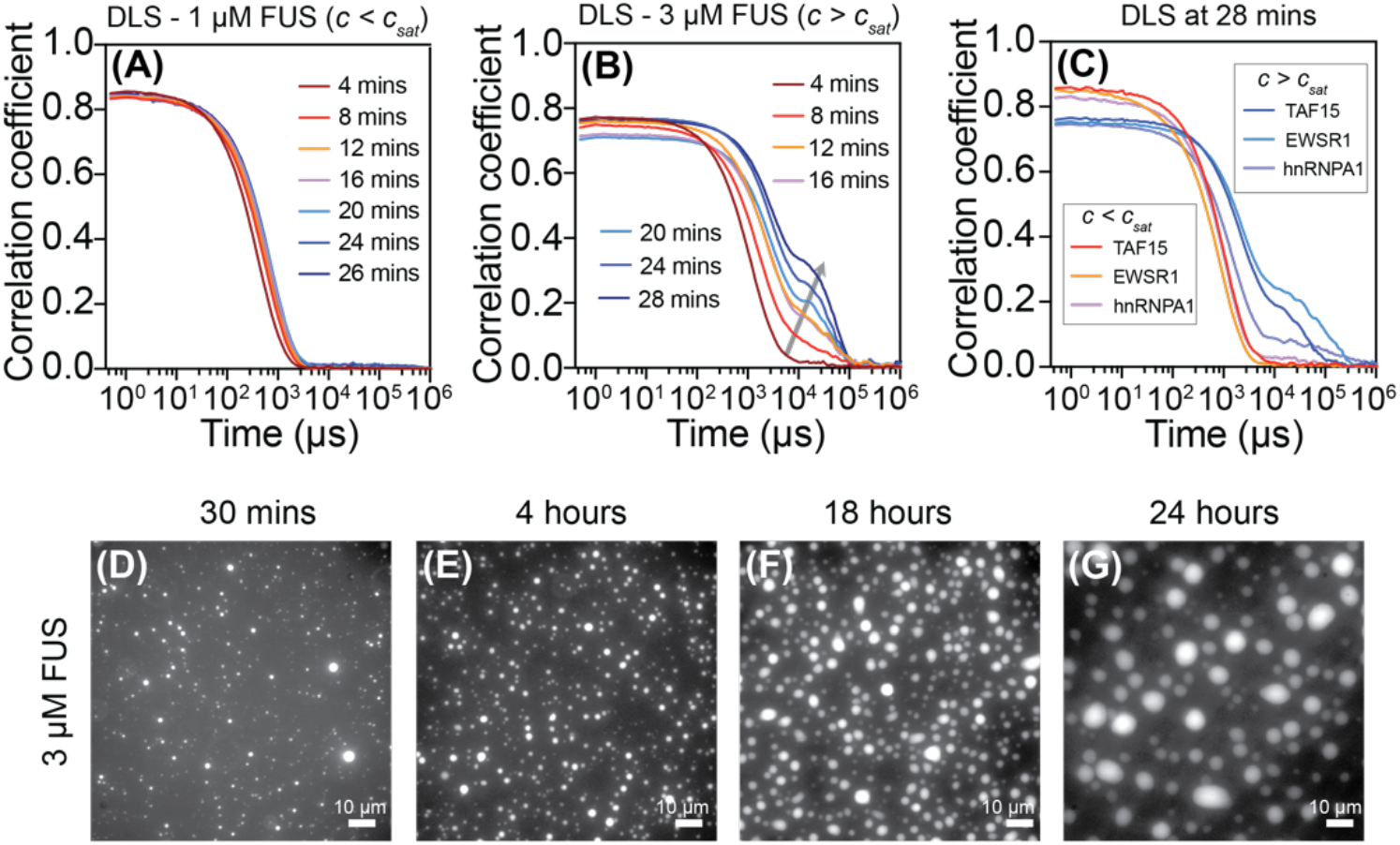
Clusters grow into micron-scale bodies above *c*_sat_. (A) Temporal evolution of autocorrelation functions from DLS measurements for untagged FUS. Below *c*_sat_, the sizes of mesoscale clusters reach a steady state. (B) Above *c*_sat_, we observe increased amplitudes of the autocorrelation function at longer times, (annotated using a gray arrow). (C) DLS data for hnRNPA3, EWSR1-SNAP, and TAF15-SNAP. Below *c*_sat_ the cluster sizes reach a steady state. (D)-(G) Evidence of the growth into micron-scale condensates, displaying coarsening whereby fewer condensates grow by absorbing smaller species. This is made clear by the long-time evolution of micron-scale condensates formed by 3 μM untagged FUS containing 5% FUS-eGFP.

### Cluster formation and macroscopic phase separation can be decoupled

Solutes such as 1,6-hexanediol (HD) can suppress phase separation and dissolve micron-scale condensates (29, 44). We used DLS to measure the impact of increased HD concentration on low abundance mesoscale clusters vs. macroscopic phase separation. These data show a dose-dependent response to HD on macroscopic phase separation of untagged full-length FUS. Specifically, the presence of a dense phase characterized by the appearance of slow modes in the autocorrelation functions, is weakened and abrogated at increased concentrations of HD (Figs. 10A, 10B, 10C) However, profiles of autocorrelation functions, namely the presence of fast modes that are consistent with heterogeneous distributions of clusters, including the mesoscale clusters, persist even upon the addition of up to 1% w/v of HD. This is made clear by comparing Fig. 10B to Fig. 9A. Similar results were obtained when we queried the effects of ATP, which is thought to be condensate dissolving hydrotrope at the concentrations used here (45) – see Fig. 10C as compared to Fig. 9A vs. Fig. 9B. Further, the distributions of clusters formed at *c*_sat_ (≈2 μM) show minimal changes in the presence of HD and / or ATP (Fig. 10D).

**Fig. 10:**
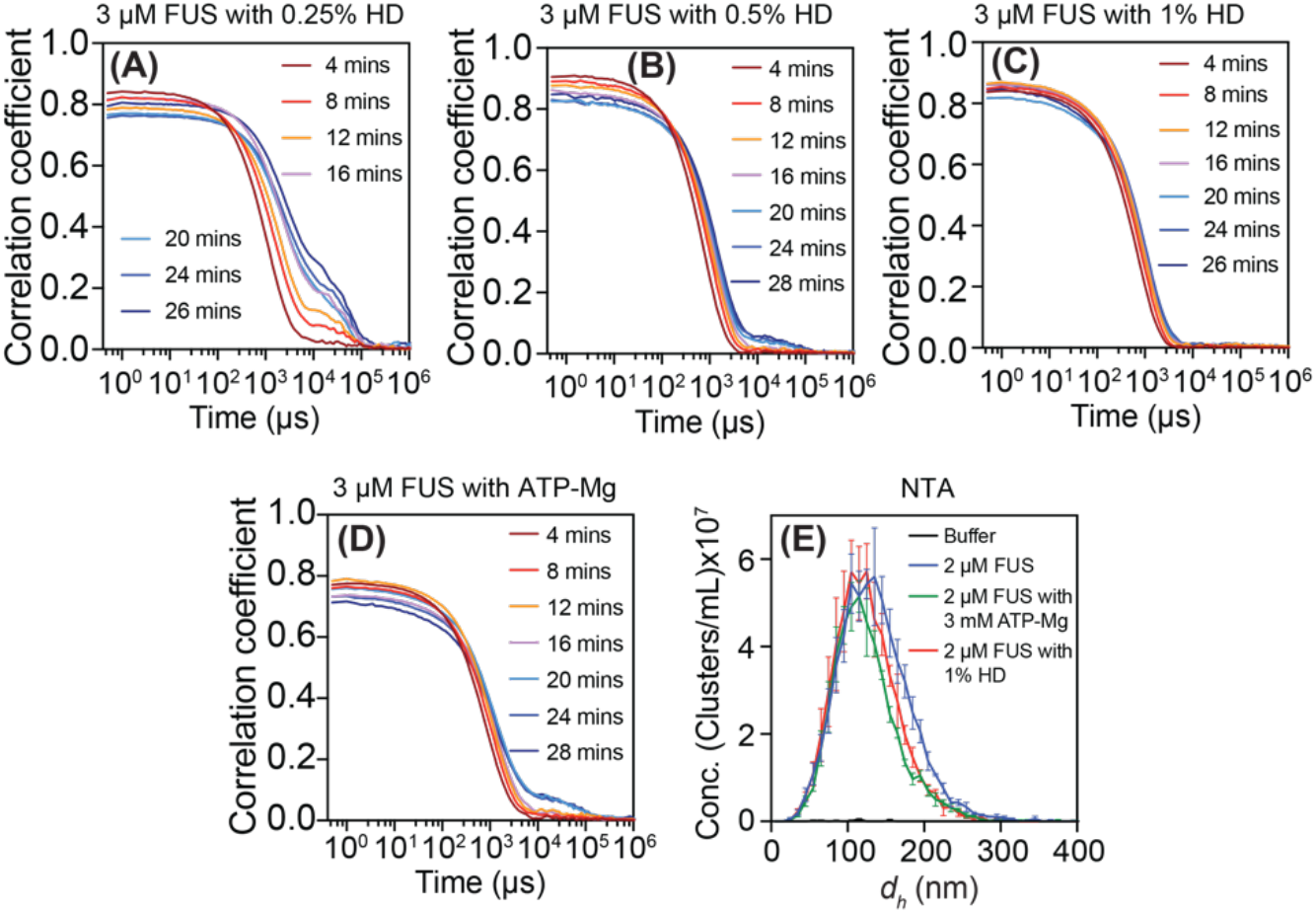
Solutes dissolve condensates while having a minimal effect on clusters. For untagged FUS, above *c*_sat_, there are slow modes in the autocorrelation functions – see Fig. 9B. This feature is weakened in (A) 0.25% and (B) 0.5% and lost in (C) 1% 1,6 hexanediol (HD), which dissolves condensates for concentrations above *c*_sat_. However, clusters that form via fast modes persist upon the addition of HD. (C) Similar observations were obtained in the presence of 3 mM ATP-Mg. (D) NTA data, collected at a concentration of 2 μM for untagged FUS, show that the solutes have discernible effects on the distribution of cluster sizes. Within statistical error of the measurements, the effects of solutes on clusters in subsaturated solutions are small when compared to the impact of solutes on the loss of slow modes above *c*_sat_.

We interpret the results of solute titrations to mean that there are at least two separable energy scales that contribute to condensate formation. We propose that solutes such as HD and ATP primarily affect solvent quality, whereby the Flory χ parameter is altered to impact the overall solubility profiles of FUS molecules. This is consistent with the observations that polyols such as HD lower the macroscopic surface tension of water (46). This will have a generic impact on solvent quality. While solutes such as HD and ATP primarily affect the Flory χ parameter and macroscopic phase separation influenced by *c*_sat_, they have a minimal effect on cluster formation (Fig. 10D). We reason that cluster formation is likely to be influenced by distinct, chemistry-specific interactions, that can be separable using solutes but are generally strongly coupled to driving forces for macroscopic phase separation.

### Impacts of mutagenesis on cluster formation andphase separation

We used a combination of mutagenesis experiments and experiments based on changes to solution conditions to query the extent of coupling between interactions that drive cluster formation vs. macroscopic phase separation. We measured the sensitivity of cluster formation and phase separation of full-length FUS to changes in pH (*SI Appendix*, Fig. S16). These experiments show that increasing the net charge weakens cluster formation in subsaturated solutions (*SI Appendix*, Fig. S16A, B). This leads to a clear upshift of the *c*_sat_ for phase separation (*SI Appendix*, S16D). Likewise, increasing the concentration of monovalent salts also weakens cluster formation (*SI Appendix*, Fig. S17). Therefore, increasing the net charge above a system-specific threshold or screening of electrostatic interactions weaken cluster formation *and* macroscopic phase separation.

Previous studies showed that the sequence-specific *c*_sat_ values of 22 different RNA binding proteins were linearly correlated with (*n*_R_*n*_Y_^−1^ (9), where *n*_R_ and *n*_Y_ refer to the numbers of Arg and Tyr residues that are jointly present within disordered PLDs and RBDs. Reducing the numbers of Arg or Tyr residues shifts *c*_sat_ upward. To test the contributions of chemistry-specific interactions to the coupling between cluster formation and macroscopic phase separation, we replaced 24 Arg residues in the RBD of full-length FUS with Gly residues. These mutations shift *c*_sat_ up by at least an order of magnitude (9) and concomitantly lower the abundance of mesoscale clusters by over an order of magnitude (Fig. 11A, *SI Appendix*, Fig. S18A). Likewise, replacement of aromatic Tyr residues to Ser in the PLD of full-length FUS abrogates phase separation in the low micromolar range while also lowering the abundance of mesoscale clusters by over an order of magnitude (Fig. 11B, *SI Appendix*, Fig. S18B-D). These results highlight a strong coupling between clustering and macroscopic phase separation when Arg or aromatic residues are removed (*SI Appendix*, Fig. S19, S20). In contrast we find that substitution of 10 Asp and 4 Glu residues to Gly within the RBD of full-length FUS stabilizes cluster formation (Fig. 11C, *SI Appendix*, Fig. S18E). Specifically, at a concentration of 2 μM, the abundance of clusters increases by a factor of 1.8 when compared to wild type FUS. However, the increase in abundance of clusters is not accompanied by evidence of condensate formation even at a bulk concentration of 4 μM (*SI Appendix* Fig. S21).

**Fig. 11:**
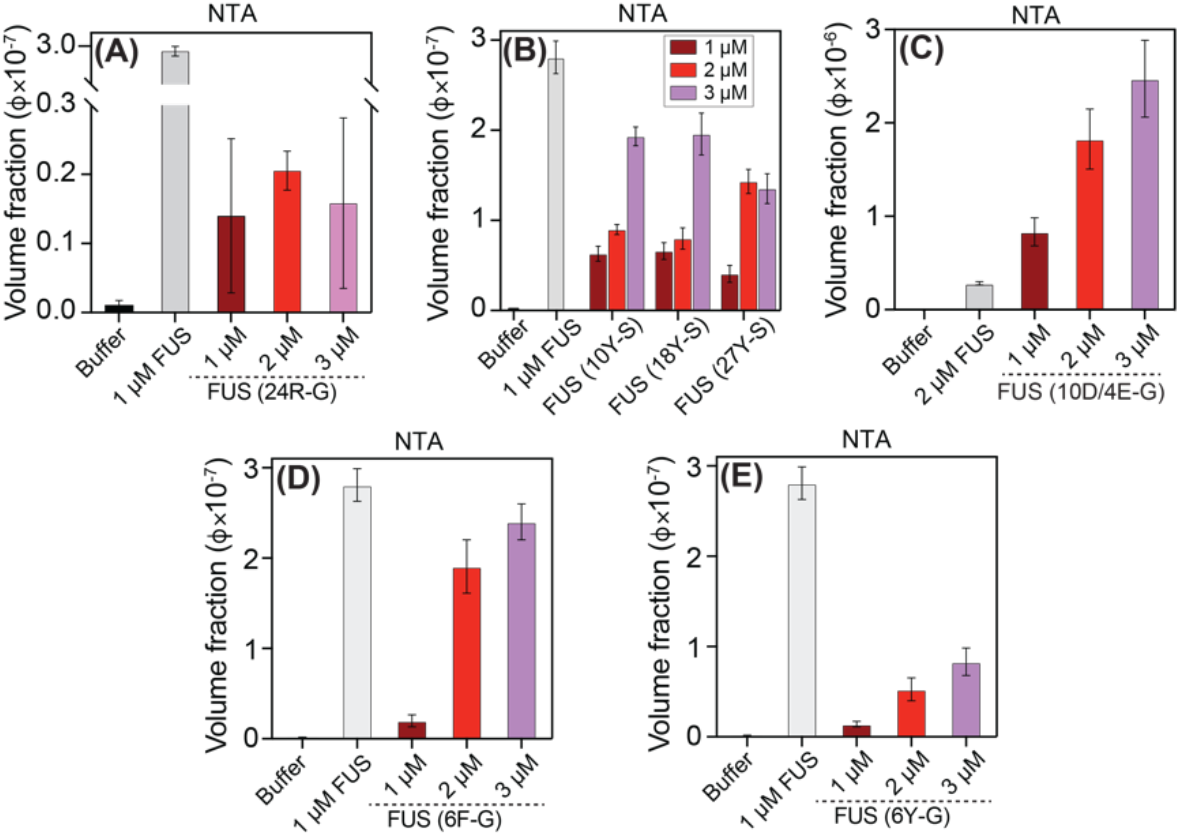
Mutagenesis experiments reveal chemistry-specific effects of different residue types on the formation of mesoscale clusters in subsaturated solutions of FUS. (A) NTA data show that substituting 24 Arg residues to Gly within the RBD reduces the abundance of mesoscale clusters. (B) Replacing aromatic residues within the PLD of full-length FUS shows a valence-dependent reduction in the abundance of mesoscale clusters. (C) Replacing Asp / Glu with Gly in the RBD of full-length FUS, enhances chemistry-specific interactions that stabilize clusters. (D) Substitution of six Phe residues with Gly in the RBD of full-length FUS lowers the abundance of clusters. (E) Likewise, substitution of six Tyr with Ser in the RBD of full-length FUS also lowers the abundance of clusters.

Next, we queried the impact of replacing six of the Phe residues within the RBD of full-length FUS to Gly (Fig. 11D, *SI Appendix*, Fig. S18F) vs. replacing six of the Tyr residues in the RBD to Ser (Fig. 11E, *SI Appendix*, Fig. S18G). Although both Phe and Tyr are aromatic residues, the effects of replacing these moieties are quantitatively different. At equivalent concentrations, the abundance of mesoscale clusters, detected using NTA, is at least three-fold lower when Tyr residues are substituted with Ser when compared to substituting Phe residues with Gly. Therefore, in FUS, Tyr is a stronger driver than Phe of clustering in subsaturated solutions. In both cases, macroscopic phase separation is also not observed for bulk concentrations up to 10 μM (*SI Appendix*, Fig. S22). The differences between Phe and Tyr are reminiscent of recent results for the PLD of hnRNP-A1 (47).

### Impact of the diversity of chemistry-specific interactions on cluster formation and phase separation

The distribution of residue types is different across full-length FUS compared to the PLD and RBD alone (Fig. 12A). The measured *c*_sat_ values of PLDs are at least two orders of magnitude higher than those of full-length proteins (9, 48). We asked whether this change in *c*_sat_ was accompanied by changes in cluster formation. Indeed, NTA measurements show that at equivalent molar concentrations, the abundance of mesoscale clusters formed by the PLD of FUS is lower by a factor of 60 when compared to the abundance of such clusters formed by full-length FUS (Fig. 12B, *SI Appendix*, Fig. S23A). The DLS measurements show that macroscopic phase separation of the PLD was not observed for concentrations up to 150 μM (*SI Appendix*, Fig. S23B). The sequence of the RBD features a significant number of aromatic and anionic residues and its measured *c*_sat_ is ≈20 μM (*SI Appendix*, Fig. S23C). This is an order of magnitude lower than that of the PLD. NTA data show that at a molar concentration of 1 μM, mesoscale clusters formed by the RBD (Fig. 12C, *SI Appendix*, Fig. S23D) are at least ten times more abundant than clusters formed by the PLD.

**Fig. 12:**
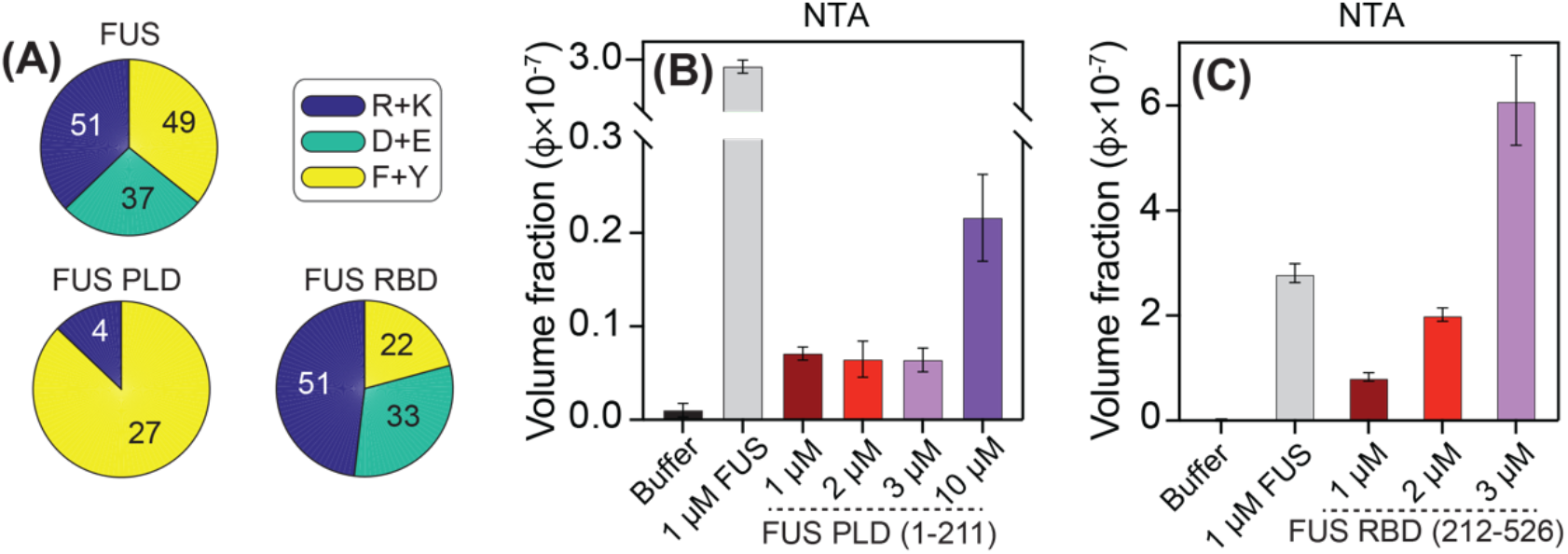
Alterations to chemistry-specific interactions impact the abundance of clusters. (A) RBDs and PLDs of FUS feature differences in amino acid compositions. (B) NTA data show that the lower valence and absence of Arg-π interactions reduces the abundance of mesoscale clusters for the PLD of FUS by roughly two orders of magnitude when compared to full-length FUS. (C) In contrast, the RBD of FUS forms mesoscale clusters that are an order of magnitude more abundant than the PLD.

### Simulations show that systems featuring at least two energy scales behave very differently from those with a single energy scale

The simplest way to introduce distinct energy scales into a polymeric system is via the framework of associative polymers (9, 22, 25–28, 49–53). These are macromolecules with attractive groups known as *stickers* that are interspersed by *spacers* (22, 24, 25, 51, 54). Stickers can form non-covalent reversible crosslinks with one another (29, 47, 55, 56). The strengths of inter-sticker interactions are governed by the functional groups that define stickers. Therefore, in a stickers-and-spacers framework, inter-sticker interactions are specific interactions whereas spacers define the intrinsic χ and contribute to the overall solubility profile of the polymer (29, 47, 55, 56).

We used the LaSSI simulation engine, developed for lattice models of polymers (26), to model systems featuring a single energy scale vs. models for associative polymers that feature at least two energy scales. First, we considered a model homopolymer where all units are identical, (column 1 in Fig. 13). Here, all interactions are uniform and isotropic, and they are described by single energy scale, ε. For the finite-sized homopolymer considered here, phase separation requires that ε be negative (χ must be positive), and the magnitude of ε must be greater than 0.1*k_B_T*. Results for ε values of −0.15*k_B_T* and −0.2*k_B_T* are shown in rows 1 and 2 of column 1 in Fig. 13. Making ε stronger by factor of 4/3, realized by lowering ε from −0.15*k_B_T* and −0.2*k_B_T*, decreases φ_sat_ by three orders of magnitude (column 1, rows 1 and 2 in Fig. 13). This is because phase separation is a highly cooperative process when it is governed by a single energy scale (21). Above a threshold value, small changes to the energy scale ε and hence χ will dramatically alter the overall solubility, and this is manifest as large changes to *c*_sat_. The dense phases formed by homopolymers are essentially polymer melts with volume fractions of φ_den_ ≈ 1 (*SI Appendix*, Fig. S24).

**Fig. 13:**
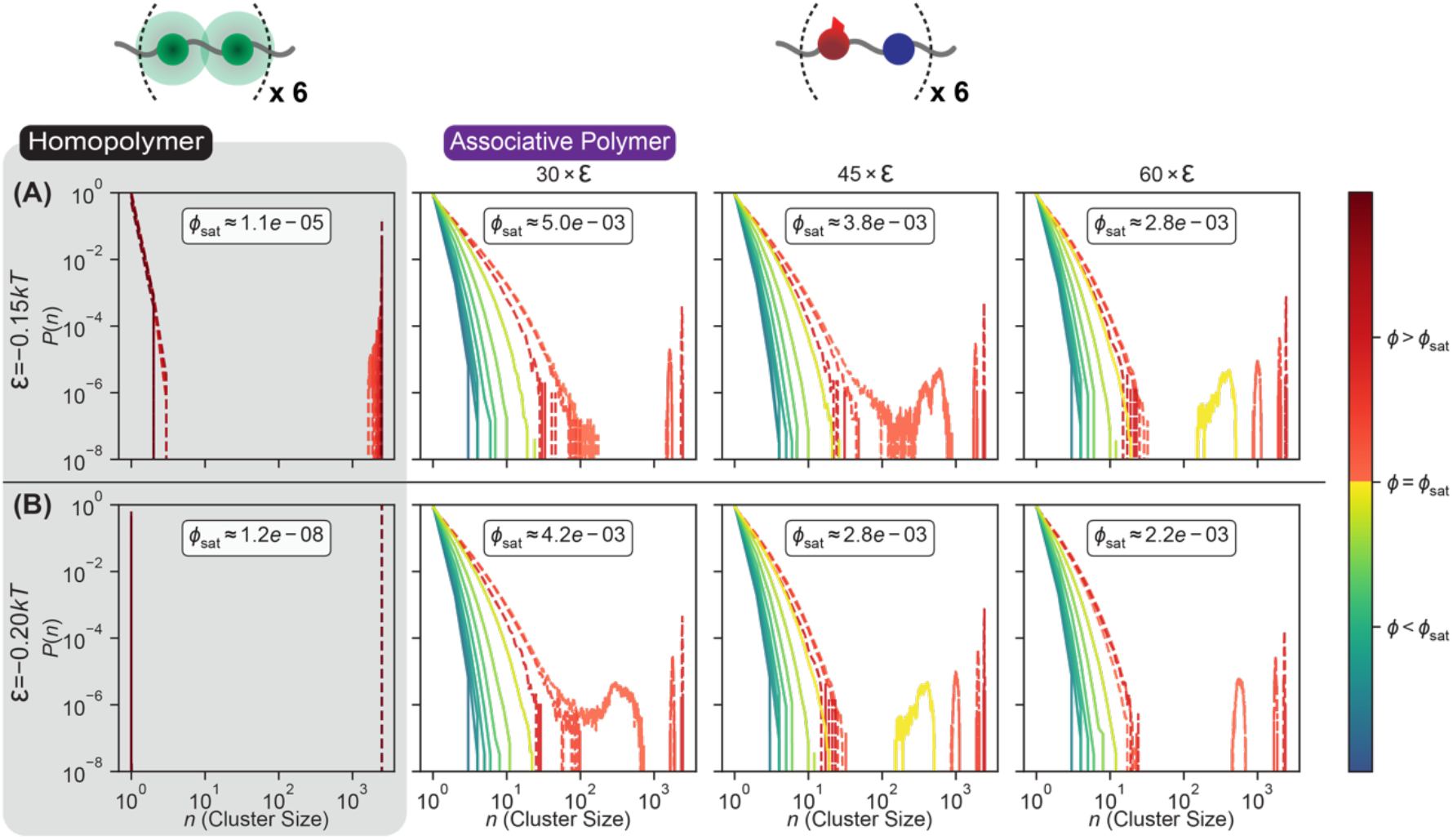
Results from LaSSI simulations highlight the distinctions between homopolymers governed by a single energy scale vs. associative polymers with two energy scales. Each polymer has twelve beads. For homopolymers (results shown in column 1), all beads are the same. For associative polymers, the red beads are stickers, and the blue beads are spacers. We quantify the probability *P*(*n*) of observing clusters comprising *n* molecules. In each panel, the dashed lines represent distributions computed from LaSSI simulations that were performed above the modelspecific *c*_sat_, and the solid lines represent distributions computed in subsaturated solutions. The colors of the dashed and solid lines represent the extent of supersaturation (*s* > 0 for φ > φ_sat_) or subsaturation (*s* < 0 for φ < φ_sat_), respectively. Results in column 1 are for the homopolymer described by a single energy scale ε, whereas results in columns 2-4 are for associative polymers featuring two energy scales namely, the excluded volumes of spacers and the specific sticker-sticker interactions with anisotropic interactions worth *g*ε, where *g* = 30, 45, and 60, respectively.

The introduction of distinct energy scales alters the overall phase behavior. In our simple model for associative polymers, each bead is designated as being a sticker or a spacer (red vs. blue beads in Fig. 13). Stickers form specific anisotropic interactions with one another. For a given value of ε, the strengths of anisotropic inter-sticker interactions are set to be *g*ε, where *g* equals 30, 45, or 60 (columns 2-4 in Fig. 13). The spacers do not engage in attractive spacer-spacer or spacer-sticker interactions. Their only contribution is to the generic excluded volume. For values of ε where the homopolymers undergo phase separation, we observe three distinct effects of introducing the two different energy scales. First, the value of the saturation concentration φ_sat_ is renormalized to be larger than that of the equivalent homopolymer. Although φ_sat_ decreases with increasing *g*, the decrease is considerably smaller when compared to what is observed for the homopolymer when the magnitude of ε is increased by similar extents. Second, heterogeneous distributions of clusters form in subsaturated solutions. The size distributions of clusters are heavy tailed, and their quantitative features are sensitive to both *g* and ε. This highlights the coupling between the effects of specific sticker-sticker interactions and solubility determining sticker-spacer as well as spacer-spacer interactions. Third, the interplay between stickers and spacers causes a dilution of polymer concentrations in the dense phase (*SI Appendix*, Fig. S24). Instead of a polymer melt, the dense phase is a condensate-spanning network defined by a network of inter-sticker crosslinks that is diluted by spacers.

## Discussion

In this work, we characterized species that form in subsaturated solutions of phase separating RNA binding proteins with disordered PLDs and RBDs. For a purely phase separation-based process described by classical Flory-Huggins style theories (10, 11), subsaturated solutions should feature dispersed monomers and very few small clusters, if any, at any given time. It is worth noting that these simple theories have been the mainstay for quantitative descriptions and analysis of phase transitions driven by multivalent proteins and nucleic acids *in vitro* and in cells (12–15, 21, 42, 57–62). Surprisingly, we find that subsaturated solutions do not conform to expectations based on systems that are characterized by a single energy scale such as the Flory *χ*-parameter.

Our observations suggest that the overall phase behavior of FUS and FUS-like molecules is governed by more than one energy scale. The presence of different energy scales is best illustrated by the response to solutes, in which clusters and condensates respond differently to HD and ATP. Phase separation is likely driven mainly by composition-specific interactions that determine the solubility profiles of FUS and FUS-like molecules. However, our data also show that the sequence features of FUS and FUS-like molecules can engender a strong coupling between the driving forces for cluster formation and phase separation. Mutations to stickers weaken cluster formation and increase *c*_sat_. This indicates that the mechanism of phase separation is intimately tied to the structure of the underlying size distribution of clusters that form in subsaturated solutions. The picture that emerges is of networks of chemistry-specific, inter-residue interactions (7, 9, 47, 48, 63) driving cluster formation and determining the extent of coupling between cluster formation and phase separation.

Mesoscale clusters of low overall abundance have been reported for subsaturated solutions of folded proteins that form crystalline solids (64, 65). These clusters, discovered by Gliko et al. (64), were found to have fixed size, irrespective of protein concentration, and are likely to be manifestations of microphase separation (30). In contrast, we observe continuous growth in the sizes and abundance of clusters with concentration. As discussed below, and demonstrated using simulations summarized in Fig. 13, the framework of associative polymers provides a plausible explanation for our observations. Associative polymers undergo two types of transitions namely, *percolation without phase separation* – also known as sol-gel transitions – or *phase separation coupled to percolation* (Fig. 14) (24–27, 52, 53). Above a system-specific threshold concentration known as the percolation threshold or *c*_perc_ (26–28, 53), non-stoichiometric clusters become connected into a percolated or system-spanning network (26–28, 53). Quantitatively, *c*perc is determined by the numbers (valence) of stickers, diversity of sticker types (28), and the strengths of inter-sticker interactions (27, 28). If *c*_perc_ < *c*_sat_, then associative polymers undergo percolation without phase separation. Alternatively, if *c*_sat_ < *c*_perc_ < *c*_den_, where *c*_den_ is the concentration of the dense phase formed via phase separation (53, 66), then phase separation and percolation are coupled (Fig. 14). In this scenario, the condensates that form are akin to spherical microgels (67) whereby the percolated network is condensate spanning (Fig. 14). Tanaka recognized that the effects of specific inter-sticker interactions can be cast as a renormalized Flory χ-parameter whereby χ shifts to χ′=χ+Δχ such that: 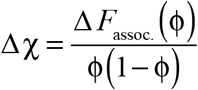 (22). Here, ϕ is the volume fraction of the polymer of interest. Accounting for specific inter-sticker interactions introduces a composition dependence to the renormalized *χ*-parameter, and this describes the formation of clusters of different sizes even in deeply subsaturated solutions (22, 23). Both the renormalization of χ and hence *c*_sat_ as well as the formation of dense phases through phase separation coupled to percolation, as pictured in Fig. 14, are readily observed in the LaSSI simulations when the model accounts for specific anisotropic inter-sticker interactions that are distinct from spacer-mediated, solubility determining interactions.

**Fig. 14:**
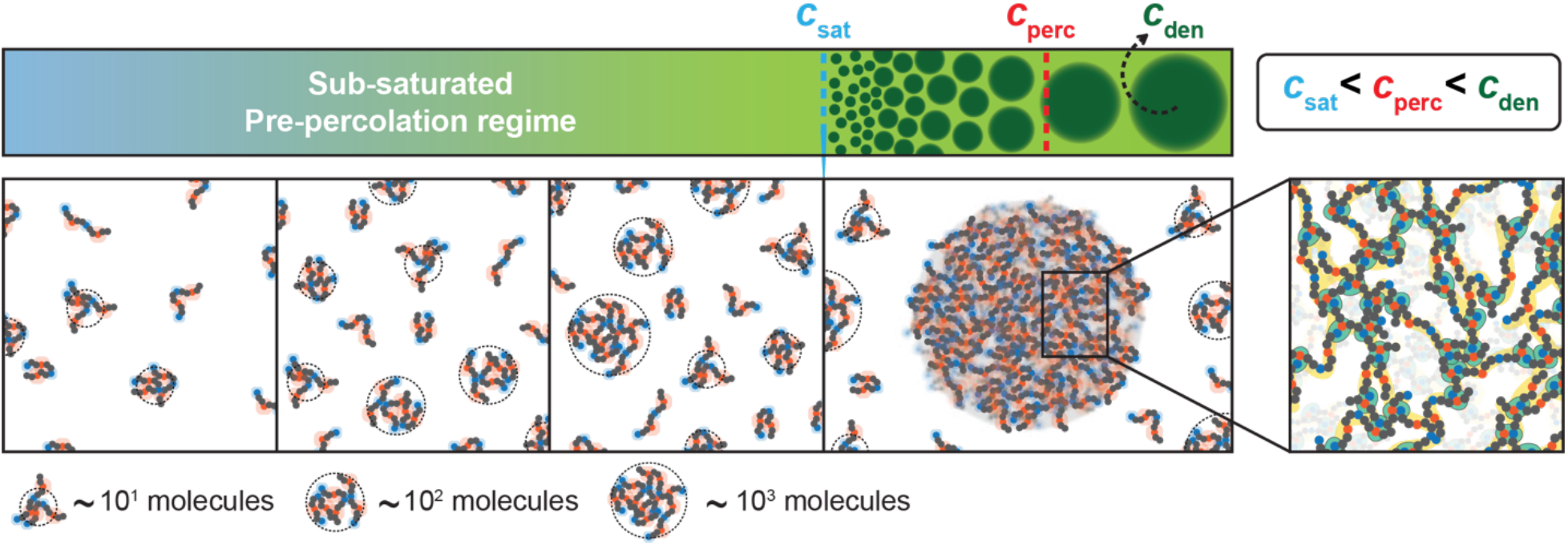
Phase transitions of associative polymers. Schematic showing that reversible physical crosslinks among networks of stickers lead to finite-sized clusters whose sizes and abundance grow continuously with concentration even below *c*_sat_. Above *c*_sat_, clusters grow through via macroscopic phase separation, mainly via monomer addition and coalescence, to form condensate spanning networks, a feature that emerges because *c*_sat_ < *c*_perc_ < *c*_den_.

### Precedents for clusters in subsaturated solutions

The multivalent domain-linker systems studied by Rosen and coworkers are exemplars of associative polymers featuring an interplay between site-specific domain-motif interactions and linker-mediated solubility profiles (53, 66, 68–71). In their original work Li et al. (66) investigated the phase behaviors of systems where phase transitions are driven by multivalent interactions among poly-SH3 and poly-PRM molecules. Existence of a threshold concentration for phase transitions requires that there be at least three SH3 domains and three proline-rich modules (PRMs) within the associating molecules. This is consistent with the presence of a valence-dependent percolation threshold (72, 73). Li et al. (66) showed that as the valence (numbers) of SH3 domains and PRMs increases, the threshold concentration for phase transitions decreases. Importantly, Li et al. (66) reported the presence of clusters in subsaturated solutions. These were detectable using DLS and small-angle x-ray scattering. Further, the phase transitions were referred to by Li et al., as *macroscopic liquid-liquid phase separation that is thermodynamically coupled to sol-gel transitions*. This phrasing is synonymous with the process we describe here as *phase separation coupled to percolation*.

### Biological relevance of clusters that form in subsaturated solutions

Our results lead us to propose that RNA binding proteins from the FET family belong to the class of polymers known as associative polymers. Strong, inter-sticker interactions have been predicted to suppress phase separation and percolation by promoting the formation of clusters of precise stoichiometries through saturating interactions (74). In contrast, finely tuned hierarchies of sticker vs. spacer interactions with average interactions being of order 2-5*k_B_T* can enable strong coupling between cluster formation and phase separation (Fig. 13)(75). Our findings provide a bridge that connects recent findings regarding dynamic clusters comprising of multivalent molecules (76, 77). They also show that chemistry- or site-specific sticker-mediated interactions and solubility determining spacer-mediated interactions can be separable (27, 29, 47, 48, 52, 53, 78–84), and suggested to be of importance for biological functions (85).

Heterogeneous distributions of pre-percolation clusters are likely to be present at endogenous concentrations, which tend to be in the nanomolar to low micromolar range in live cells (86). What then are the potential consequences of clusters in subsaturated solutions in cells? Process control via biomolecular condensates requires that phase separation be robust and reproducible (3). In a classical phase separation system, the barrier to nucleation will determine the response time. Clusters will likely speed up the response time by lowering the barrier for phase separation. Accordingly, we postulate that mesoscale clusters poise RNA binding proteins for robust condensate formation in response to stimuli. Importantly, to date, most work on regulation of biological condensates has focused primarily on regulation of macroscopic phase separation (59, 78, 87). Our discovery that heterogeneous distributions of clusters exist in subsaturated solutions suggests that regulation could occur at the level of shaping the distribution of cluster sizes. For example, chaperones are known to modulate size distributions of self-associating molecules (12), and we could easily imagine chaperones acting to reshape the size distributions of clusters described in this work. Additionally, deleterious interactions with cellular components that lead to disease (88), and the dynamical arrest of condensate growth (89) could also occur at the level of clusters.

## Materials and Methods

Details of all the constructs used in the experiments, the protein expression and purification protocols that distinguish methods A and B, the setup and analysis of DLS, NTA, MFD, MCS, and TEM experiments, raw data from DLS measurements, and the LaSSI simulations are described in the *SI Appendix*.

## ACKNOWLEDGMENTS

We thank Mina Farag, Tyler Harmon, Adam Klosin, Alex Holehouse, Tanja Mittag, Doayuan Qian, Kiersten Ruff, Vijay Rangachari, Samuel Safran, Jie Wang, and members of the Hyman and Pappu labs for helpful discussions. This work was funded by a direct grant from the Max Planck Society (to A.A.H.), a grant from the NOMIS foundation (to A.A.H), the Wellcome trust (209194/Z/17/Z to A.A.H.), the European Research Council through the ERC grant PhysProt (to T.P.J.K, agreement no. 337969), the Wellcome Trust and the Frances and Augustus Newman foundation (to T.P.J.K.), SPP2191 from the Deutsche Forschungsgemeinschaf (to S.A. and C.A.M.S), the US National Institutes of Health (5R01NS1056114 and R01NS121114 to R.V.P), and the St. Jude Children’s Research Hospital collaborative research consortium on membraneless organelles (to R.V.P). We are grateful for technical support provided by Régis Lemaitre and Barbara Borgonovo of the protein expression, purification, and characterization facility of the MPI-CBG, as well as Michaela Wilsch-Bräuninger and Jana Mesenser of the TEM facility at MPI-CBG. We thank Andrei Pozniakovsky for DNA constructs of all proteins.

