## Supporting Information Appendix for "Phase separating RNA binding proteins form heterogeneous distributions of clusters in subsaturated solutions"

**Section A. Materials**

**Table S1: List of reagents, sources, and vendor identifiers if any**

| REAGENTS | SOURCE | IDENTIFIER |
|---|---|---|
| **CHEMICALS** | | |
| TRIS | Carl Roth Germany | 77-86-1 |
| Potassium Chloride (KCl) | Merck Germany | 7447-40-7 |
| Glycerol | VWR chemicals | 56-81-5 |
| cOmplete$^{TM}$ | Roche Germany | 11697498001 |
| Imidazole | Sigma Aldrich Germany | 288-32-4 |
| DTT | Alfa Aesar Germany | 578-51-7 |
| Hexane di-ol | Sigma Aldrich Germany | 629-11-8 |
| ATP-Na | Jena Bioscience | 987-65-5 |
| Maltose | Sigma Aldrich | 6363-53-7 |
| **Bacterial and Virus Strains** | | |
| Sf9 cells | Expression Systems | Cat#94-001F |
| **Recombinant proteins** | | |
| FUS GFP | Wang et. al. 2018 (1) | TH1204 |
| FUS SNAP | Wang et. al. 2018 (1) | TH0901 |
| FUS | Wang et. al. 2018 (1) | TH0901 |
| FUS PLD (1-211) | Wang et. al. 2018 (1) | TH0994 |
| FUS RBD (211-526) | Wang et. al. 2018 (1) | TH0996 |
| FUS (RBD, 24R-G) | Wang et. al. 2018 (1) | TH1006 |
| FUS (RBD, 10D/4E -G) | This study | TH1740 |
| FUS (PLD,10Y-S) | This study | TH1427 |
| FUS (PLD, 18Y-S) | This study | TH1815 |
| FUS (PLD, 27Y-S) | Wang et. al. 2018 (1) | TH0992 |
| FUS (RBD, 6F-G) | This study | TH1917 |
| FUS (RBD, 6Y-S) | This study | TH1918 |
| TAF15 SNAP | Wang et. al. 2018 (1) | TH1203 |
| EWSR1 SNAP | Wang et. al. 2018 (1) | TH1276 |
| hnRNPA3 | Wang et. al. 2018 (1) | TH1202 |
| TDP 43-GFP | Wang et. al. 201 (1)8 | TH3006 |

**Constructs, protein expression and purification: Method A**

The construct / protein sequences used are listed in Section C. Details of materials used for the preparation of samples are as follows:

*Lysis buffer*: 50 mM Tris-HCl pH 7.4, 1 M KCl, and 5% Glycerol.

*Protease inhibitor*: cOmplete™, EDTA-free Protease Inhibitor Cocktail Tablets.

*NTA elution buffer*: 50 mM Tris-HCl pH 7.4, 1 M KCl, 5% Glycerol and 300 mM Imidazole. *MPB elution buffer*: 50 mM Tris-HCl pH 7.4, 1 M KCl, 5% Glycerol and 30 mM Maltose. *Storage buffe*r: 50 mM Tris-HCl pH 7.4, 500 mM KCl, and 5% Glycerol, and 1 mM DTT.

*Purification method*: All proteins were first expressed in 1 L SF9 (1 million/mL) insect cells with 5 mL P2 virus and harvested 72 hours post infection. Cells were collected by centrifugation for 5 min at 2,000 rpm. The pellets were re-suspended in lysis buffer. Protease inhibitors were added to the lysis buffer solution. The cells were lysed by sonication. The crude lysate was clarified by centrifugation for 20 min at 17,000 rpm. After centrifugation, the supernatant was passed through with Ni-NTA agarose column using Ismatec peristaltic pump. The protein-bound beads inside the column were further washed with 10 column volumes of lysis buffer and the mixture of 10 mM of Imidazole with lysis buffer, respectively, to remove non-specific bound proteins from the Ni-NTA column. The proteins were eluted with the NTA elution buffer.

The eluted protein solution was passed through MBP resins using gravimetric column. The protein bound beads were washed with 10 column volume of lysis buffer. The proteins were eluted with the MBP elution buffer. The entire process was monitor using Bradford assay solution and each fraction were investigated with Gel electrophoreses.

To cleave the His-MBP tag, 3C prescission protease was added to the eluted protein at a 1:100 ratio. The mixture was incubated at RT for 4 hours and was purified over the gel filtration chromatography (ÄKTA with Superdex-200 increase column; GE Healthcare) equilibrated with storage buffer. Peak fractions were pooled and immediately use for the DLS or NTA experiments. Further, to obtain untagged protein TEV protease were added at a 1:50 ratio and incubated at RT for 6 hours. The untagged protein was purified over the gel filtration chromatography (ÄKTA with Superdex-200 increase 10/300 column; GE Healthcare) equilibrated with storage buffer. Peak fractions were pooled and concentrated using Amicon 15 30.000 MWCO at 4000 rpm in RT. Protein concentration was determined by measuring absorbance at 280 nm using a NanoDrop ND-1000 spectrophotometer (Thermo Scientific). The 260/280 ratio of all purified proteins were measured between 0.52 to 0.56. (Note: Proteins were stored with tags at -80 C, and prior to experiments the proteins were thawed, tag was cleaved, run gel filtration chromatography, peak fractions were pooled and concentrated and used for the experiments. This way the data reproducibility was good. If the proteins were frozen and thawed, the reproducibility of the experiments varies.)

**Constructs, protein expression and purification: Method B**

FUS SNAP protein expression and purification using method B. Details of materials used for the preparation of samples are as follows:

*Lysis buffer:* 50 mM Tris/HCl, pH 8.2, 1 M NaCl, 25 mM Imidazole, 5% (w/v) Glycerol, 1 mM DTT.

*Buffer A:* 50 mM Tris/HCl, pH 8.2, 1 M NaCl, 25 mM Imidazole, 5% (w/v) Glycerol, 1 mM DTT.

*Buffer B*: 50 mM Tris/HCl, pH 8.7, 1 M NaCl, 250 mM Imidazole, 5% (w/v) Glycerol, 1 mM DTT.

*Buffer C*: 50 mM Tris/HCl, pH 8.7, 1 M NaCl, 250 mM Imidazole, 5% (w/v) Glycerol, 1 mM DTT, 10 mM Maltose.

*Buffer D*: 50 mM Tris/HCl, pH 7.5, 0.5 M KCl, 5% (w/v) Glycerol, 1 mM DTT.

*Purification method*: FUS-SNAP was expressed in 2 L SF9 (2 million/mL) insect cells with 20 mL P3 virus and harvested 72 hours post infection. Cells were collected by centrifugation for 5 min at 1,500 rpm. The pellets were re-suspended in lysis buffer. Protease inhibitors were added to the lysis buffer solution with 100 μL of Benzonase. The cells were lysed by using a shear fluid homogenizer (LM10 Microfluidizer from Microfluidics) at 5000 psi. The crude lysate was clarified by centrifugation for 60 min at 25,000 rpm at 15°C.

3 x 5 mL HisTrap Cytiva Columns Position 2, 25 mL HiPrep Amylose (NEB HiFow Amylose 25 ml XK16/20) Position 8, and 320 mL Superdex 200 pg 26/60 Column Position 7 were mounted in ÄKTA explorer. HisTrap Columns were equilibrated with Buffer A, loading cell lysate followed by washing with Buffer A and elution with Buffer B (3 column volume). HiPrep Amylose column was equilibrated with Buffer B, loading the eluted protein from HisTrap Columns, followed by washing with buffer B. 3C PreScission Protease was loaded into the Amylose column and incubated for 8 hours at RT. The protein was eluted using Buffer B. The Amylose column was regenerated using Buffer C. Superdex 200 pg column was equilibrated with Buffer D. The eluted protein from Amylose column was loaded and run the SEC using Buffer D. Peak fractions were pooled and concentrated using Amicon 15 10.000 MWCO at 15000 rpm in RT. Protein concentration was determined by measuring absorbance at 280 nm using a NanoDrop ND-1000 spectrophotometer (Thermo Scientific). The 260/280

ratio of all purified proteins were measured between 0.55. The concentrated protein was aliquoted, and flash frozen with liquid $N_2$.

**Section B. Measurement and Computational Methods**

**Dynamic light scattering experiments (DLS)**

DLS measurements were performed using the Zetasizer Nano ZSP Malvern instrument (measurement range of 0.4 nm to 10 µm). The Nano ZSP instrument incorporates noninvasive backscattering technology. This enables the measurement of time-dependent fluctuations of the intensity of scattered light as scatterers undergo Brownian motion. The analysis of these intensity fluctuations enables the determination of the diffusion coefficients of particles, which are converted into a size distribution using the Stokes-Einstein equation (2). The sample solutions were illuminated by a 632.8 nm laser, and the intensity of light scattered at an angle of 173° was measured using a photodiode.

In DLS, the autocorrelation function of the scattered light is used to extract the size distribution of the dissolved particles. The first order electric field correlation function of laser light scattered by a monomodal or monodisperse population of macromolecules can be written as a single exponential of the form:

$$G(\tau) = 1 + b\exp\left(-2D_t q^2 \tau\right); \quad \text{(S.1)}$$

Here, $b$ is a constant that is determined by the optics and geometry of the instrument, $D_t$ is the translational diffusion coefficient of the particles, and $\tau$ is the characteristic decay time. The scattering vector $q$ is given by:

$$|q| = \frac{4\pi n_0}{\lambda_0}\sin\left(\frac{\theta}{2}\right); \quad \text{(S.2)}$$

Here, $n_0$ is the refractive index of the solvent, $\lambda_0$ is the wavelength, and $\theta$ is the scattering angle. For populations composed of a single type of scatterer, the distribution function of decay rates can be derived from a simple fit of the experimental estimates of the logarithm of the correlation function in Eq. S.1 to a polynomial. These methods, which apply to monomodal distributions of sizes of scatterers can be used to extract the translational diffusion coefficient, from which one can estimate the hydrodynamic radius $R_h$ of the scatterers. For this, one uses the Stokes–Einstein relation:

$$D_t = \left(\frac{k_B T}{6\pi\eta R_h}\right); \quad \text{(S.3)}$$

Here, $k_B$ is the Boltzmann constant ($1.381 \times 10^{-23}$J/K) and η is the absolute (or dynamic) viscosity of the solvent. In this work, we used the hydrodynamic diameter $d_h$ (i.e., $d_h = 2R_h$) as preferred way to quantify particle sizes.

*DLS measurements:* All solutions were filtered using 0.2 μm membranes (Millex®-GS units) purchased from Millipore™. All experiments were conducted with following settings on the Malvern instrument: Material – protein; Dispersant – 20 mM Tris buffers with 100 mM KCl salts; Mark-Houwnik parameters; Temperature: 25 °C with equilibration time – 120 seconds, Measurement angle: 173˚. Each spectrum represents the average of 12 scans, each 10 seconds in duration. For every measurement, we recorded the autocorrelation function, intensity, and number.

All proteins were freshly purified and used after chromatography purification with standard stock solution buffer. For typical measurement, the final buffer composition consists of 20 mM Tris 7.4 and 100 mM KCl. The samples were prepared by adding freshly prepared stock proteins followed by dilution buffer, and mixed thoroughly by pipetting 4 to 6 times. The samples were equilibrated for 2 mins at 25 °C and the data were recorded in 2-minute intervals.

**Nanoparticle Tracking Analysis (NTA)**

Nanoparticle tracking analysis was performed using NS300 from Malvern instruments (measurement range of 20 nm to 1μm). The system was accompanied with a NanoSight syringe pump to inject the samples for the experiments. NTA measurements utilize the properties of light scattering and Brownian motion to obtain the size distributions and concentrations of particles in liquid suspension. A laser beam (488 nm) was passed through the sample chamber, and the particles in suspension were visualized using a 20x magnification microscope. The video file of particles moving under Brownian motion was captured using a camera mounted on the microscope that operates at 30 frames per second. The software tracks particles individually and uses the Stokes-Einstein equation resolve particles based on their hydrodynamic diameters.

All proteins were freshly purified and used after chromatographic purification with standard stock solution buffer. All buffers were filtered through 0.22 μm polyvinylidene fluoride membrane filter (Merck, Germany). All proteins stock solutions were centrifuged at 20000 RCF for 5 mins at room temperature prior to measurements. For typical measurements, the final buffer consists of 20 mM Tris 7.4 and 100 mM KCl. The samples were prepared by adding freshly prepared and centrifuged stock proteins followed by dilution buffer, and mixed thoroughly by pipetting 4 to 6 times. The samples were equilibrated for 2 minutes at 25˚C and the data were recorded 6 minutes after sample preparation and equilibration.

**Transmission electron microscopy (TEM)**

TEM micrographs were acquired using Morgagni TEM (ThermoFisher) operated at 80kV with a Morada camera (EMSIS). Stock FUS-SNAP was diluted to 2 µM at 20 mM Tris &.4 and 100 mM KCl. 3 µL of sample was loaded on a carbon-coated copper grid, incubated for 6 mins, and then gently drawn off using a filter paper wick. This was followed by addition of 2 µL of 1.5% PTA pH 7.5. After 30 seconds the solution was gently drawn off by filter paper wick.

**Förster resonance energy transfer (FRET)**

For FRET experiments, the SNAP tagged FUS proteins were mixed with SNAP-Surface Alexa Fluor 546 or 488 (NEB) at a 1:1.5 ratio at RT for 2 hr. Free dye was removed using Zeba Spin Desalting Columns (Thermo Scientific, Lot # QH222764) equilibrated with the storage buffer. FUS-SNAP containing 10% dye labelled FUS-SNAP-488 or FUS-SNAP-546 was mixed separately with buffer to yield a final protein concentration of 0.5 µM in 20 mM Tris 7.4 with 20 mM KCl. Then the solutions were mixed in equal volume. A 96 well plate (microplate, PS, half area, µClear, Med. binding, Black, Greiner Bio-one) was loaded with 100 µl of 0.5 µM FUS-SNAP 488, 0.5 µM FUS-SNAP-546, and the mixture. The spectra were recorded from 530 nm to 600 nm (10 nm bandwidth) with the TECAN plate reader using an excitation wavelength of 460±10 nm. For control, we used only Alexa 488 and Alexa 546 in similar range of concentrations with same buffer conditions. Additionally, we used high salt conditions (200 mM KCl) to dissolve the clusters and recorded the spectra at same settings.

**Anisotropy Measurements**

*Setup*. The anisotropy measurements were conducted on a confocal fluorescence microscope (FV1000 Olympus, Hamburg, Germany) using a polarized pulsed diode-laser (LDH-D-C-485, PicoQuant, Berlin, Germany) at 485 nm. Laser light was directed into a 60x water immersion objective (NA=1.2) by a dichroic beam splitter and focused into the sample close to the diffraction limit. The light emitted was collected by the same objective and separated into two polarizations (parallel and perpendicular) relative to the excitation beam. The fluorescence signal was further divided into two spectral ranges (BS 560, AHF, Tübingen, Germany). Bandpass filters for eGFP fluorescence (HC 525/39) were placed in front of the detectors. The signal from single photon sensitive detectors (PDM50-CTC, Micro Photon Devices, Bolzano, Italy and HPMC-100-40, Becker&Hickl, Berlin, Germany, respectively) was recorded photon-by-photon with picosecond accuracy (HydraHarp400, PicoQuant) and analyzed using custom software (LabVIEW based). The temperature during all titration steps was 21.5 ± 0.5 °C.

The anisotropy of Nile red with FUS-SNAP as titrant was measured with the same confocal setup using a supercontinuum laser (SuperK Extreme, NKT Photonics, Birkerød, Denmark) at 514 nm. The spectral ranges were separated by dichroic beamsplitters (BS 560 and 630 DCXR, AHF, Tübingen, Germany). Bandpass filters for eGFP (HC 525/39) and Nile red fluorescence (HC 607/70, HC 715/120) were placed in front of the detectors.

*Titration procedures*. FUS-SNAP stained with 100 nM FUS-eGFP (both prepared using method B) were measured in a 50 mM Tris-HCl (pH 7.6) solution on a FUS coated cover glass. To account for the increasing KCl concentration with FUS-SNAP titration Tris-HCl buffer was added accordingly keeping the salt content constant until the saturation concentration resulting in a total eGFP dilution of 38 v%. FUS-SNAP was titrated into 20 nM Nile red solution at 50 mM KCl and 50 mM Tris-HCl (pH 7.6 at 21 °C) whilst keeping both dye and salt concentration constant with Nile red containing buffer. The brightness of Nile red increases in a hydrophobic environment due to longer fluorescence lifetime shifting the emission spectrum to shorter wavelenghts. Due to the direct binding of Nile red to FUS the anisotropy reflects the actual rotational diffusion of FUS oligomers not obscured by dye-linker motions as seen in the eGFP measurements.

**Microfluidic Confocal Spectroscopy (MCS)**

Microfluidic devices, shown by the design in Fig. S7 (3), were first fabricated as SU-8 molds (MicroChem) through standard photolithographic processes, and then produced as polydimethylsiloxane (PDMS) slabs, which were bonded onto thin glass coverslips (4). The devices were operated by placing gel-loading tips filled with buffer and protein sample in their corresponding inlet ports (Figure S1B) and pulling solution through the devices in withdraw-mode at a flow rate of 150 μL/h using automated syringe pumps (neMESYS, Cetoni).

All experiments were conducted with FUS-eGFP fusion protein prepared using method A. The protein, stored in 500 mM KCl, 20 mM TRIS-HCl pH 7.4, was diluted with buffers of 20 mM TRIS-HCl to the indicated protein and KCl concentrations as stated. During the experiment, the sample was placed into the sample inlet of the device (Fig. S7A) and the corresponding buffer containing the same concentration of KCl and TRIS-HCl into the buffer inlet. The co-flowing buffer was supplemented with 0.05% Tween-20 to prevent surface sticking of the protein to PDMS and glass surfaces.

Experiments were conducted by scanning the confocal spot of a custom-built confocal microscope through the central four channels of the microfluidic device (insert in Fig. S7A). A schematic of the optical unit is shown in Fig. S7B. Briefly, the setup is equipped with a 488-nm laser line (Cobolt 06-MLD) for excitation of GFP fluorophores and a single-photon counting avalanche photo diode (SPCM-14, PerkinElmer) for subsequent detection of emitted fluorescence photons. Further details of the optical unit have been described previously. During the scanning of the device, 200 evenly spaced locations within the central four channels of the device were surveyed and detected for 4 seconds. Examples of individual 4 second traces both with and without FUS clusters are shown in Fig. S7C. Clusters were classified as peaks that exceeded 5 standard deviations above the mean fluorescence intensity of each trace. These peaks were quantified according to location (Fig. S7D; top panel) against the mean signal of each trace (Fig. S7D; bottom panel). The average number of clusters $\overline{n}_{\text{clusters}}$ was then quantified by averaging each of the four groups of peaks, corresponding to the four central channels. This was used in the calculation of cluster concentration according to:

$$F_{\text{total}} = \left(\frac{\overline{n}_{\text{clusters}}}{t}\right)\left(\frac{hd_{\text{step}}}{\frac{\pi}{4}zw}\right); \qquad \text{(S.4)}$$

Here, $t$ is the time each trace was collected for (4 seconds), $h$ is the height of the microfluidic channel (28 μm), $d_{\text{step}}$ is the width of each step (5.64 μm), and $z$ and $w$ were the height and width of the confocal spot (3 μm and 0.4 μm, respectively). From Equation (S.4), which yields the flux of clusters $F_{\text{total}}$, the concentration of clusters could be determined according to Equation (S.5), with $Q_{\text{sample}}$ being the flow rate of the sample (15 μL/h) and $N_{\text{A}}$ being the Avogadro constant (5):

$$c_{\text{cluster}} = \left(\frac{F_{\text{total}}}{N_{\text{A}}Q_{\text{sample}}}\right); \qquad \text{(S.5)}$$

**Computational Modeling**

Simulations were performed using a customized version of the LaSSI simulation engine (6). For each of the simulations, a total of $N_{\text{tot}}$ = 2500 polymers were. Twenty different concentrations were sampled and these, written in terms of volume fractions, range from $\phi_{\text{min}} = 2\times10^{-5}$ to $\phi_{\text{max}} = 5\times10^{-2}$. The number of sites, $L$, on the $L\times L\times L$ cubic lattice was decreased from 1013 to 84 to increase the concentration by over three orders of magnitude. Radial density profiles from the centers-of-mass of the largest droplets were used to calculate the coexisting concentrations $\phi_{sat}$ and $\phi_{den}$. For additional details, see the *SI Appendix* in the work of Ruff et al., (7). Cluster size distributions were calculated from the simulations by quantifying the cluster composition of the system where a criterion of a maximal distance of $\sqrt{3}$ is used.

The basic architecture of the two models is the same: polymers that contain 12 beads connected by tether-like linkers of length 2. For the Homopolymer, all beads are of the same type, while for the Associative Polymer we have both stickers and spacers, as shown in Fig. 13.

**Table S2: Frequencies of different Monte Carlo moves**

| Move | Frequency | Relative Frequency |
|---|---|---|
| Anisotropic Small Cluster | 0.24 | 10 |
| Anisotropic Large Cluster | 0.02 | 1 |
| Proximity Small Cluster | 0.24 | 10 |
| Proximity Large Cluster | 0.02 | 1 |
| Local | 39.79 | 1666 |
| Sticker Rotation | 3.98 | 166 |
| Slithering Snake | 19.89 | 833 |
| Chain Translation | 7.96 | 333 |
| Pivot | 7.96 | 333 |
| Co-Local | 7.96 | 333 |
| Multi-Local | 3.98 | 166 |
| Double Pivot | 7.96 | 333 |

Additional details regarding the moves may be found in the original work by Choi et al., (6). *Multi-Local* moves and *Cluster* moves are new to the current version of LaSSI, and they are explained below:

*Multi-Local moves*: In these moves, a bead in the system is picked at random. The selected bead, and the beads that are covalently bonded to it are removed from the lattice. All beads are attempted to be replaced within a 2-lattice site radius of the first bead. Steric clash or incorrect linker lengths results in move rejection. If appropriate lattice-sites are found, the standard Metropolis-Hastings criterion is used for acceptance.

*Cluster Moves*: In general, these moves attempt to translate certain clusters in the system. For the *Anisotropic* variants, the clustering criterion is the existence of a physical bond between beads on different chains. The physical-bond is pair-wise exclusive: one bead/sticker can only physically bond with one other bead. The maximal possible distance for a physical bond if $\sqrt{3}$. For the *Proximity* variants, the clustering criterion is based on a maximal bead-to-bead distance of $\sqrt{3}$. This corresponds to the $3^3 - 1 = 27 - 1 = 26$ lattice sites at most one lattice site away from a bead. For either variant, a translation of the resulting cluster is then attempted within a radius of $\frac{L}{2}$. Steric clash at the newly proposed location trivially results in move rejection.

For the *Large* variants the second-largest cluster is picked for the move. To obtain the second-largest cluster, the total cluster composition of the system is calculated. Suppose that $M_1 \geq M_2$ represent the largest and second-largest cluster sizes, respectively. If there is only one cluster in the system, $M_1 = M_2$, and that cluster if picked. If there are multiple clusters of size $M_2$, then one is randomly picked from that set. Thus, the move samples uniformly from the cluster-size-distribution of the system. For the *Small* variants, a random chain's cluster is picked. The cluster for a random is chain is computed. If the size of the cluster is larger than $\frac{N_{Tot}}{2}$ , where $N_{Tot}$ is the total number of chains in the system (2500 in this work), the move is rejected – otherwise, the cluster is picked. Therefore, these moves uniformly sample from the cluster-weight-distribution.

To ensure detailed balance, the clusters are recalculated at the new location. For the *Proximity* variants, if the new location results in a different cluster, the moves are rejected. For the *Anisotropic* variants, the internal structure of the cluster is unchanged and thus the moves are accepted if there is no steric clash.

*Simulation Protocol*: All simulations start with random initial conditions. For $t_{EQ} = 5 \times 10^7$ MC steps, the simulation temperature is $T_{EQ} = 5T^*$. A constraining potential is applied to the system: $V(\vec{r}, \Delta T) = \Delta T(\vec{r} - \vec{r}_{cen})^2$ where $\vec{r}_{cen} = \left\langle \frac{L}{2}, \frac{L}{2}, \frac{L}{2} \right\rangle$ is the center of the box, $\Delta T = T_{EQ} - T_1$. $T_1 = 1T^*$ corresponds to the first target temperature. This potential condenses all the chains to the center of the lattice. For Model B, the anisotropic interactions are turned off during this time.

After $t_{EQ}$ MC steps, anisotropic interactions are turned on, and the simulation temperature is exponentially annealed as such $T(t) = T_1 + T_{EQ} e^{-\frac{4t}{t_{EQ}}}$. Since $V(\vec{r}, \Delta T)$ depends on $\Delta T$, the annealing also results in the relaxation of the constraining potential. The simulations are run for $t_1 = 1.25 \times 10^{10}$ MC steps (first cycle) after which the simulation temperature is discontinuously increased to $T_2 = 2T^*$ and run for another $t_2 = 1.25 \times 10^{10}$ MC steps (second cycle).

Data are acquired in the last half of each cycle at a frequency of $f_{data} = 2.5 \times 10^6$ MC steps, resulting in 2500 samples for each condition per run. The average of those 2500 samples is reported at the end of each simulation. The standard error of the mean between replicates is used as a measure of uncertainty. Lastly, data plots are generated using Python packages NumPy, Matplotlib and Seaborn. Adobe Illustrator is used to make the figures.

**Section C. Amino Acid Sequences of Proteins used in Various Spectroscopic Studies**

**1. FUS-eGFP: This sequence includes the full-length FUS (unshaded), a linker that is cleavable by a TEV protease (shaded in yellow), and the eGFP (shaded in green).**

MASNDYTQQATQSYGAYPTQPGQGYSQQSSQPYGQQSYSGYSQSTDTSGYGQSSYSSYGQSQ
NTGYGTQSTPQGYGSTGGYGSSQSSQSSYGQQSSYPGYGQQPAPSSTSGSYGSSSQSSSYGQ
PQSGSYSQQPSYGGQQQSYGQQQSYNPPQGYGQQNQYNSSSGGGGGGGGGGNYGQDQSSMSS
GGGSGGGYGNQDQSGGGGSGGYGQQASDRGGRGRGGSGGGGGGGGGGYNRSSGGYEPRGRGG
GRGGRGGMGGSDRGGFNKFGGPRDQGSRHDSEQDNSDNNTIFVQGLGENVTIESVADYFKQI
GIIKTNKKTGQPMINLYTDRETGKKGEATVSFDDPPSAKAAIDWFDGKEFSGNPIKVSFATR
RADFNRGGGNGRGGRGRGMGRGGYGGGGSGGGGRGGFPSGGGGGGGQQRAGDWKCPNPTCEN
MNFSWRNECNQCKAPKPDGPGGGPGGSHMGGNYGDDRRGGRGGYDRGGYRGRGGDRGGFRGG
RGGGDRGGFGPGKMDSRGEHRQDRRERPYGAPGSSSGRENLYFQGMVSKGEELFTGVVPILV
ELDGDVNGHKFSVSGEGEGDATYGKLTLKFICTTGKLPVPWPTLVTTLTYGVQCFSRYPDHM
KQHDFFKSAMPEGYVQERTIFFKDDGNYKTRAEVKFEGDTLVNRIELKGIDFKEDGNILGHK
LEYNYNSHNVYIMADKQKNGIKVNFKIRHNIEDGSVQLADHYQQNTPIGDGPVLLPDNHYLS
TQSALSKDPNEKRDHMVLLEFVTAAGITLGMDELYK

**2. FUS-SNAP: This sequence includes the full-length FUS (unshaded), a linker that is cleavable by a TEV protease (shaded in yellow), and the SNAP (shaded in gray).**

MASNDYTQQATQSYGAYPTQPGQGYSQQSSQPYGQQSYSGYSQSTDTSGYGQSSYSSYGQSQ
NTGYGTQSTPQGYGSTGGYGSSQSSQSSYGQQSSYPGYGQQPAPSSTSGSYGSSSQSSSYGQ
PQSGSYSQQPSGGQQQSYGQQQSYNPPQGYGQQNQYNSSSGGGGGGGGGGNYGQDQSSMSSG
GGSGGGYGNQDQSGGGGSGGYGQQASDRGGRGRGGSGGGGGGGGGGYNRSSGGYEPRGRGGG
RGGRGGMGGSDRGGFNKFGGPRDQGSRHDSEQDNSDNNTIFVQGLGENVTIESVADYFKQIG
IIKTNKKTGQPMINLYTDRETGKLKGEATVSFDDPPSAKAAIDWFDGKEFSGNPIKVSFATR
RADFNRGGGNGRGGRGRGGPMGRGGYGGGGSGGGGRGGFPSGGGGGGGQQRAGDWKCPNPTC
ENMNFSWRNECNQCKAPKPDGPGGGPGGSHMGGNYGDDRRGGRGGYDRGGYRGRGGDRGGFR
GGRGGGDRGGFGPGKMDSRGEHRQDRRERPYGAPGSSSGRENLYFQGMDKDCEMKRTTLDSP
LGKLELSGCEQGLHRIIFLGKGTSAADAVEVPAPAAVLGGPEPLMQATAWLNAYFHQPEAIE
EFPVPALHHPVFQQESFTRQVLWKLLKVVKFGEVISYSHLAALAGNPAATAAVKTALSGNPV
PILIPCHRVVQGDLDVGGYEGGLAVKEWLLAHEGHRLGKPGLG

**3. FUS**

MASNDYTQQATQSYGAYPTQPGQGYSQQSSQPYGQQSYSGYSQSTDTSGYGQSSYSSYGQSQ
NTGYGTQSTPQGYGSTGGYGSSQSSQSSYGQQSSYPGYGQQPAPSSTSGSYGSSSQSSSYGQ
PQSGSYSQQPSYGGQQQSYGQQQSYNPPQGYGQQNQYNSSSGGGGGGGGGGNYGQDQSSMSS
GGGSGGGYGNQDQSGGGGSGGYGQQASDRGGRGRGGSGGGGGGGGGGYNRSSGGYEPRGRGG
GRGGRGGMGGSDRGGFNKFGGPRDQGSRHDSEQDNSDNNTIFVQGLGENVTIESVADYFKQI

GIIKTNKKTGQPMINLYTDRETGKLKGEATVSFDDPPSAKAAIDWFDGKEFSGNPIKVSFAT
RRADFNRGGGNGRGGRGRGGPMGRGGYGGGGSGGGGRGGFPSGGGGGGGQQRAGDWKCPNPT
CENMNFSWRNECNQCKAPKPDGPGGGPGGSHMGGNYGDDRRGGRGGYDRGGYRGRGGDRGGF
RGGRGGGDRGGFGPGKMDSRGEHRQDRRERPY

**4. PLD of FUS (1-211)**

MASNDYTQQATQSYGAYPTQPGQGYSQQSSQPYGQQSYSGYSQSTDTSGYGQSSYSSYGQSQ
NTGYGTQSTPQGYGSTGGYGSSQSSQSSYGQQSSYPGYGQQPAPSSTSGSYGSSSQSSSYGQ
PQSGSYSQQPSYGGQQQSYGQQQSYNPPQGYGQQNQYNSSSGGGGGGGGGGNYGQDQSSMSS
GGGSGGGYGNQDQSGGGGSGGYGQQ

**5. RBD of FUS (212-526)**

DRGGRGRGGSGGGGGGGGGGYNRSSGGYEPRGRGGGRGGRGGMGGSDRGGFNKFGGPRDQGS
RHDSEQDNSDNNTIFVQGLGENVTIESVADYFKQIGIIKTNKKTGQPMINLYTDRETGKLKG
EATVSFDDPPSAKAAIDWFDGKEFSGNPIKVSFATRRADFNRGGGNGRGGRGRGGPMGRGGY
GGGGSGGGGRGGFPSGGGGGGGQQRAGDWKCPNPTCENMNFSWRNECNQCKAPKPDGPGGGP
GGSHMGGNYGDDRRGGRGGYDRGGYRGRGGDRGGFRGGRGGGDRGGFGPGKMDSRGEHRQDR
RERPY

**6. Full-length FUS with 24 Arg in its RBD substituted to Gly (FUS-24-R-to-G)**

MASNDYTQQATQSYGAYPTQPGQGYSQQSSQPYGQQSYSGYSQSTDTSGYGQSSYSSYGQSQ
NTGYGTQSTPQGYGSTGGYGSSQSSQSSYGQQSSYPGYGQQPAPSSTSGSYGSSSQSSSYGQ
PQSGSYSQQPSYGGQQQSYGQQQSYNPPQGYGQQNQYNSSSGGGGGGGGGGNYGQDQSSMSS
GGGSGGGYGNQDQSGGGGSGGYGQQASDGGGGGGGGSGGGGGGGGGGYNRSSGGYEPGGGGG
GGGGGGGMGGSDGGGFNKFGGPRDQGSRHDSEQDNSDNNTIFVQGLGENVTIESVADYFKQI
GIIKTNKKTGQPMINLYTDRETGKLKGEATVSFDDPPSAKAAIDWFDGKEFSGNPIKVSFAT
RRADFNGGGGNGGGGGGGGGPMGGGGYGGGGSGGGGGGGFPSGGGGGGGQQRAGDWKCPNPT
CENMNFSWRNECNQCKAPKPDGPGGGPGGSHMGGNYGDDRGGGGGGYDGGGYGGGGGDGGGF
GGGGGGGDGGGFGPGKMDSGGEHRQDRRERPY

**7. Full-length FUS with 10 Asp and 4 Glu residues in the RBD substituted to Gly (FUS-10D/4E-G)**

MASNDYTQQATQSYGAYPTQPGQGYSQQSSQPYGQQSYSGYSQSTDTSGYGQSSYSSYGQSQ
NTGYGTQSTPQGYGSTGGYGSSQSSQSSYGQQSSYPGYGQQPAPSSTSGSYGSSSQSSSYGQ
PQSGSYSQQPSYGGQQQSYGQQQSYNPPQGYGQQNQYNSSSGGGGGGGGGGNYGQDQSSMSS
GGGSGGGYGNQDQSGGGGSGGYGQQASDRGGRGRGGSGGGGGGGGGGYNRSSGGYEPRGRGG
GRGGRGGMGGSDRGGFNKFGGPRDQGSRHDSEQDNSDNNTIFVQGLGENVTIESVADYFKQI
GIIKTNKKTGQPMINLYTDRETGKLKGEATVSFDDPPSAKAAIDWFDGKEFSGNPIKVSFAT
RRAGFNRGGGNGRGGRGRGGPMGRGGYGGGGSGGGGRGGFPSGGGGGGGQQRAGGWKCPNPT
CGNMNFSWRNGCNQCKAPKPGGPGGGPGGSHMGGNYGGGRRGGRGGYGRGGYRGRGGGRGGF
RGGRGGGGRGGFGPGKMGSRGGHRQGRRGRPY

**8. Full-length FUS with 10 Tyr residues in the PLD substituted to Ser (FUS-10Y-to-S)**

MASNDYTQQATQSYGASPTQPGQGYSQQSSQPSGQQSYSGSSQSTDTSGSGQSSYSSSGQSQ
NTGYGTQSTPQGSGSTGGYGSSQSSQSSYGQQSSSPGYGQQPAPSSTSGSYGSSSQSSSYGQ
PQSGSSSQQPSYGGQQQSSGQQQSSNPPQGYGQQNQYNSSSGGGGGGGGGGNYGQDQSSMSS

GGGSGGGYGNQDQSGGGGSGGYGQQASDRGGRGRGGSGGGGGGGGGGYNRSSGGYEPRGRGG
GRGGRGGMGGSDRGGFNKFGGPRDQGSRHDSEQDNSDNNTIFVQGLGENVTIESVADYFKQI
GIIKTNKKTGQPMINLYTDRETGKLKGEATVSFDDPPSAKAAIDWFDGKEFSGNPIKVSFAT
RRADFNRGGGNGRGGRGRGGPMGRGGYGGGGSGGGGRGGFPSGGGGGGGQQRAGDWKCPNPT
CENMNFSWRNECNQCKAPKPDGPGGGPGGSHMGGNYGDDRRGGRGGYDRGGYRGRGGDRGGF
RGGRGGGDRGGFGPGKMDSRGEHRQDRRERPY

**9. Full-length FUS with 18 Tyr residues in the PLD substituted to Ser (FUS-18Y-to-S)**
MASNDSTQQATQSYGASPTQPGQGSSQQSSQPSGQQSYSGSSQSTDTSGSGQSSYSSSGQSQ
NTGSGTQSTPQGSGSTGGSGSSQSSQSSYGQQSSSPGYGQQPAPSSTSGSSGSSSQSSSSGQ
PQSGSSSQQPSYGGQQQSSGQQQSSNPPQGYGQQNQSNSSSGGGGGGGGGNSGQDQSSMSS
GGGSGGGYGNQDQSGGGGSGGYGQQASDRGGRGRGGSGGGGGGGGGGYNRSSGGYEPRGRGG
GRGGRGGMGGSDRGGFNKFGGPRDQGSRHDSEQDNSDNNTIFVQGLGENVTIESVADYFKQI
GIIKTNKKTGQPMINLYTDRETGKLKGEATVSFDDPPSAKAAIDWFDGKEFSGNPIKVSFAT
RRADFNRGGGNGRGGRGRGGPMGRGGYGGGGSGGGGRGGFPSGGGGGGGQQRAGDWKCPNPT
CENMNFSWRNECNQCKAPKPDGPGGGPGGSHMGGNYGDDRRGGRGGYDRGGYRGRGGDRGGF
RGGRGGGDRGGFGPGKMDSRGEHRQDRRERPY

**10. Full-length FUS with 27 Tyr residues in the PLD substituted to Ser (FUS-27Y-to-S)**
MASNDSTQQATQSSGASPTQPGQGSSQQSSQPSGQQSSSGSSQSTDTSGSGQSSSSSSGQSQ
NTGSGTQSTPQGSGSTGGSGSSQSSQSSSGQQSSSPGSGQQPAPSSTSGSSGSSSQSSSSGQ
PQSGSSSQQPSSGGQQQSSGQQQSSNPPQGSGQQNQSNSSSGGGGGGGGGNSGQDQSSMSS
GGGSGGGSGNQDQSGGGGSGGSGQQASDRGGRGRGGSGGGGGGGGGGYNRSSGGYEPRGRGG
GRGGRGGMGGSDRGGFNKFGGPRDQGSRHDSEQDNSDNNTIFVQGLGENVTIESVADYFKQI
GIIKTNKKTGQPMINLYTDRETGKLKGEATVSFDDPPSAKAAIDWFDGKEFSGNPIKVSFAT
RRADFNRGGGNGRGGRGRGGPMGRGGYGGGGSGGGGRGGFPSGGGGGGGQQRAGDWKCPNPT
CENMNFSWRNECNQCKAPKPDGPGGGPGGSHMGGNYGDDRRGGRGGYDRGGYRGRGGDRGGF
RGGRGGGDRGGFGPGKMDSRGEHRQDRRERPY

**11. Full-length FUS with 6 Phe residues in the RBD substituted to Gly (FUS-6F-to-G)**
AMASNDYTQQATQSYGAYPTQPGQGYSQQSSQPYGQQSYSGYSQSTDTSGYGQSSYSSYGQS
QNTGYGTQSTPQGYGSTGGYGSSQSSQSSYGQQSSYPGYGQQPAPSSTSGSYGSSSQSSSYG
QPQSGSYSQQPSYGGQQQSYGQQQSYNPPQGYGQQNQYNSSSGGGGGGGGGNYGQDQSSMS
SGGGSGGGYGNQDQSGGGGSGGYGQQASDRGGRGRGGSGGGGGGGGGGYNRSSGGYEPRGRG
GGRGGRGGMGGSDRGGGNKGGGPRDQGSRHDSEQDNSDNNTIFVQGLGENVTIESVADYFKQ
IGIIKTNKKTGQPMINLYTDRETGKLKGEATVSFDDPPSAKAAIDWFDGKEFSGNPIKVSFA
TRRADGNRGGGNGRGGRGRGGPMGRGGYGGGGSGGGGRGGGPSGGGGGGGQQRAGDWKCPNP
TCENMNFSWRNECNQCKAPKPDGPGGGPGGSHMGGNYGDDRRGGRGGYDRGGYRGRGGDRGG
GRGGRGGGDRGGGGPGKMDSRGEHRQDRRERPY

**12. Full-length FUS with 6 Tyr residues in the RBD substituted to Ser (FUS-6Y-to-S)**
AMASNDYTQQATQSYGAYPTQPGQGYSQQSSQPYGQQSYSGYSQSTDTSGYGQSSYSSYGQS
QNTGYGTQSTPQGYGSTGGYGSSQSSQSSYGQQSSYPGYGQQPAPSSTSGSYGSSSQSSSYG
QPQSGSYSQQPSYGGQQQSYGQQQSYNPPQGYGQQNQYNSSSGGGGGGGGGNYGQDQSSMS
SGGGSGGGYGNQDQSGGGGSGGYGQQASDRGGRGRGGSGGGGGGGGGGSNRSSGGSEPRGRG
GGRGGRGGMGGSDRGGFNKFGGPRDQGSRHDSEQDNSDNNTIFVQGLGENVTIESVADYFKQ

IGIIKTNKKTGQPMINLYTDRETGKLKGEATVSFDDPPSAKAAIDWFDGKEFSGNPIKVSFA
TRRADFNRGGGNGRGGRGRGGPMGRGGSGGGGSGGGGRGGFPSGGGGGGGQQRAGDWKCPNP
TCENMNFSWRNECNQCKAPKPDGPGGGPGGSHMGGNSGDDRRGGRGGSDRGGSRGRGGDRGG
FRGGRGGGDRGGFGPGKMDSRGEHRQDRRERPY

**13. TAF15-SNAP: This sequence includes the full-length TAF15 (unshaded), a linker that is cleavable by a TEV protease (shaded in yellow), and the SNAP tag (shaded in gray).**
MSDSGSYGQSGGEQQSYSTYGNPGSQGYGQASQSYSGYGQTTDSSYGQNYSGYSSYGQSYSQ
SYGGYENQKQSSYSQQPYNNQGQQQNMESSGSQGGRAPSYDQPDYGQQDSYDQQSGYDQHQG
SYDEQSNYDQQHDSYSQNQQSYHSQRENYSHHTQDDRRDVSRYGEDNRGYGGSQGGGRGRGG
YDKDGRGPMTGSSGGDRGGFKNFGGHRDYGPRTDADSESDNSDNNTIFVQGLGEGVSTDQVG
EFFKQIGIIKTNKKTGKPMINLYTDKDTGKPKGEATVSFDDPPSAKAAIDWFDGKEFHGNII
KVSFATRRPEFMRGGGSGGRRGRGGYRGRGGFQGRGGDPKSGDWVCPNPSCGNMNFARRNS
CNQCNEPRPEDSRPSGGDFRGRGYGGERGYRGRGGRGGDRGGYGGDRSGGGYGGDRSSGGGY
SGDRSGGGYGGDRSGGGYGGDRGGGYGGDRGGGYGGDRGGGYGGDRGGYGGDRGGGYGGDRG
GYGGDRGGYGGDRGGYGGDRGGYGGDRSRGGYGGDRGGGSGYGGDRSGGYGGDRSGGGYGGD
RGGGYGGDRGGYGGKMGGRNDYRNDQRNRPYGAPGSSSGRENLYFQGMDKDCEMKRTTLDSP
LGKLELSGCEQGLHRIIFLGKGTSAADAVEVPAPAAVLGGPEPLMQATAWLNAYFHQPEAIE
EFPVPALHHPVFQQESFTRQVLWKLLKVVKFGEVISYSHLAALAGNPAATAAVKTALSGNPV
PILIPCHRVVQGDLDVGGYEGGLAVKEWLLAHEGHRLGKPGLG

**14. EWSR1-SNAP: This sequence includes the full-length EWSR1 (unshaded), a linker that is cleavable by a TEV protease (shaded in yellow), and the SNAP tag (shaded in gray).**
MASTDYSTYSQAAAQQGYSAYTAQPTQGYAQTTQAYGQQSYGTYGQPTDVSYTQAQTTATYG
QTAYATSYGQPPTGYTTPTAPQAYSQPVQGYGTGAYDTTTATVTTTQASYAAQSAYGTQPAY
PAYGQQPAATAPTRPQDGNKPTETSQPQSSTGGYNQPSLGYGQSNYSYPQVPGSYPMQPVTA
PPSYPPTSYSSTQPTSYDQSSYSQQNTYGQPSSYGQQSSYGQQSSYGQQPPTSYPPQTGSYS
QAPSQYSQQSSSYGQQSSFRQDHPSSMGVYGQESGGFSGPGENRSMSGPDNRGRGRGGFDRG
GMSRGGRGGGRGGMGSAGERGGFNKPGGPMDEGPDLDLGPPVDPDEDSDNSAIYVQGLNDSV
TLDDLADFFKQCGVVKMNKRTGQPMIHIYLDKETGKPKGDATVSYEDPPTAKAAVEWFDGKD
FQGSKLKVSLARKKPPMNSMRGGLPPREGRGMPPPLRGGPGGPGGPGGPMGRMGGRGGDRGG
FPPRGPRGSRGNPSGGGNVQHRAGDWQCPNPGCGNQNFAWRTECNQCKAPKPEGFLPPPFPP
PGGDRGRGGPGGMRGGRGGLMDRGGPGGMFRGGRGGDRGGFRGGRGMDRGGFGGGRRGGPGG
PPGPLMEQMGGRRGGRGGPGKMDKGEHRQERRDRPYGAPGSSSGRENLYFQGMDKDCEMKRT
TLDSPLGKLELSGCEQGLHRIIFLGKGTSAADAVEVPAPAAVLGGPEPLMQATAWLNAYFHQ
PEAIEEFPVPALHHPVFQQESFTRQVLWKLLKVVKFGEVISYSHLAALAGNPAATAAVKTAL
SGNPVPILIPCHRVVQGDLDVGGYEGGLAVKEWLLAHEGHRLGKPGLG

**15. hnRNPA3**
MEVKPPPGRPQPDSGRRRRRRGEEGHDPKEPEQLRKLFIGGLSFETTDDSLREHFEKWGTLT
DCVVMRDPQTKRSRGFGFVTYSCVEEVDAAMCARPHKVDGRVVEPKRAVSREDSVKPGAHLT
VKKIFVGGIKEDTEEYNLRDYFEKYGKIETIEVMEDRQSGKKRGFAFVTFDDHDTVDKIVVQ
KYHTINGHNCEVKKALSKQEMQSAGSQRGRGGGSGNFMGRGGNFGGGGGNFGRGGNFGGRGG
YGGGGGGSRGSYGGGDGGYNGFGGDGGNYGGGPGYSSRGGYGGGGPGYGNQGGGYGGGGGYD
GYNEGGNFGGGNYGGGGNYNDFGNYSGQQQSNYGPMKGGSFGGRSSGSPYGGGYGSGGGSGG
YGSRRFGAP

**16. TDP 43-eGFP: This sequence includes the full-length TDP-43 (unshaded), a linker that is cleavable by a TEV protease (shaded in yellow), and the eGFP (shaded in green).**

MSEYIRVTEDENDEPIEIPSEDDGTVLLSTVTAQFPGACGLRYRNPVSQCMRGVRLVEGILH
APDAGWGNLVYVVNYPKDNKRKMDETDASSAVKVKRAVQKTSDLIVLGLPWKTTEQDLKEYF
STFGEVLMVQVKKDLKTGHSKGFGFVRFTEYETQVKVMSQRHMIDGRWCDCKLPNSKQSQDE
PLRSRKVFVGRCTEDMTEDELREFFSQYGDVMDVFIPKPFRAFAFVTFADDQIAQSLCGEDL
IIKGISVHISNAEPKHNSNRQLERSGRFGGNPGGFGNQGGFGNSRGGGAGLGNNQGSNMGGG
MNFGAFSINPAMMAAAQAALQSSWGMMGMLASQQNQSGPSGNNQNQGNMQREPNQAFGSGNN
SYSGSNSGAAIGWGSASNAGSGSGFNGGFGSSMDSKSSGWGMGAPGSSSGRENLYFQGMVSK
GEELFTGVVPILVELDGDVNGHKFSVSGEGEGDATYGKLTLKFICTTGKLPVPWPTLVTTLT
YGVQCFSRYPDHMKQHDFFKSAMPEGYVQERTIFFKDDGNYKTRAEVKFEGDTLVNRIELKG
IDFKEDGNILGHKLEYNYNSHNVYIMADKQKNGIKVNFKIRHNIEDGSVQLADHYQQNTPIG
DGPVLLPDNHYLSTQSALSKDPNEKRDHMVLLEFVTAAGITLGMDELYK

### Section D: Compendium of supplemental figures

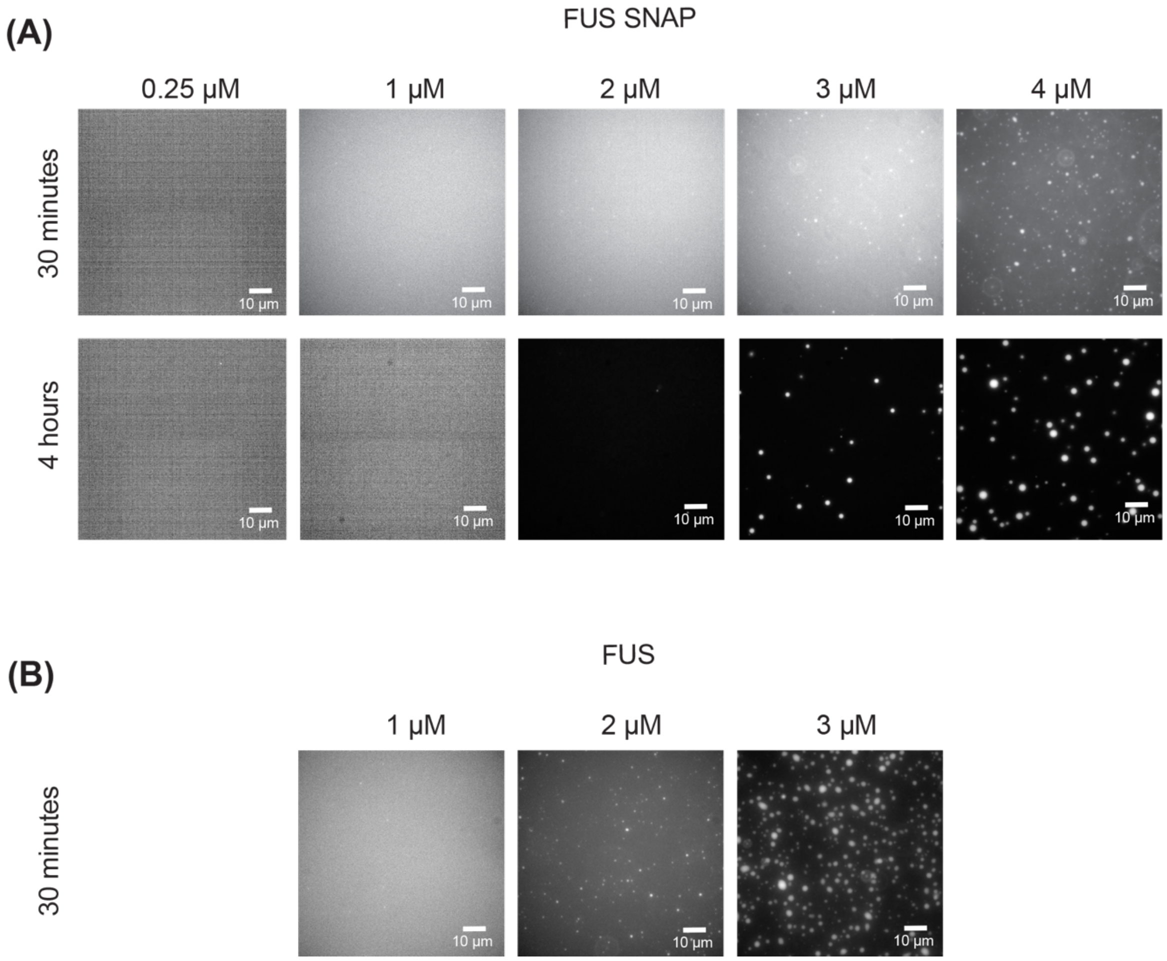


**Fig. S1: Microscopy-based verifications of the existence of $c_{sat}$ for FUS molecules.** (A) The top row shows microscopy images collected 30 minutes after sample preparation for different bulk concentrations of FUS-SNAP. In these experiments, 5% of the molecules are FUS-eGFP. Micron-scale species are only visible for concentrations above $c_{sat}$. The bottom row, which shows images collected after 4-hours corroborates the veracity of $c_{sat}$ that was estimated using an absorbance assay as described in the main text. (B) Images collected for untagged FUS, 30 minutes after sample preparation. The images, collected at three different concentrations corroborate the veracity of the estimate for $c_{sat} \approx 2$ μM for untagged FUS. All samples were prepared in 20 mM Tris, pH 7.4, 100 mM KCl at ≈ 25˚C.

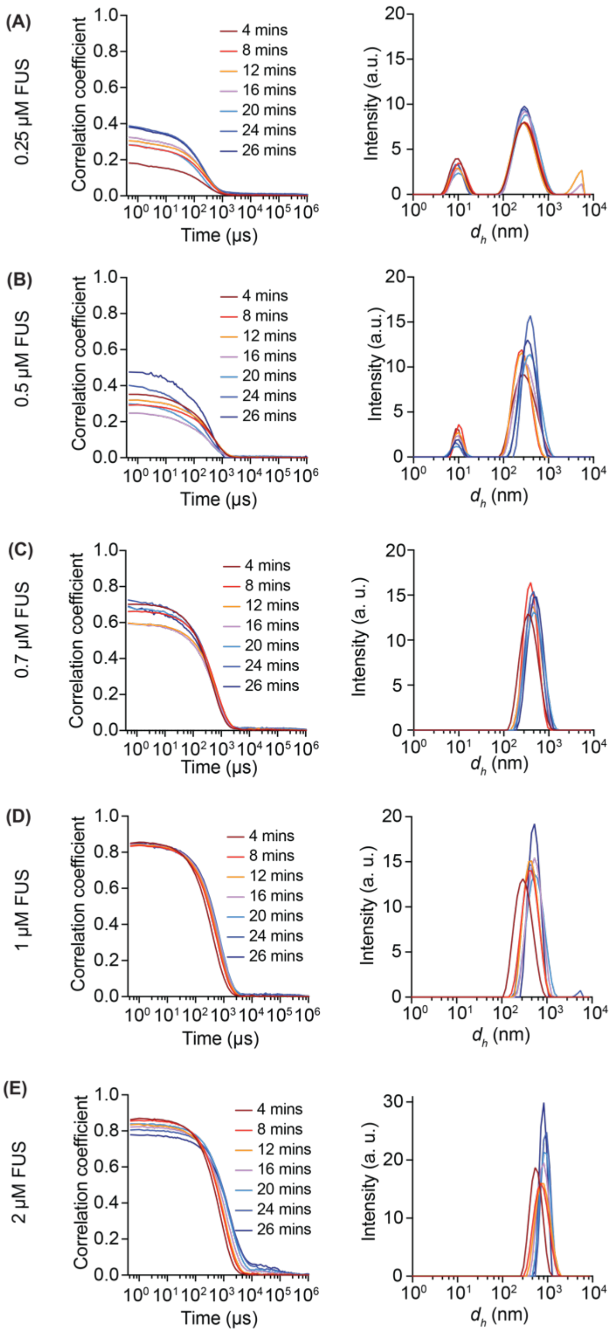


**Fig. S2: Raw DLS data for untagged FUS in subsaturated solutions.** Each panel has two columns. The column on the left shows the autocorrelation functions collected at different time points after sample preparation and the start of the data collection. The amplitudes of autocorrelation functions are indicators of the sizes of scatterers in solution. The timescale for decay of the autocorrelation function is useful for assessing the apparent hydrodynamic diameters of scatterers. Autocorrelation functions can be converted to intensity profiles. These profiles, shown in the right column, are typically dominated by the largest species in solution. Each row corresponds to a distinct bulk concentration as marked on the left. All samples were prepared in 20 mM Tris, pH 7.4, 100 mM KCl at ≈ 25˚C.

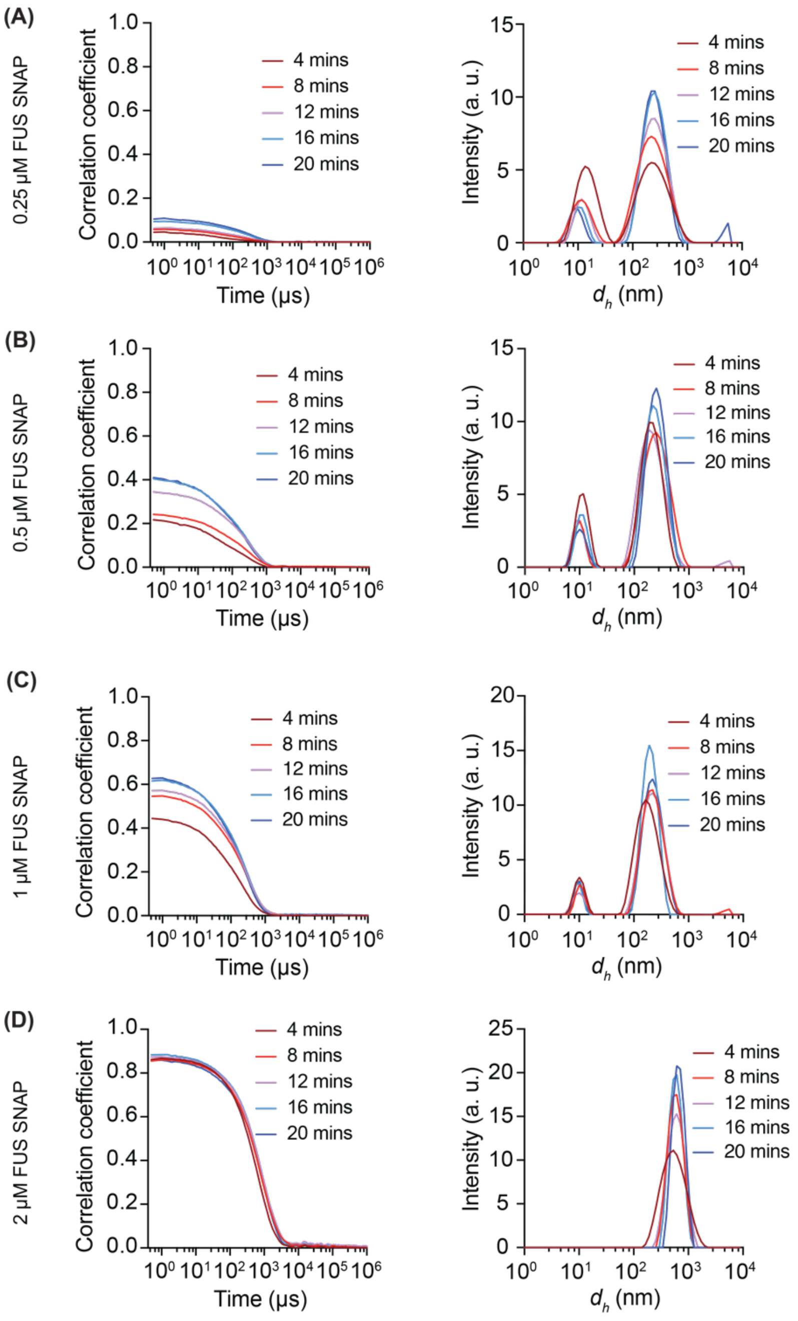


**Fig. S3: Raw DLS data for FUS-SNAP ($c_{sat}$ ≈ 3 µM) in subsaturated solutions.** The FUS-SNAP molecules used in these experiments were prepared using method A. As in Fig. S2, each panel has two columns. The column on the left shows the autocorrelation functions collected at different time points after sample preparation and the start of the data collection. The amplitudes of autocorrelation functions are indicators of the sizes of scatterers in solution. The timescale for decay of the autocorrelation function is useful for assessing the apparent hydrodynamic diameters of scatterers. The autocorrelation functions, converted to intensity profiles, are in the right column. Each row corresponds to a distinct bulk concentration of FUS-SNAP as marked on the left. All samples were prepared in 20 mM Tris, pH 7.4, 100 mM KCl at ≈ 25˚C.

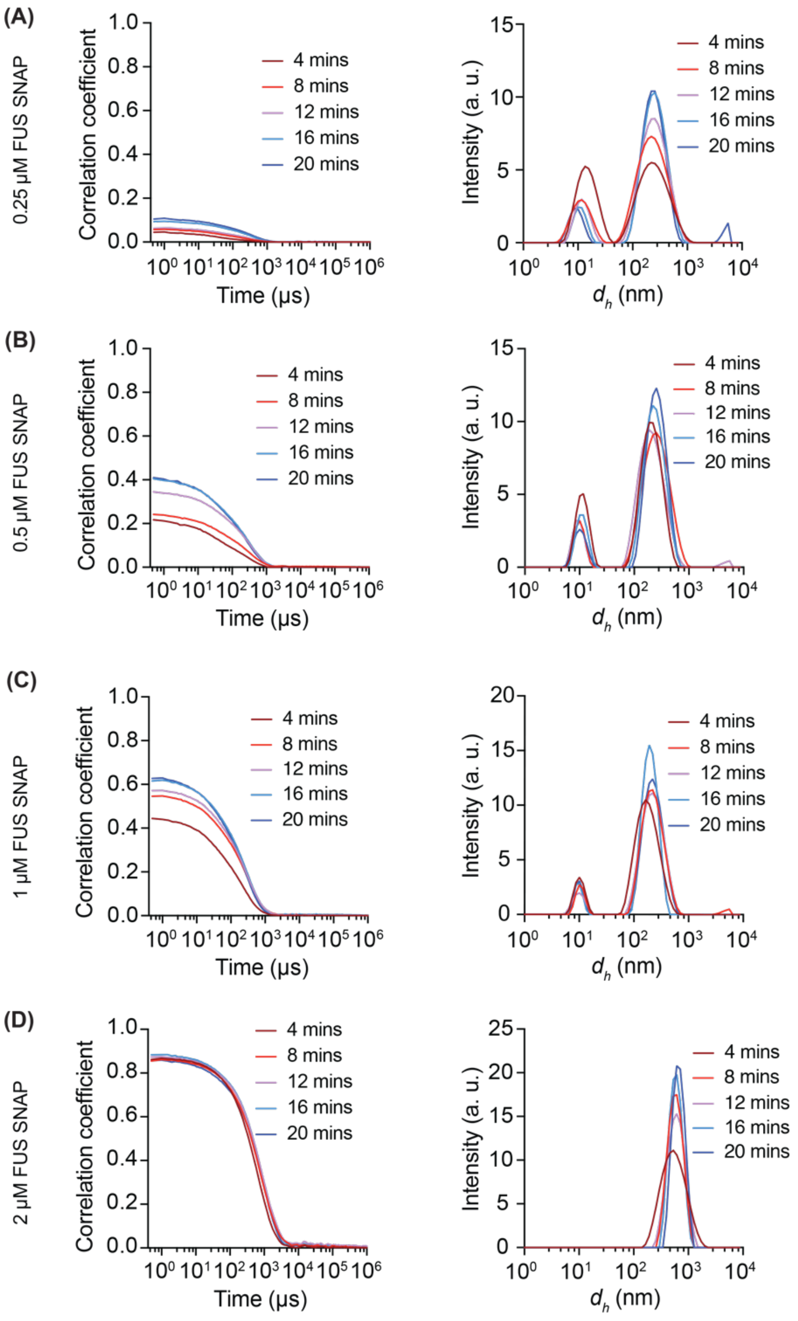


**Fig. S4: Raw DLS data for FUS-SNAP in subsaturated solutions.** The FUS-SNAP molecules used in these experiments were prepared using method B. Comparison of data shown here to the data shown in Fig. S3 indicate that the presence of DLS detectable mesoscale clusters, and their concentration dependence, are robust to the methods used to prepare FUS-SNAP samples. Method B includes a step wherein the samples are treated with benzonase, a generic nuclease. This should degrade the presence of any trace RNA contaminants that might be present in the sample. The data with and without this benzonase treatment step are equivalent to one another. This suggests that the properties of subsaturated solutions that we interrogate in our experiments are due to self-associations of FUS molecules. All samples were prepared in 20 mM Tris, pH 7.4, 100 mM KCl at ≈ 25˚C.

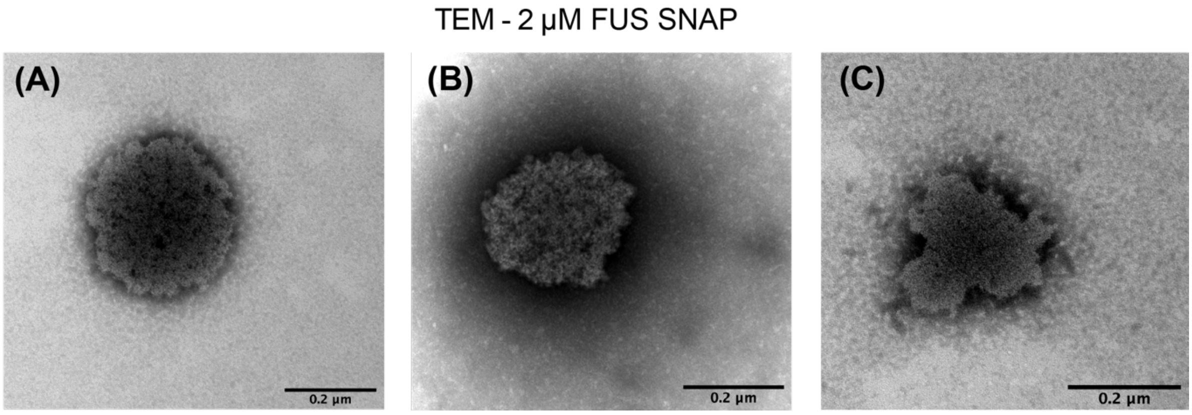


**Fig. S5: Additional representative TEM images for FUS-SNAP.** The images shown in panels A, B, and C show a roughly spherical morphology albeit with undulations that are visible on the length scales being interrogated. All samples were prepared in 20 mM Tris, pH 7.4, 100 mM KCl at ≈ 25˚C.

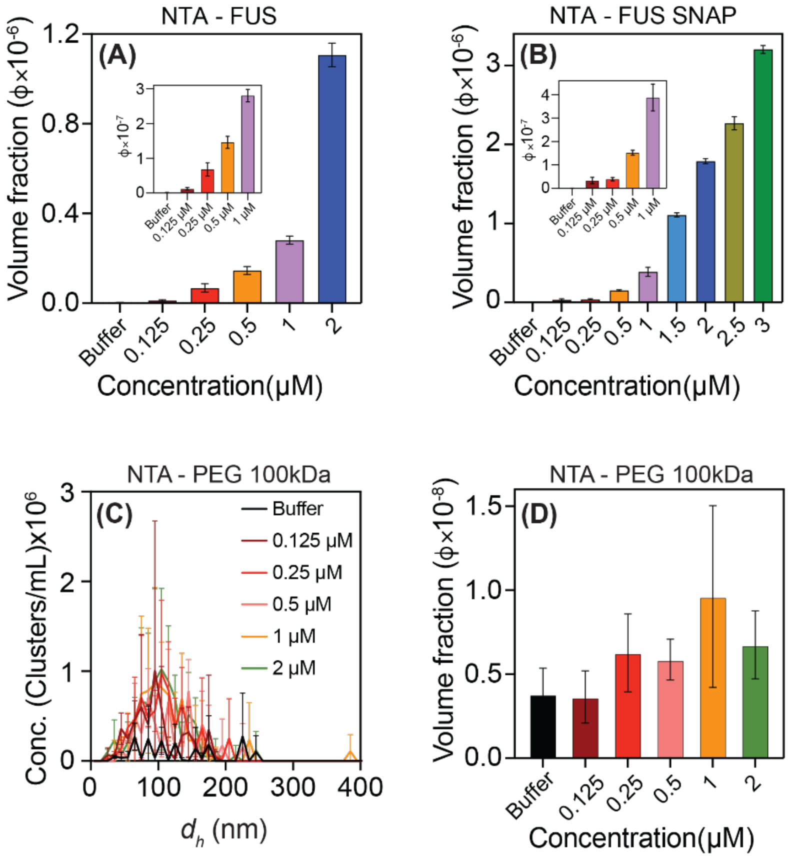


**Fig. S6: Concentration dependence of the abundance of mesoscale clusters formed by FUS and FUS-SNAP molecules as measured using NTA.** Panels (A) and (B) show data for untagged FUS and FUS-SNAP, respectively. The samples were prepared using method A. To calibrate expectations, we also performed NTA measurements for polyethylene glycol (PEG 100 kDa). PEG is a water-soluble polymer. As such, it should show negligible self-associations as a function of increased concentration. This is confirmed by comparing the distribution of detectable scatterers measured by NTA for PEG-100 kDa obtained at different polymer concentrations. Within error, these traces are only minutely different from that of the buffer alone. The low volume fraction of scatterers, quantified in panel (D) is consistent with the predominantly monomeric nature of PEG-100 kDa. All samples were prepared in 20 mM Tris, pH 7.4, 100 mM KCl at ≈ 25˚C.

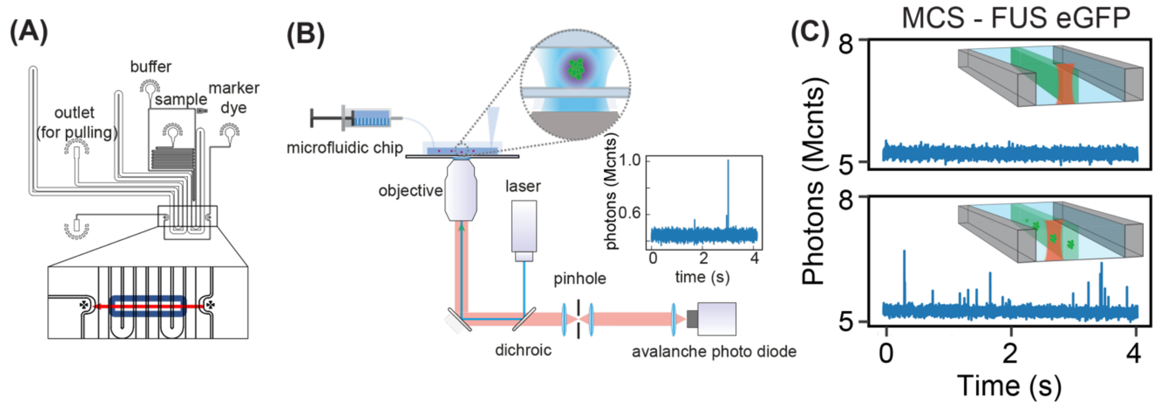


**Fig. S7: Working principles of microfluidic confocal detection (MCD). (A)** Schematic of microfluidic device used in MCD experiments. The insert and red line indicate the actual location within the channel which was scanned by the confocal spot. **(B)** Diagram of the working principle of the experiment and the optical setup. A 488 nm laser was used to excite the GFP-tagged fusion protein FUS. Emitted photons were counted on an avalanche photo diode, yielding raw traces as shown in the inserted plot. **(C)** When scanning the channel, 4 seconds of signal were acquired at each of 200 locations, evenly spaced through the channel. Clusters were quantified as signal peaks which exceeded 5 standard deviations above the mean signal for a given trace. **(D)** Clusters were quantified according to their position within the central four channels of the device (top panel, channels indicated by the dashed rectangles) and grouped according to the peaks in the mean intensity (bottom panel). The average of the total clusters in each of the channels was used for subsequent cluster concentration calculations.

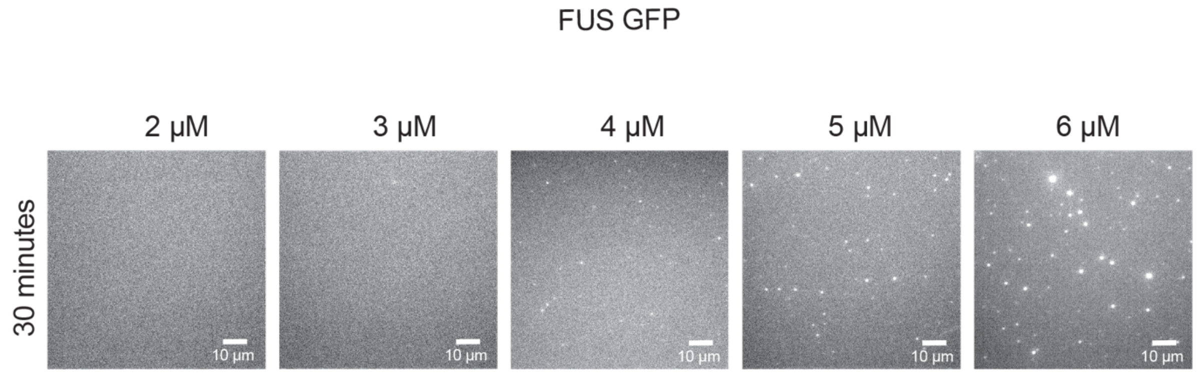


**Fig. S8: Visual confirmation that $c_{sat}$ for FUS-eGFP prepared using method A is ≈ 4 μM.** Microscopy images are shown here for different bulk concentrations of FUS-eGFP collected 30-minutes after sample preparation in 20 mM Tris, pH 7.4, 100 mM KCl at ≈ 25˚C

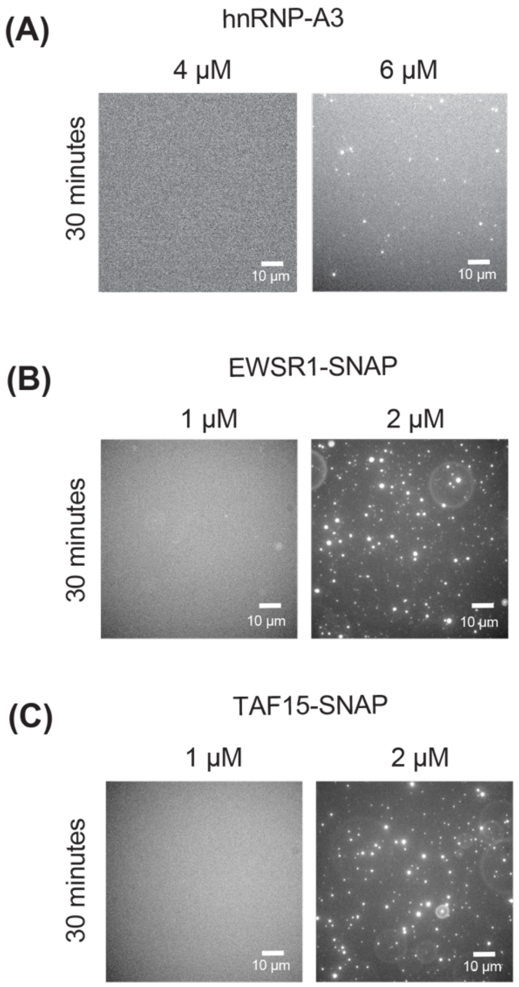


**Fig. S9: Saturation concentrations were measured for four different proteins in 20 mM Tris, pH 7.4, 100 mM KCl at ≈ 25˚C.** The measured $c_{sat}$ values were visually confirmed using microscopy for (A) hnRNP-A3 ($c_{sat} \approx 6$ µM), (B) EWSR1-SNAP ($c_{sat} \approx 2$ µM), and (C) TAF15-SNAP ($c_{sat} \approx 2$ µM).

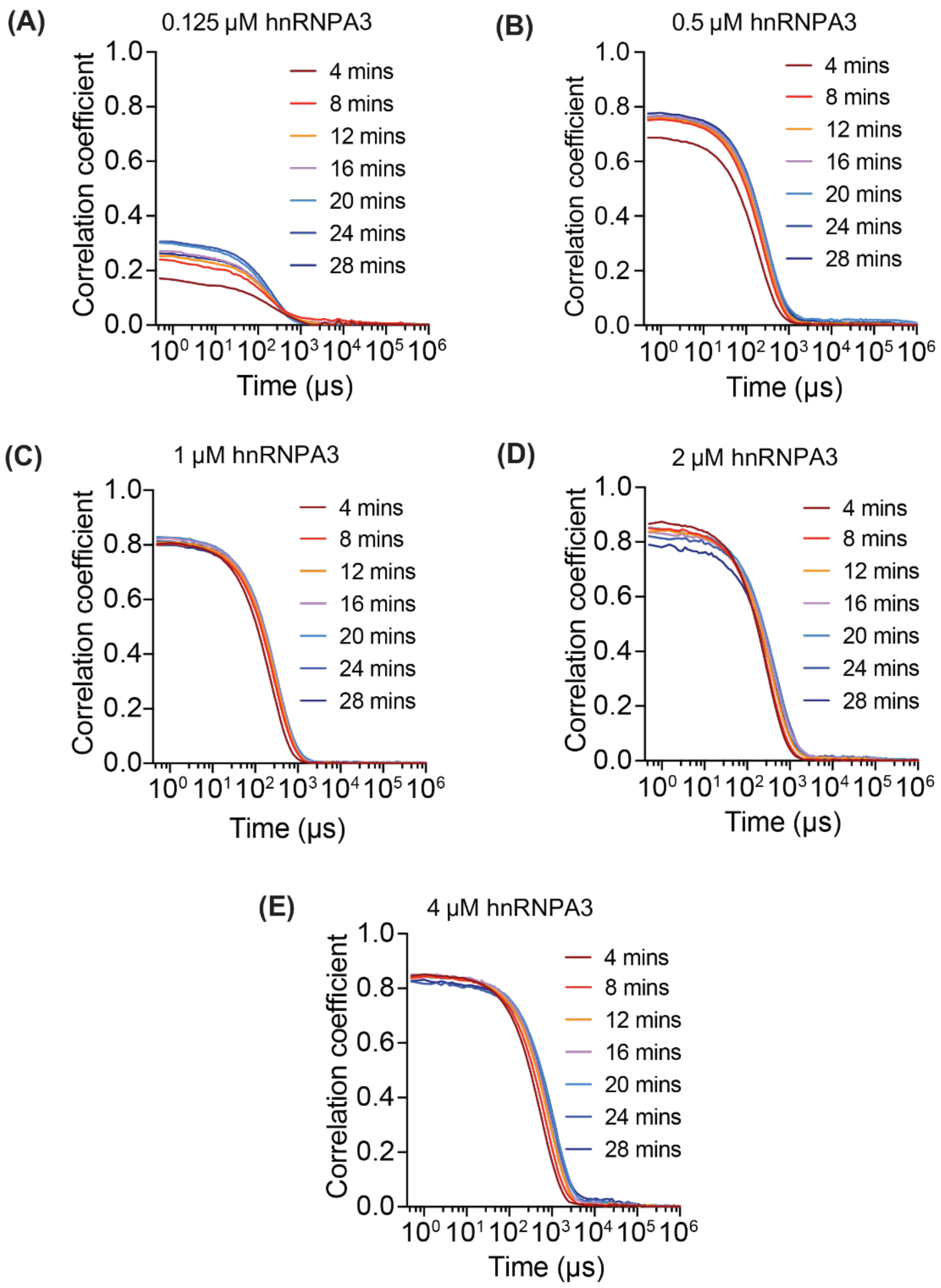


**Fig. S10: Raw DLS data for subsaturated solutions of hnRNPA3 ($c_{sat}$ ≈ 6 μM).** The data are shown as autocorrelation functions collected at different time points for different bulk concentrations representing different degrees of subsaturation. The amplitudes of the autocorrelation function are governed by the scattering intensities and the decay times are governed by the sizes of scatterers. The data clearly show that the sizes of mesoscale clusters increase with increasing concentration. And data were collected in 20 mM Tris, pH 7.4, 100 mM KCl, at ≈ 25˚C.

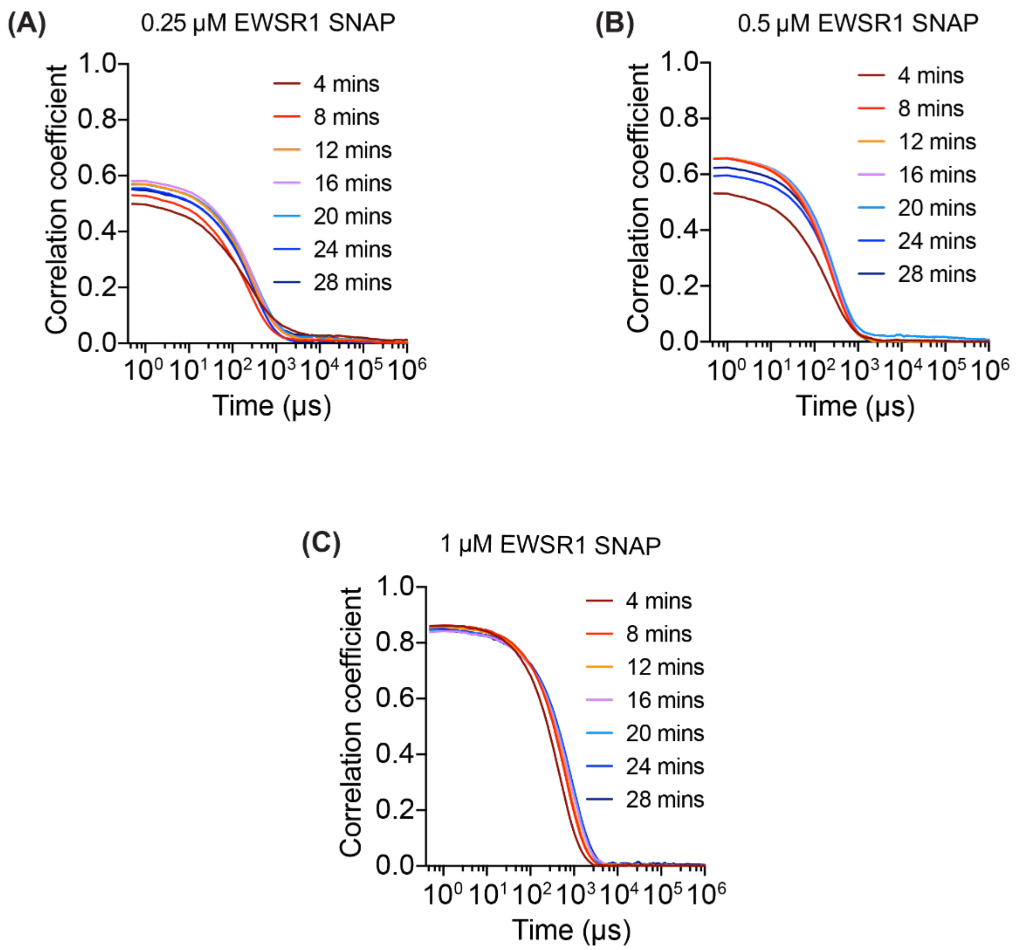


**Fig. S11: Raw DLS data for subsaturated solutions of EWSR1-SNAP ($c_{sat}$ ≈ 2 µM).** The data are shown as autocorrelation functions collected at different time points for different bulk concentrations representing different degrees of subsaturation. The amplitudes of the autocorrelation function are governed by the scattering intensities and the decay times are governed by the sizes of scatterers. The data clearly show that the sizes of mesoscale clusters increase with increasing concentration. Samples were prepared using method A. And data were collected in 20 mM Tris, pH 7.4, 100 mM KCl, at ≈ 25˚C.

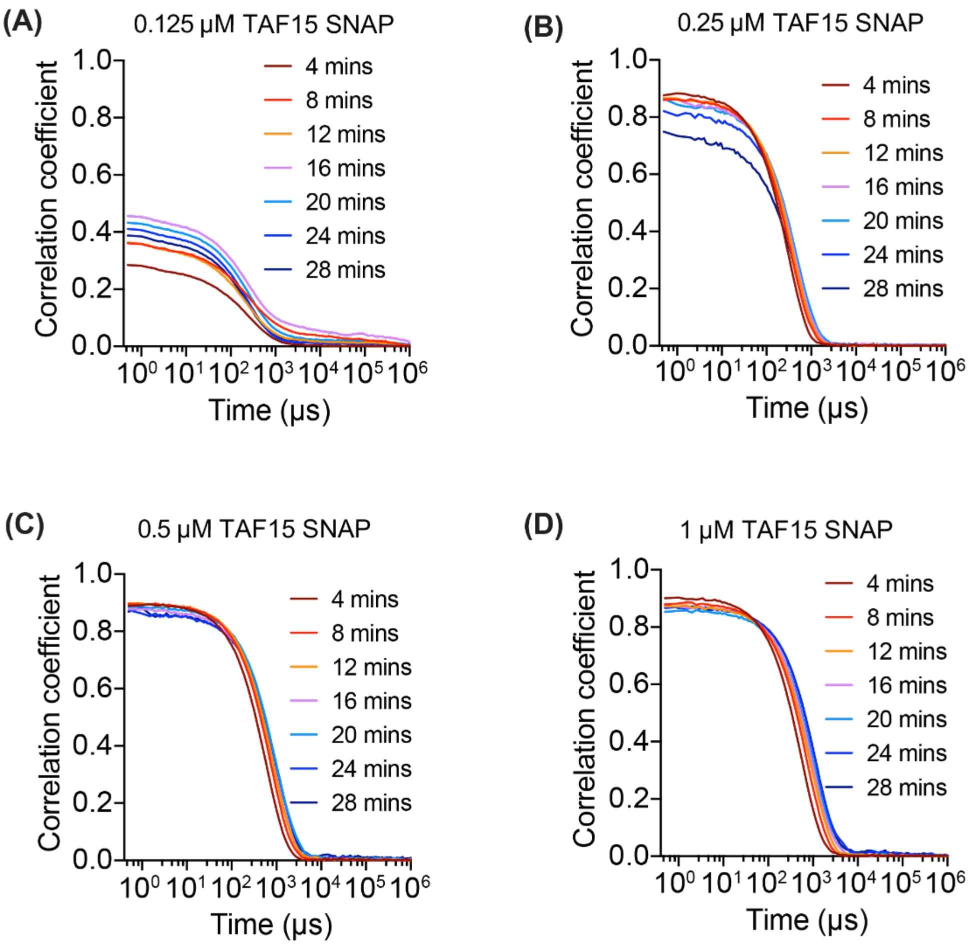


**Fig. S12: Raw DLS data for subsaturated solutions of TAF15-SNAP ($c_{sat} \approx 2$ µM).** The data are shown as autocorrelation functions collected at different time points for different bulk concentrations representing different degrees of subsaturation. The amplitudes of the autocorrelation function are governed by the scattering intensities and the decay times are governed by the sizes of scatterers. The data clearly show that the sizes of mesoscale clusters increase with increasing concentration. Samples were prepared using method A. And data were collected in 20 mM Tris, pH 7.4, 100 mM KCl, at ≈ 25˚C.

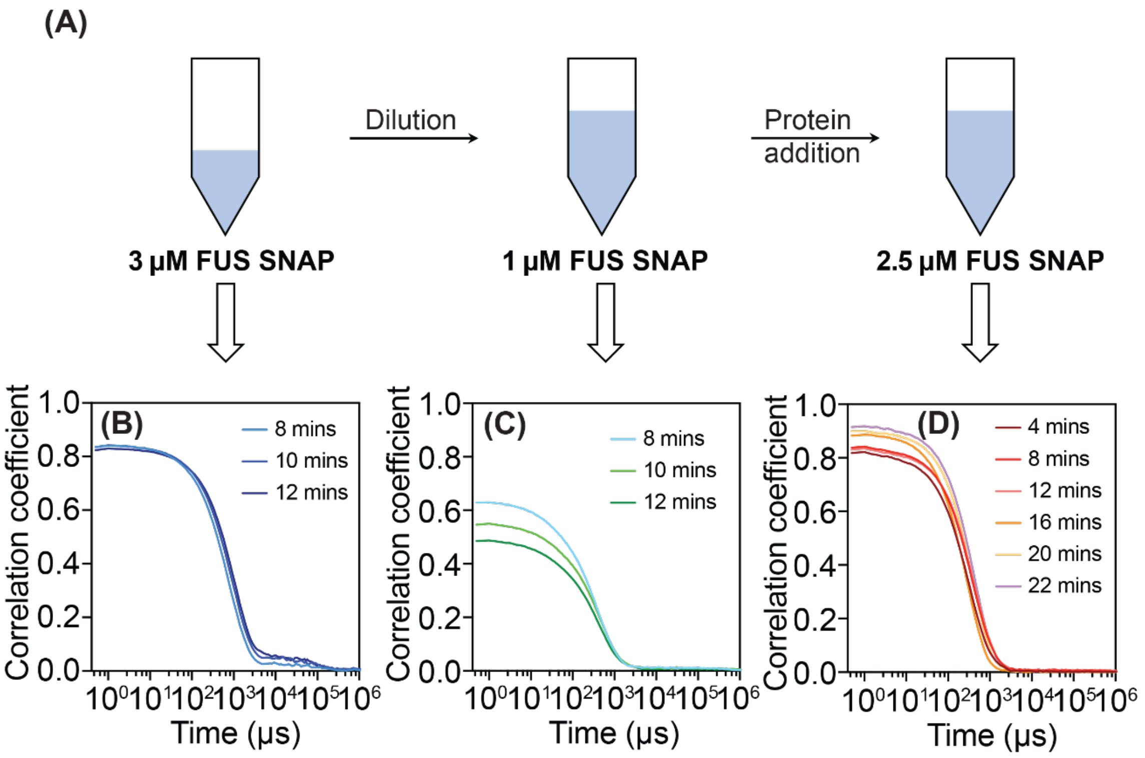


**Fig. S13: Establishing for the reversibility of cluster formation.** (A) Schematic summarizing the protocol used to dilute and concentrate FUS-SNAP samples (prepared using method A) starting from a reference bulk concentration of 3 μM. (B) Autocorrelation functions collected using DLS for a 3 μM solution of FUS-SNAP. This sets the reference amplitude maximum of the and decay time for the autocorrelation function. (C) Dilution by a factor of two causes a diminution in the maximum and shortens the decay time of the autocorrelation function. This indicates that dilution decreases the average sizes of clusters. (D) Concentrating the solution by a factor of 1.66 leads to autocorrelation functions with a larger amplitude and longer decay times. This is indicative of the average cluster sizes increasing as the concentrations increase.

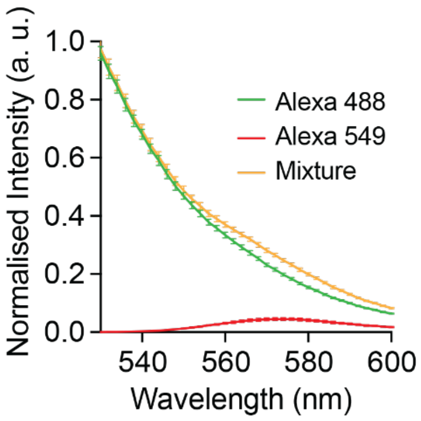


**Fig. S14: Control experiments performed using free dyes.** These experiments indicate that the observations of FRET signals for FUS molecules arise because of the exchange of labeled FUS molecules and not because of the dyes alone.

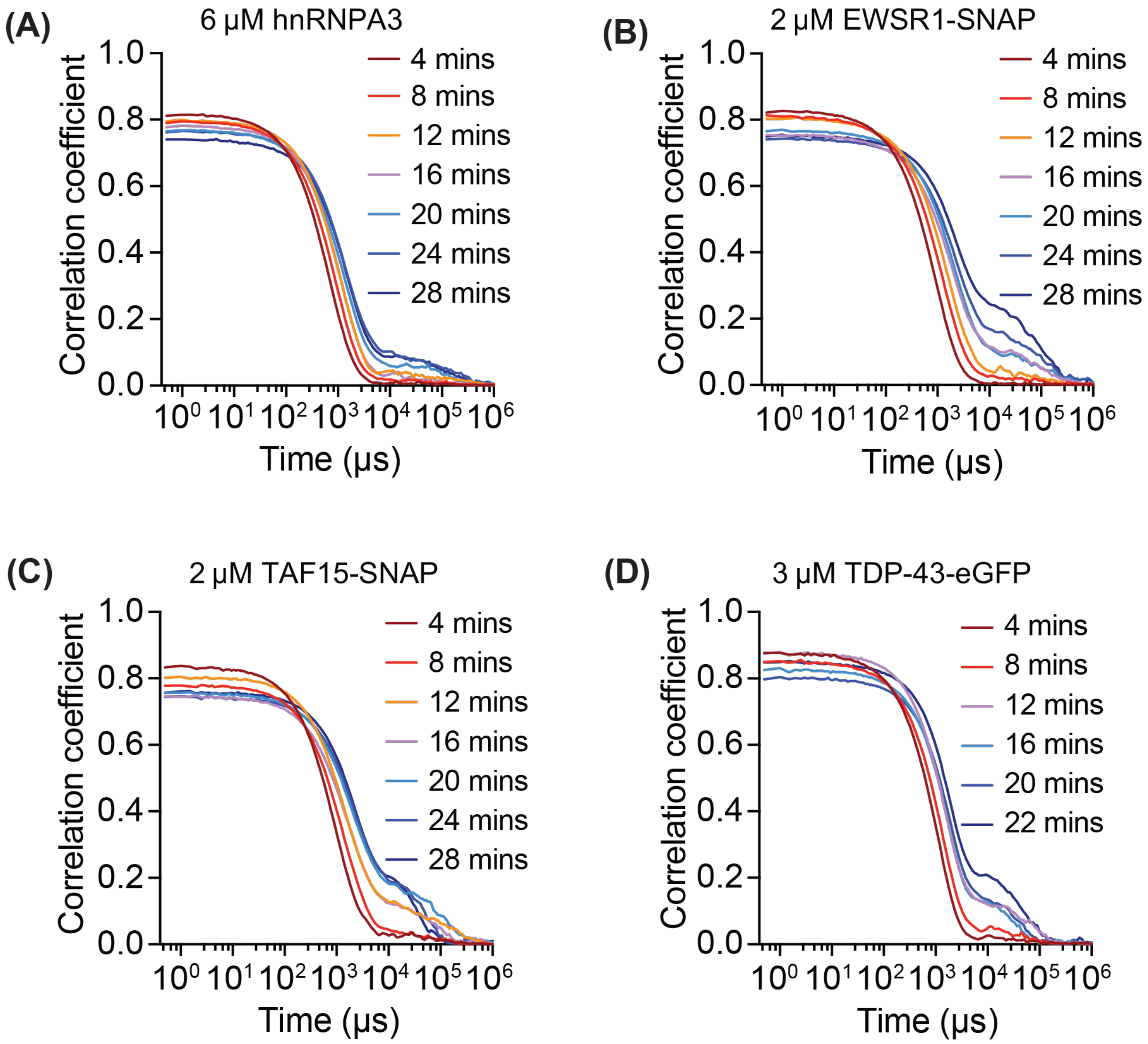


**Fig. S15: Autocorrelation functions from DLS measurements collected at the protein-specific $c_{sat}$ values for (A) hnRNP-A3, (B) EWSR1-SNAP, (C) TAF15-SNAP, and (D) TDP-43-eGFP.** These data show the presence of slow modes in the autocorrelation function profiles. These slow modes are only manifest at or above $c_{sat}$ and are absent below $c_{sat}$. And data were collected in 20 mM Tris, pH 7.4, 100 mM KCl, at ≈ 25˚C.

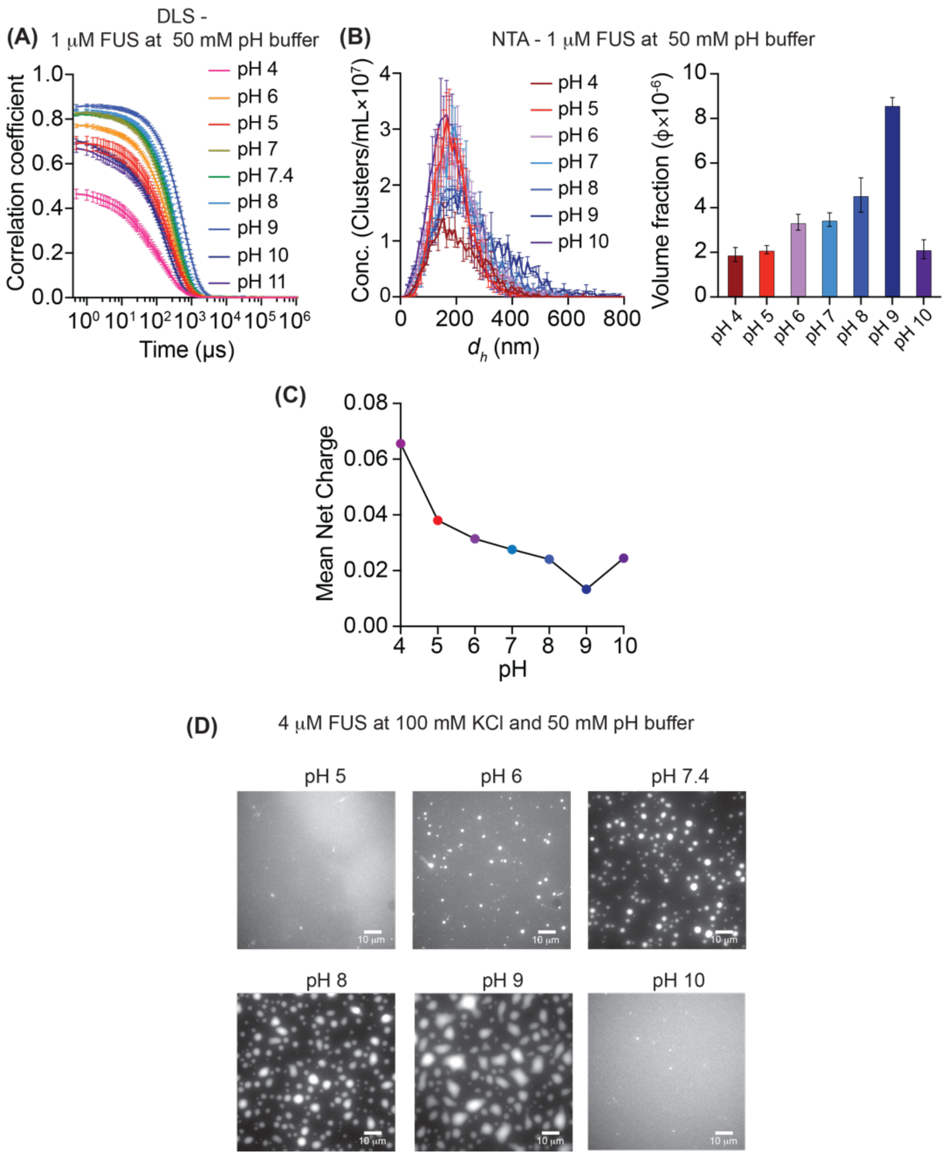


**Fig. S16: Impact of pH and linked changes to net charge on cluster formation and macroscopic phase separation of untagged FUS.** (A) Autocorrelation functions measured using DLS at the 8-minute time point for 1 μM untagged FUS. A series of traces are shown for solutions buffered to be at different pH values. The maximal amplitude and slowest decay are obtained for a pH of 9.0. (B) NTA data shown as distributions and as (C) abundance quantifying the distributions of mesoscale clusters at different pH values. (D) Estimates of the mean net charge of untagged FUS as a function of pH. These calculations made using unmodified $pK_a$ values for ionizable residues suggests that increasing the net charge weakens cluster formation. (E) Weakening cluster formation by increasing the mean net charge also weakens macroscopic phase separation. This is made visually evident using microscopy images, which show how condensates decrease / increase in size as the net charge increases / decreases. The scalebar is 10 μm and in the microscopy experiments 5% of the molecules are FUS-eGFP and the remaining 95% are untagged FUS. Samples were prepared using method A.

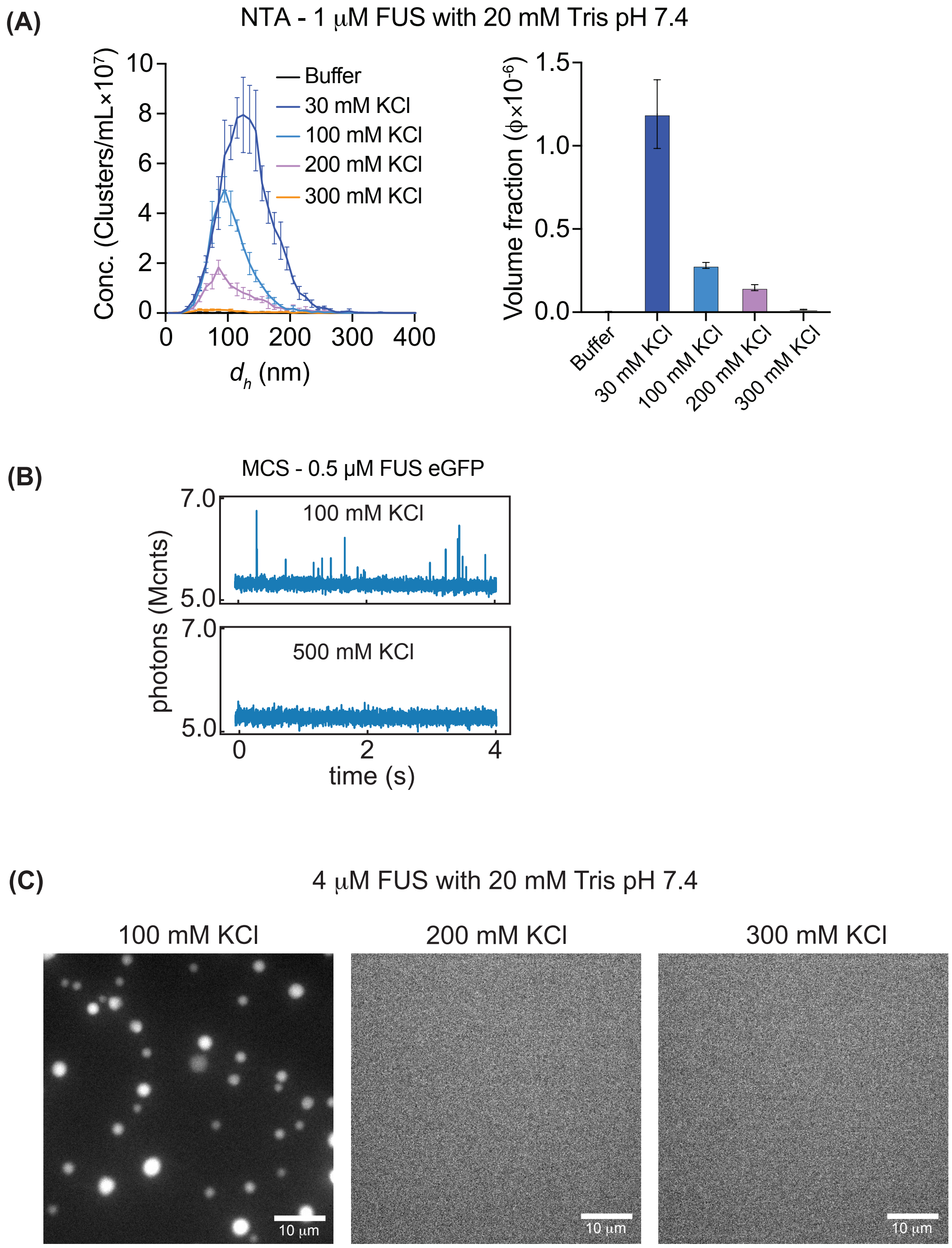


**Fig. S17: Increased concentration of KCl weakens cluster formation and macroscopic phase separation.** (A) The distribution of cluster sizes, measured using NTA, shifts toward smaller values as the concentration of KCl is increased. Data are shown here for 1 μM untagged FUS prepared using method A. (B) The abundance of mesoscale clusters formed by 1 μM untagged FUS decreases with increased concentration of KCl. (C) Comparative MCS data show that fluorescence spikes present at 100 mM KCl for 0.5 μM FUS-eGFP are lost in the presence of 500 mM KCl. (C) Loss of mesoscale clusters with increased KCl concentration causes a loss of macroscopic phase separation. This is shown using microscopy data collected at a concentration of 4 μM of untagged FUS. For a fixed protein concentration, increasing the concentration of KCl leads to a loss of condensates.

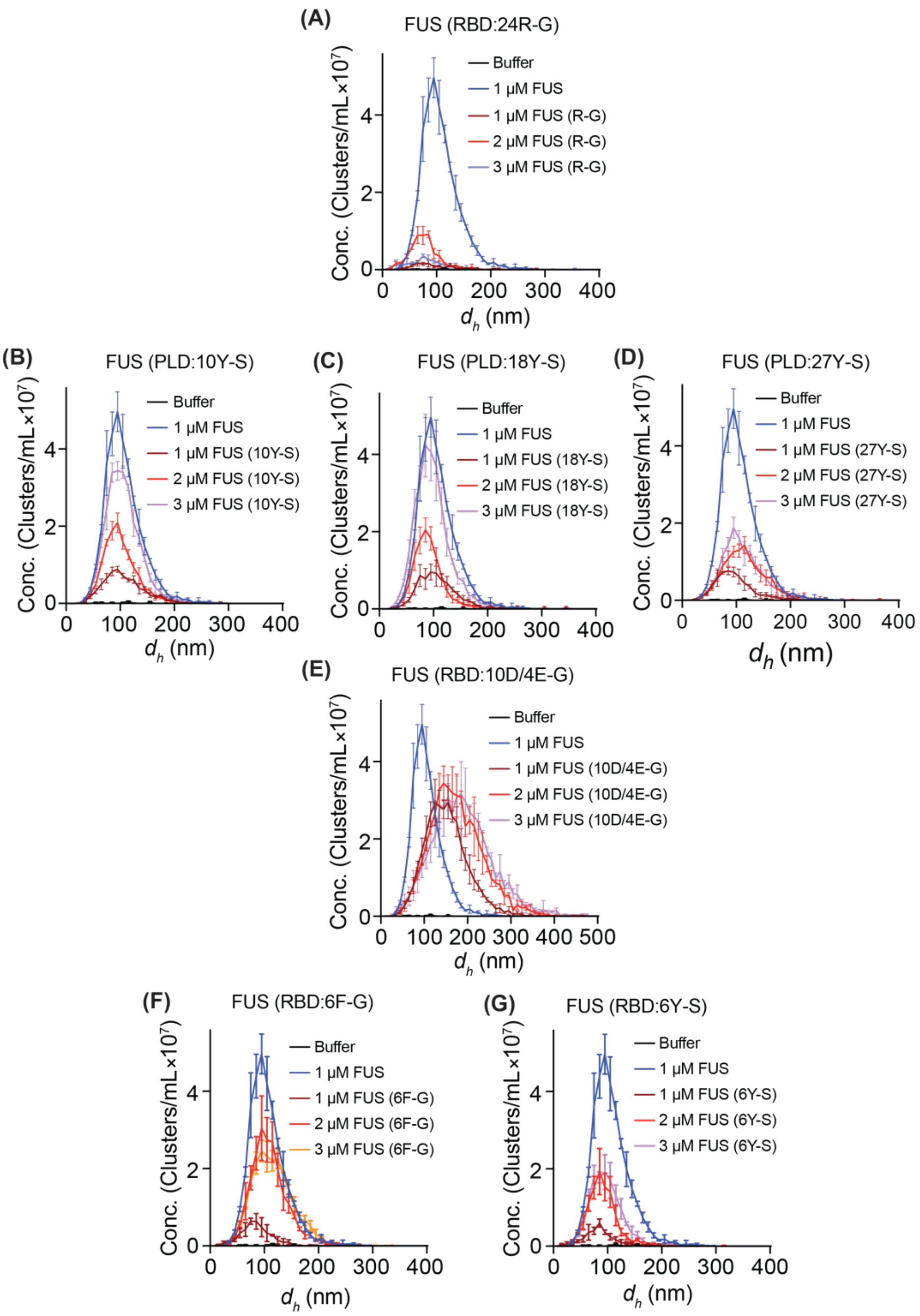


**Fig. S18: Impact of mutations within the PLD and RBD on the concentration dependence of size distributions of mesoscale clusters formed by untagged FUS.** Data are shown from NTA measurements. (A) Substitution of 24 Arg residues in the RBD of full-length FUS with Gly weakens cluster formation in subsaturated solutions. (B) Substitution of 10 Tyr residues with Ser in the PLD of full-length FUS shifts the cluster distributions toward smaller sizes. (C) Substitution of 18 Tyr residues with Ser in the PLD of full-length FUS shifts the cluster distributions toward smaller sizes. (D) Substitution of 27 Tyr residues with Ser in the PLD of full-length FUS shifts the cluster distributions toward smaller sizes. Taken together, the data in panels (B)-(D) show a valence-dependent response, whereby decreasing the number of Tyr residues in the PLD of full-length FUS has a concomitant decrease in the sizes of clusters that are formed. The chemistry-specific nature of the driving forces for cluster formation in subsaturated solutions is revealed using differences in distributions of mesoscale cluster sizes formed upon replacing six Phe residues in the RBD of FUS with Gly – panel (E) – vs. replacing six Tyr residues in the RBD of FUS with Ser – panel (F). And data were collected in 20 mM Tris, pH 7.4, 100 mM KCl, at ≈ 25˚C.

FUS (RBD, 24R-G)

**(A)**

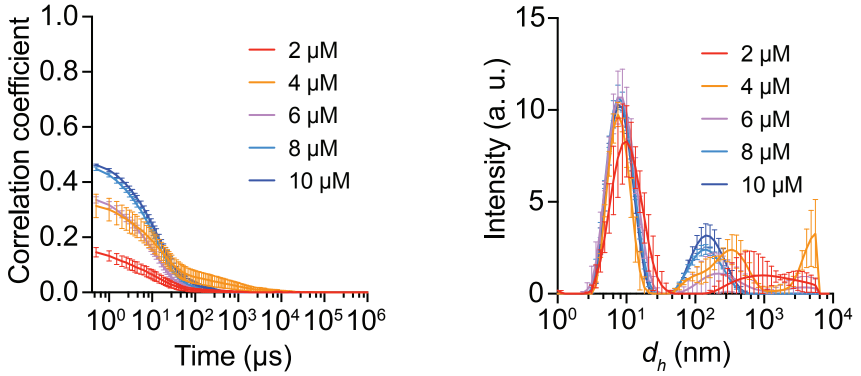


**Fig. S19: Impact of substituting 24 Arg residues within the RBD of FUS to Gly.** The data shown here are raw autocorrelation functions, gathered at different bulk concentrations of FUS (24R-G). These data demonstrate the reduced amplitude of the autocorrelation function (left), and rapid decay when compared to the wild-type FUS. The autocorrelation functions are converted to intensity profiles (right) to show that smaller species are formed when the Arg residues within the RBD are replaced by Gly. And data were collected in 20 mM Tris, pH 7.4, 100 mM KCl, at ≈ 25˚C.

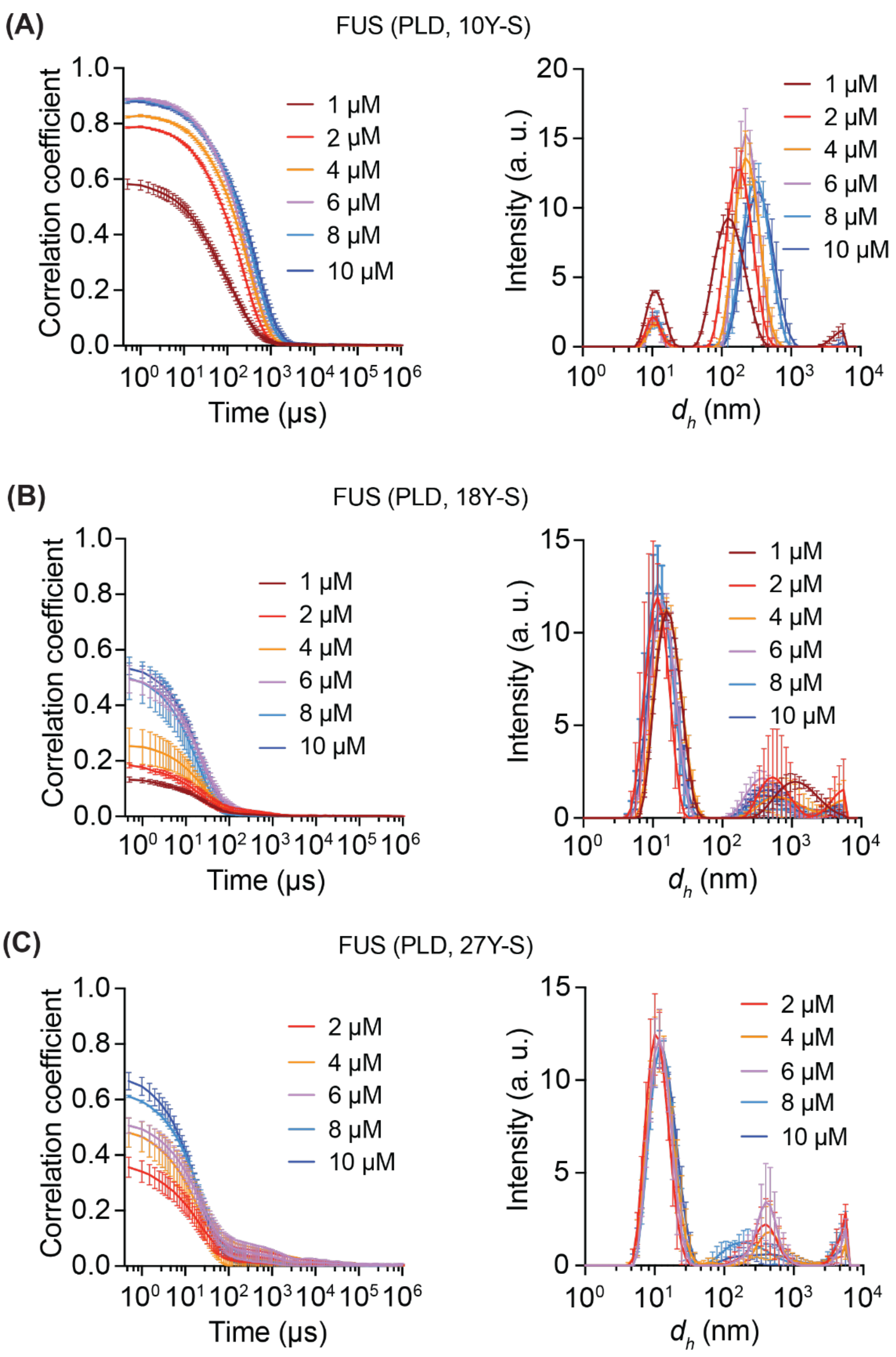


**Fig. S20: DLS data complementing the NTA distributions shown in Fig. S19.** These data show the impact of valence-dependent substitutions of Tyr residues in the PLD to Ser. (A) FUS (10Y-S), (B) FUS (18Y-S) and (C) (27Y-S). These data demonstrate the amplitude of the autocorrelation function (left), and the autocorrelation functions are converted to intensity profiles (right). And data were collected in 20 mM Tris, pH 7.4, 100 mM KCl, at ≈ 25˚C.

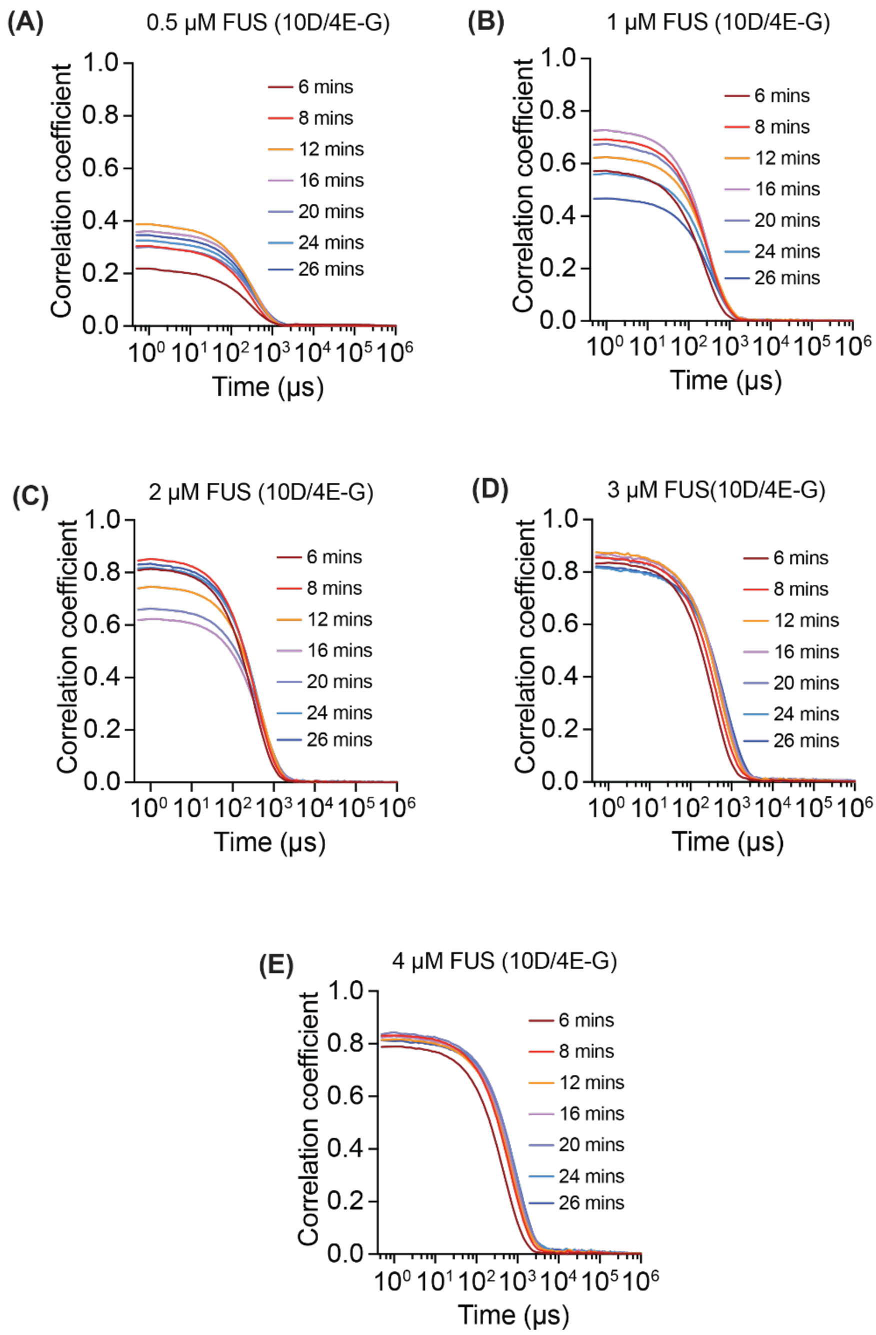


**Fig. S21: Impact of substituting 10 Asp / 4 Glu residues to Gly within the RBD of FUS on the DLS measured autocorrelation functions at different concentrations corresponding to subsaturated solutions.** When compared to the data from Fig. S2, the data shown in panels (A) – (D) suggest that the replacements of 10 Asp / 4 Glu residues to Gly within the RBD of FUS has a minimal impact of the extent of cluster formation in subsaturated solutions. However, panel (E) shows that the mutations weaken macroscopic phase separation as evidenced by the lack of slow modes being present above the $c_{sat}$ values that were quantified for wild-type FUS. These data show that mutations can maintain cluster formation while weakening macroscopic phase separation. And data were collected in 20 mM Tris, pH 7.4, 100 mM KCl, at ≈ 25˚C.

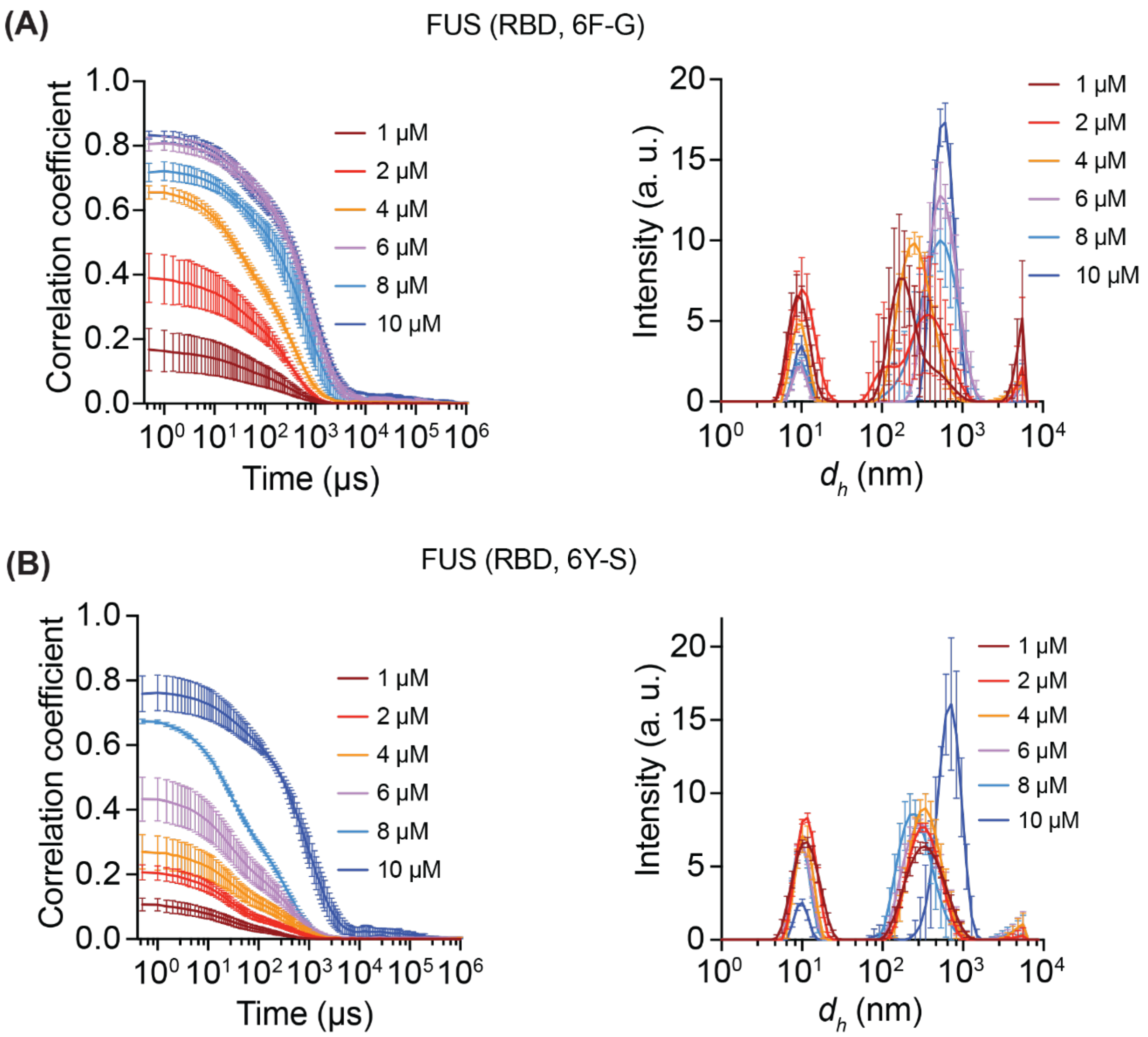


**Fig. S22: DLS data showing the chemistry-specific effects of Phe to Gly (A) vs. Tyr to Ser (B) substitutions on the formation of mesoscale clusters in subsaturated solutions.** These data demonstrate the amplitude of the autocorrelation function (left), and the autocorrelation functions are converted to intensity profiles (right). And data were collected in 20 mM Tris, pH 7.4, 100 mM KCl, at ≈ 25˚C. And data were collected in 20 mM Tris, pH 7.4, 100 mM KCl, at ≈ 25˚C.

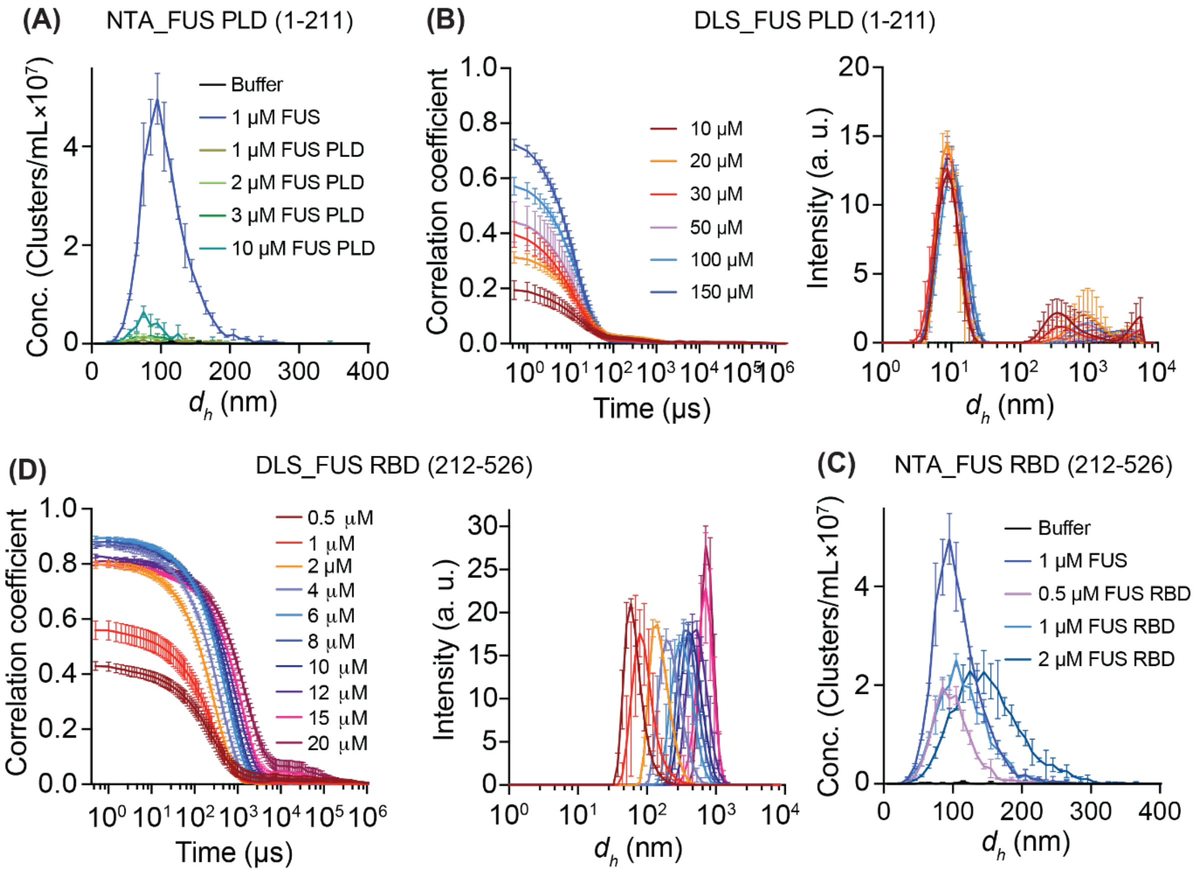


**Fig. S23: Comparisons of cluster formation and macroscopic phase separation for the PLD vs. RBD of FUS.** (A) NTA data collected at a series of concentrations for the PLD compared to the data collected for untagged, full-length FUS at 1 μM. These data show that the PLD alone is a weak driver of mesoscale clusters. (B) The NTA data are confirmed using DLS, as shown using raw autocorrelation functions collected for concentrations of the PLD that range from 10 μM to 150 μM. (C) The autocorrelation functions, when converted to intensity profiles, highlight the dominance of smaller species for the PLD. (D) The autocorrelation functions and accompanying intensity profiles – panel (E) – obtained for the RBD of FUS stand in contrast to those for the PLD. The autocorrelation functions show the onset of slow modes at an RBD concentration of 20 μM. This implies the onset of macroscopic phase separation above a $c_{sat}$ of 20 μM, which is an order of magnitude lower than the $c_{sat}$ for the PLD at similar solution conditions. (F) NTA data show the robust presence of mesoscale clusters forming in subsaturated solutions of the RBD. And data were collected in 20 mM Tris, pH 7.4, 100 mM KCl, at ≈ 25˚C.

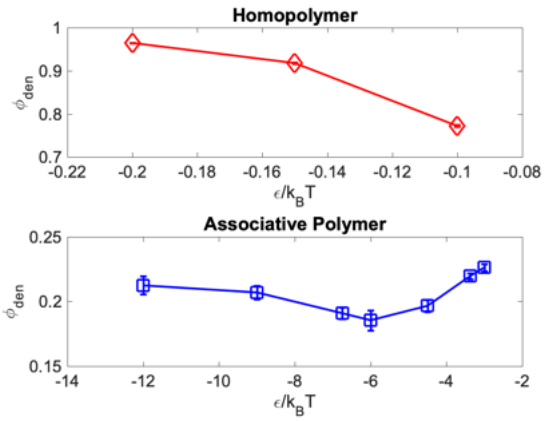


**Fig. S24: The volume fractions of polymers in dense phases formed for homopolymers vs. associative polymers.** Dense phases formed by homopolymers with a single energy scale are akin to polymer melts, $\phi_{den} \approx 1$. The dense phases formed by associative polymers are diluted compared to the homopolymers. This is because of the effects of spacers and the arrangement of stickers into crosslinked networks. Note that the energy scales for observing phase separation are renormalized by the stereospecific interactions among stickers.
